# Multiomics analysis reveals *B.* MO1 as a distinct *Babesia* species and provides insights into its evolution and virulence

**DOI:** 10.1101/2024.01.17.575932

**Authors:** Pallavi Singh, Pratap Vydyam, Tiffany Fang, Karel Estrada, Luis Miguel Gonzalez, Ricardo Grande, Madelyn Kumar, Sakshar Chakravarty, Vincent Berry, Vincent Ranwez, Bernard Carcy, Delphine Depoix, Sergio Sánchez, Emmanuel Cornillot, Steven Abel, Loic Ciampossin, Todd Lenz, Omar Harb, Alejandro Sanchez-Flores, Estrella Montero, Karine G. Le Roch, Stefano Lonardi, Choukri Ben Mamoun

## Abstract

Babesiosis, caused by protozoan parasites of the genus *Babesia*, is an emerging tick-borne disease of significance for both human and animal health. *Babesia* parasites infect erythrocytes of vertebrate hosts where they develop and multiply rapidly to cause the pathological symptoms associated with the disease. The identification of various *Babesia* species underscores the ongoing risk of new zoonotic pathogens capable of infecting humans, a concern amplified by anthropogenic activities and environmental shifts impacting the distribution and transmission dynamics of parasites, their vectors, and reservoir hosts. One such species, *Babesia* MO1, previously implicated in severe cases of human babesiosis in the midwestern United States, was initially considered closely related to *B. divergens*, the predominant agent of human babesiosis in Europe. Yet, uncertainties persist regarding whether these pathogens represent distinct variants of the same species or are entirely separate species. We show that although both *B.* MO1 and *B. divergens* share similar genome sizes, comprising three nuclear chromosomes, one linear mitochondrial chromosome, and one circular apicoplast chromosome, major differences exist in terms of genomic sequence divergence, gene functions, transcription profiles, replication rates and susceptibility to antiparasitic drugs. Furthermore, both pathogens have evolved distinct classes of multigene families, crucial for their pathogenicity and adaptation to specific mammalian hosts. Leveraging genomic information for *B.* MO1, *B. divergens*, and other members of the Babesiidae family within Apicomplexa provides valuable insights into the evolution, diversity, and virulence of these parasites. This knowledge serves as a critical tool in preemptively addressing the emergence and rapid transmission of more virulent strains.

## Introduction

Recent years have witnessed a significant rise in the number of tick-borne disease cases reported worldwide and an increase in the populations of ticks as well as medically important pathogens transmitted by these vectors [1]. In the United States, tick-borne diseases accounted for more than 75% of all vector-borne infections reported between 2004 and 2016 [2]. This threat to public health is expected to worsen with the continued changes in the natural environment, expansion of the geographic distribution of ticks and their reservoir hosts, rapid growth of the human population, and land use changes [3]. Several tick-borne bacterial, viral, and protozoan pathogens are known to cause infection in humans. Among these are *Babesia* pathogens, which infect human erythrocytes and cause human babesiosis, an emerging malaria-like illness with disease outcomes ranging from mild to severe or even fatal depending on the *Babesia* species, and the age and immune status of the infected individual [4].

*Babesia* species are protozoan parasites belonging to the order Piroplasmida and the phylum Apicomplexa. They are closely related to *Plasmodium*, *Toxoplasma* and *Theileria*, the agents of human malaria, toxoplasmosis, and theileriosis, respectively [4]. *Babesia* parasites have been found in vertebrate hosts throughout the world with some species capable of infecting multiple mammals, whereas others are host specific. Most cases of human babesiosis in Europe are caused by *Babesia divergens*, predominantly among asplenic patients [5]. These infections are accompanied by high parasite burden and are often fatal. Cases of babesiosis in individuals with intact spleens have also been reported [6–9]. *B. divergens* also infects cattle causing “red water fever” [10]. Other human babesiosis cases in Europe have been attributed to *B. venatorum* and *B. microti* [5, 11]. In the United States, cases of human babesiosis have so far been linked to at least three *Babesia* species: *Babesia microti*, which accounts for most cases reported annually; *B. duncani*, which was linked to severe babesiosis cases in Washington and California; and a *B. divergens*-like species (MO-1) reported in Missouri and Kentucky [12–14]. A previous report by Hollman and colleagues identified a parasite (NR831) that shares 99.8% sequence identity at the small subunit ribosomal RNA gene (SSU rRNA) with the MO-1 isolate [15]. The parasite was isolated from eastern cottontail rabbits (*Sylvilagus floridanus*) and *Ixodes dentatus* ticks on Nantucket Island, Massachusetts [15]. However, unlike *B. divergens*, the isolate failed to cause infection in Holstein-Friesian calves, and inoculated animals remained fully susceptible upon challenge inoculation with *B. divergens* [16].

Recently, the genome sequences of two *B*. *divergens* isolates, 1802A and Rouen 87, have been reported [17, 18]. The genome of the *B. divergens* 1802A strain, isolated from cattle, was reported to be 9.58 Mb in size and to encode 4,134 genes [18]. The genome sequence of the human reference strain, *B. divergens* Rouen 87, was reported by two separate research groups with one group reporting a genome size of 8.97 Mb encoding 4.097 genes [18], and the another reporting a genome size of 10.7-Mb encoding more than 3,741 genes [17]. This latest *B. divergens* Rouen 87 genome assembly was further improved by exploiting the previous sequence data using new computational tools and assembly strategies [19], with an updated size of 9.73 Mb encoding 4,546 genes [19]. Transcriptional data and gene expression profiling of *B. divergens* Rouen 87 free extra-cellular merozoites and intraerythrocytic parasites further provided new insights into the molecular mechanisms of invasion, gliding motility, and egress of this parasite [19]. Subsequent analyses using single-cell RNA sequencing enable construction of a pseudo-time-course trajectory of the parasite’s gene expression profiles during its intraerythrocytic life cycle, pinpointing differentially-expressed genes characteristic of each phase [20]. Unlike *B. divergens*, the biology, diversity, and virulence of *B.* MO1 remain completely unknown as does the relationship between these pathogens.

Here we report the first and complete sequence, assembly, and annotation of *B.* MO1, its transcription during its asexual development within human red blood cells. These genes are likely crucial for the parasite adaptation to the mammalian host. We further completed its epigenetic profile and genome 3D structure at 10 kb resolution, and demonstrate that these parasites express unique, complex and most likely evolving multigene families that interact with each other in a large heterochromatin cluster; reminiscence of the genome organization of genes involved in antigenic variation, observed in several *Plasmodium* species [21–23] as well as *Babesia* [23, 24] and Trypanosoma [25]. A comprehensive analysis of the genomic data, along with cell biological investigations, offers substantial evidence in favor of designating *B. MO1* as a separate species within the Babesiidae family.

## Results

### *B.* MO1 and *B. divergens* exhibit distinct replication rates during their intraerythrocytic life cycles

Available epidemiological studies suggest that the cottontail rabbit, *Sylvilagus floridanus*, serves as an animal reservoir for *B.* MO1, transmission to large mammals, including humans, is facilitated by *Ixodes dentatus* ticks (**Fig 1A**) [15]. In contrast, its close relative *B. divergens*, which infects cattle and humans, is transmitted by *I. ricinus* (**Fig 1A**) [26]. A previous study has shown successful continuous propagation of *B.* MO1 isolated from eastern cottontail rabbits (*Sylvilagus floridanus*) in human red blood cells using HL-1 growth medium supplemented with 20% human serum [15]. In this study, *B.* MO1 was cultured in vitro either in DMEM/F12 or RPMI media. Microscopic examination of *B.* MO1 cultures revealed an asynchronous replication rate, with daughter parasites dividing independently, resulting in a single infectious ring stage parasite generating 2, 3, 4, 5, 6, 7, and ultimately 8 daughter parasites (merozoites) (**Fig 1B**). Some red blood cells can host more than one parasite (multiple infections), which divide independently of each other leading to the formation of multiple stages within the same infected cell (**Fig 1B**). While multiple stages are also seen in *B. divergens*-infected human red blood cells, this parasite produces only four daughter parasites from each invading merozoite (**Fig 1C**). Since the parental *B.* MO1 was initially isolated from cottontail rabbits for in vitro propagation, it represents a heterogeneous population. Therefore, we opted to clone *B.* MO1 as well as *B. divergens* Rouen 87 to obtain pure clonal lines for continuous in vitro culture. These clonal lines were used in all experiments conducted in this study and could be propagated continuously in human erythrocytes in RPMI medium supplemented with either 20% fetal bovine serum (FBS) or Albumax (**Fig 1D**). Notably, there was a marked disparity in the growth rates between *B.* MO1 and *B. divergens*, as evidenced by the doubling of parasitemia levels every 42 to 48 hours for *B.* MO1 clones and every 16 to 18 hours for *B. divergens* clones (**Fig 1D**).

**Figure 1:**
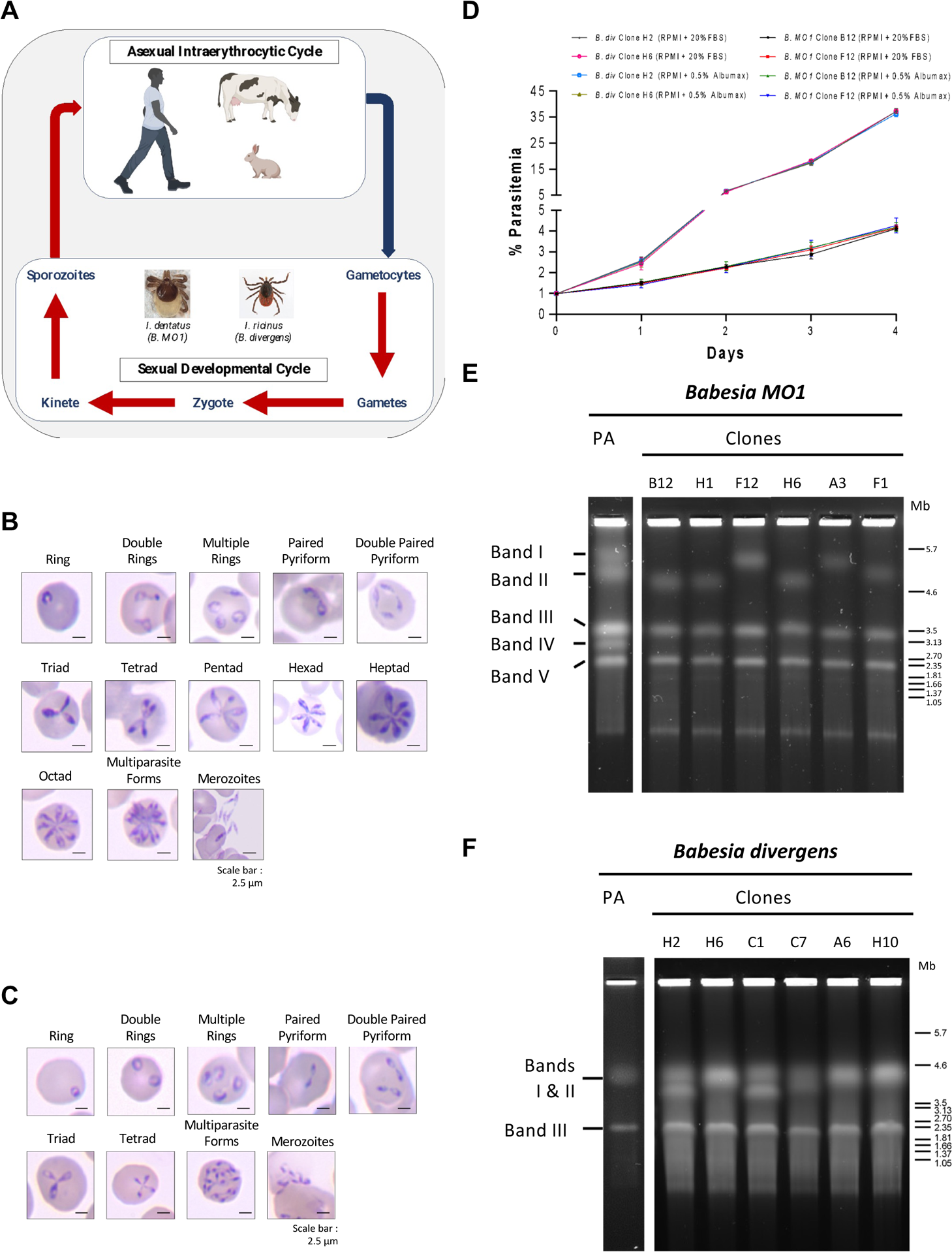
Life cycle of *B.* MO1 and *B. divergens*. **A.** Schematic representation of the life cycle of *B.* MO1 and *B. divergens* in vertebrate hosts (humans, cattle, cottontail rabbit) and tick vectors. **B.** Representative Giemsa-stained light microscopic images of the various stages of *B.* MO1 propagated in human erythrocytes in vitro. **C.** Representative Giemsa-stained light microscopic images of the various forms of *B. divergens* Rouen 87 grown in human erythrocytes in vitro. **D.** Growth of *B. divergens* Rouen 87 clones H2 and H6, and *B.* MO1 clones B12 and F12 in human RBCs in RPMI medium supplemented with either 20% FBS or 0.5% albumax over a course of 4 days. Two independent experiments were performed in triplicates. **E**. Chromosomal organization of *Babesia* MO1. PFGE shows the number and approximate sizes of bands in *B.* MO1 parental (PA) isolate: ∼5.7 Mb, ∼4.6 Mb, ∼3.5 Mb, ∼3.13 Mb and ∼2.35 Mb; the number and approximate sizes of bands in *B.* MO1 clones B12, H1, H6, and F1: ∼4.6 Mb (Chromosome I) ∼3.5 Mb (Chromosome II), and ∼2.35 Mb (Chromosome III) and *B.* MO1 clones F12 and A3: ∼5.7 Mb (Chromosome I), ∼3.5 Mb (Chromosome II), and ∼2.35 Mb (Chromosome III). The experiment was performed in biological duplicates. **F.** Chromosomal organization of *Babesia. divergens*. PFGE shows the number and approximate sizes of bands in *B. divergens* Rouen 87 parent, clones H6, A6, and H10: ∼4.3 Mb (Chromosome I and Chromosome II), and ∼2.1 Mb (Chromosome III) and *B. divergens* clones H2, C1 and C7: ∼4.3 Mb (Chromosoe I), ∼4.1 Mb (Chromosome II), and ∼2.1 Mb (Chromosome III). *Hansenula wingei* and *Schizosaccharomyces pombe* DNA chromosomes were used as DNA markers. The manufacturer’s estimate of the sizes of chromosomes are indicated in Megabase pairs [13] on the right-hand side of panels E and F. The experiment was performed in biological duplicates.

### Chromosomal organization of the nuclear genomes *B.* MO1 and *B. divergens* Rouen 87

The chromosomal arrangement of both the parent and clones of *B.* MO1 nuclear genomes was confirmed through pulse field gel electrophoresis (PFGE) analysis. PFGE on the *B.* MO1 parent displayed five bands with approximate sizes of ∼5.7 Mb, ∼4.6 Mb, ∼3.5 Mb, ∼3.13 Mb, and ∼2.35 Mb (**Fig 1E**). Interestingly, after dilution cloning of the parental isolate, all the clones obtained contained only three bands as indicated by PFGE analysis. *B.* MO1 clones F12 and A3 exhibited three bands with approximate sizes of ∼5.7 Mb (Chromosome I), ∼3.5 Mb (Chromosome II), and ∼2.35 Mb (Chromosome III). Additionally, *B.* MO1 clones B12, H1, H6, and F1 also manifested three bands, one of which had an approximate size of ∼4.6 Mb (Chromosome I), while the other two matched the sizes of bands (Chromosome II and III) observed in clones F12 and A3. These findings suggest that the parent *B.* MO1 strain comprises a mixture of different clones, each containing three chromosomes, likely undergoing significant recombination during the intraerythrocytic life cycle of the parasite (**Fig 1E**). The chromosomal profile of *B. divergens* Rouen 87 parent and clones H6, A6 and H10 revealed three bands in PFGE. Two of these bands overlapped, with approximate sizes of ∼4.3 Mb, encompassing Chromosomes I and II, while the other band measured ∼2.1 Mb (Chromosome III). Similarly, *B. divergens* Rouen 87 clones H2 and C1 displayed three band sizes of ∼4.3 Mb (Chromosome I), ∼4.1 Mb (Chromosome II), and ∼2.1 Mb (Chromosome III) (**Fig 1F**). The chromosomal profile of *B.* MO1 differed from that of several *B. divergens* clinical isolates from France and Spain **(Fig S1**). These *B. divergens* isolates exhibited three distinct chromosomes, with sizes varying between isolates, as confirmed by PFGE and Southern blot assays (**Fig S1**).

### Sequencing, genome assembly, genome annotation and assembly quality control

To gain deeper insights into the biology of *B.* MO1, genomic DNA from clone F12 and clone B12 was sequenced using PacBio HiFi technology. For clone F12, approximately 2.7 million PacBio HiFi reads were generated, with an average read length of approximately 11.5 Kb and the longest read extending to approximately 49.4 Kb. These HiFi reads accounted for approximately total yield of 31.2 billion bases, providing an expected coverage of approximately 2,600x for the *B.* MO1 genome (assuming a 12 Mb genome). The assembly of clone F12 genome (details in supplemental methods) utilized the longest HiFi reads, and its quality was independently validated using the Bionano optical map for parental *B.* MO1. The optical map consisted of eight optical molecules (**Table S1**) with a total of 11.4 million bases. The alignment of the clone F12 assembly against the optical map (**Fig S2A)** showed that except for the 5’ and 3’ ends of Chromosome I and the 3’ end of Chromosome II, there is a strong agreement between the assembly and the optical map. No indication of chimeric contigs (i.e., mis-joins) could be detected.

Our assembly revealed a deficiency in covering the telomeric ends of Optical Molecule 1, with approximately 0.7 Mb missing from the 5’ end and about 0.5 Mb from the 3’ end. Molecule 2, on the other hand, has comprehensive coverage through the assembly of Chromosome II and an additional contig. Optical Molecule 3 lacks approximately 0.1 Mb at the 3’ end. While the overall assembly quality for all chromosomes is high, it is apparent that some telomeres may not have fully assembled due to their repetitive nature. In our investigation of the clone F12 assembly’s terminal regions using RepeatMasker [27], we identified interstitial telomeric repeat sequences. Notably, an ∼11 Kb ITS was found at the 5’ end of Chromosome II, a ∼7 Kb ITS at the 3’ end of Chromosome II, and a ∼5 Kb ITS at the 5’ end of Chromosome III (**Fig S2A**). The combined size of the eleven unplaced contigs is approximately ∼965 Kb, and none of these contigs contains an ITS. For the B12 clone, we obtained around ∼2.8 million PacBio HiFi reads, with an average read length of ∼11.9 Kb and the longest read measuring ∼50.1 Kb. The HiFi reads totaled ∼33.8 billion bases, translating to an anticipated ∼2,800x coverage of the *B.* MO1 genome (assuming a 12 Mb genome). The B12 genome assembly, using the longest HiFi reads (details in Supplemental Methods), was aligned against the optical map, as illustrated (**Fig S2B**). A strong agreement between the clone B12 assembly and the optical map for the parental *B.* MO1 was established, with the notable exceptions of the 5’ and 3’ ends of Chromosome I and the 3’ end of Chromosome II. Upon reevaluation of the terminal regions of B12 assembly using RepeatMasker, we identified an ∼9 Kb ITS at the 5’ end of Chromosome II, an ∼8 Kb ITS at the 3’ end of Chromosome II, a ∼7 Kb ITS at the 5’ end of Chromosome III, and a ∼9 Kb ITS at the 3’ end of Chromosome III (**Fig S2B**). The cumulative size of the nine unplaced contigs is ∼1071 Kb, and similar to the F12 assembly, none of these contigs contains an ITS. As shown in **Fig S3**, which displays a synteny plot generated using SyRI [28] and plotSR [29], a robust syntenic agreement is evident among the assemblies of F12, M12, and the parental *B.* MO1. The primary distinctions include a 256 Kb insertion on Chromosome I in F12 compared to B12 and a ∼136 Kb insertion on Chromosome III in B12 compared to F12. Additional variations occur at the telomeres, likely attributed to incomplete assemblies of repetitive regions or recombination events occurring between chromosomes within repetitive regions leading to variation in chromosome size between clones.

For the *B. divergens* Rouen 1987 strain, we acquired approximately ∼186 thousand Oxford Nanopore (ONT) reads, with an average ONT read length of ∼5.4 Kbp and the longest read extending to 156 Kbp. The total ONT reads amounted to around 1 billion bases, resulting in an anticipated ∼100x coverage of the *B. divergens* genome. Using the Canu assembler [30], we carried out the genome assembly, which was later polished with Illumina reads (details in Supplemental Methods). For further examination of the *B. divergens* Rouen assembly, we conducted a third search for interstitial telomeric repeat sequences [17] using RepeatMasker. This analysis revealed a ∼2 Kb ITS at the 5’ end of Chromosome I, a ∼2 Kb ITS at the 3’ end of Chromosome I, a ∼2 Kb ITS at the 3’ end of Chromosome II, a ∼1 Kb ITS at the 5’ end of Chromosome III, and a ∼1 Kb ITS at the 3’ end of Chromosome III. Notably, the longest unplaced contig, spanning 219 Kb, exhibited ∼1 Kb ITS sequences at both ends, hinting at the possibility of a short, fourth chromosome. The cumulative size of the nine unplaced contigs is approximately ∼363 Kb. The key statistics of these new genome assemblies are summarized in **Table I**. Notably, the data indicates that the *B.* MO1 F12 and B12 assemblies, along with the Rouen assembly, share similarities in total length (11 Mb) (**Table II**). They also exhibit the same chromosome count, and comparable N50, GC content, and genome content completeness against the apicomplexa_odb10 database as assessed by BUSCO v5 [31] (**Table II**).

**Table I.** Genome comparison and gene statistics.

|  | *Babesia divergens*<br>(Rouen-87) | *Babesia* MO1<br>(F12 clone) | 2018 *B. divergens*<br>Rouen assembly |
| --- | --- | --- | --- |
| Total gene models /<br>annotated | 5,274 / 3,558 | 4,569 / 2,795 | 4,546 / 3,386 |
| Exon mean / median /<br>mode | 472.85 / 185 / 75 & 102 | 458.065 / 176 / 99 | 727 / 352 / 65 |
| Intron mean / median /<br>mode | 605.89 / 40 / 33 | 421.24 / 39 / 33 | 328 / 105 / 36 |
| Average exons per<br>gene | 2.65 | 3.14 | 1.68 |

**Table II.** Assembly statistics of *Babesia divergens* Rouen and *Babesia MO1*.

|  | *B. divergens*<br>(Rouen 87) | *Babesia* MO1<br>(clone F12) | *Babesia* MO1<br>(clone B12) | 2018 *B. divergens*<br>Rouen<br>(assembly<br>ASM107745v2) |
| --- | --- | --- | --- | --- |
| Total length (Mb) | 10.78 | 11.03 | 10.8 | 9.73 |
| Total chromosomes | 3 | 3 | 3 | Undetermined #<br>5 scaffolds > 1Mb |
| Unplaced contigs | 7 | 14 | 9 | 141 |
| Mean contig length<br>(Mb) | 1.07 | 0.787 | 0.899 | 0.069 |
| Longest contig (Mb) | 4.35 | 3.98 | 3.67 | 2.20 |
| N50/L50<br>(Mb/contigs) | 3.95 / 2 | 3.65 / 2 | 3.49 / 2 | 1.08 / 4 |
| GC content | 45 | 45 | 45 | 43 |
| BUSCO v5<br>(Apicomplexa<br>lineage) | 437/446 Complete<br>1/446 Fragmented<br>8/446 Missing | 434/446 Complete<br>2/446 Fragmented<br>10/446 Missing | 434/446 Complete<br>2/446 Fragmented<br>10/446 Missing | 437/446 Complete<br>1/446 Fragmented<br>8/446 Missing |

We further compared the new genome assemblies at the nucleotide level. A comparison between the clone F12 assembly against the clone B12 assembly (**Fig S4A)** showed a high level of sequence similarity in the non-telomeric regions of Chromosomes I-III, pronounced repetitive content at both ends of Chromosome I and III, while Chromosome II exhibited significant repetition primarily at the 3’ end, and notable repetitive content in the majority of unplaced contigs. A similar pattern was found when the assemblies of *B.* MO1 clone F12 and the parental strain were compared (**Fig S4B**). Interestingly, comparison of the assemblies between *B. divergens* Rouen 87 strain and the *B.* MO1 F12 clone revealed that Chromosome II in *B. divergens* Rouen corresponds to Chromosome I in *B.* MO1 F12, with sequence similarity breaking at the telomeres, Chromosome III in *B. divergens* Rouen corresponds to Chromosome III in *B.* MO1 F12, with sequence similarity breaking at the telomeres, and Chromosome I in *B. divergens* Rouen strain corresponds to Chromosome II in *B.* MO1 F12, with a notable ∼600 Kb insertion that appears to be highly repetitive in *B. divergens*.

Gene annotations for the *B.* MO1 F12 clone were conducted using FunAnnotate (https://github.com/nextgenusfs/funannotate) and PAP (https://github.com/kjestradag/PAP) pipelines. The gene annotations for *B. divergens* Rouen 87 strain were transferred to the improved assembly using the PATT (https://github.com/kjestradag/PATT) pipeline. The gene models for *B.* MO1 were established based on annotations from evolutionarily related species, and further refined using PacBio Iso-seq data specific to *B.* MO1 (refer to Methods for details). These analyses yielded 4569 gene models for *B.* MO1 clone F12 and 5,274 for *B. divergens* (**Table I**). The annotated genome of *B.* MO1 revealed that all the enzymes of the glycolytic pathway and tricarboxylic acid cycle are present in the genome (**Table III and IV**). Our analysis also identified 20 members of GPI-anchored proteins (**Table V**) and 21 members of Apicomplexan Apetala 2 (ApiAP2) family (**Table VI**).

**Table III.**
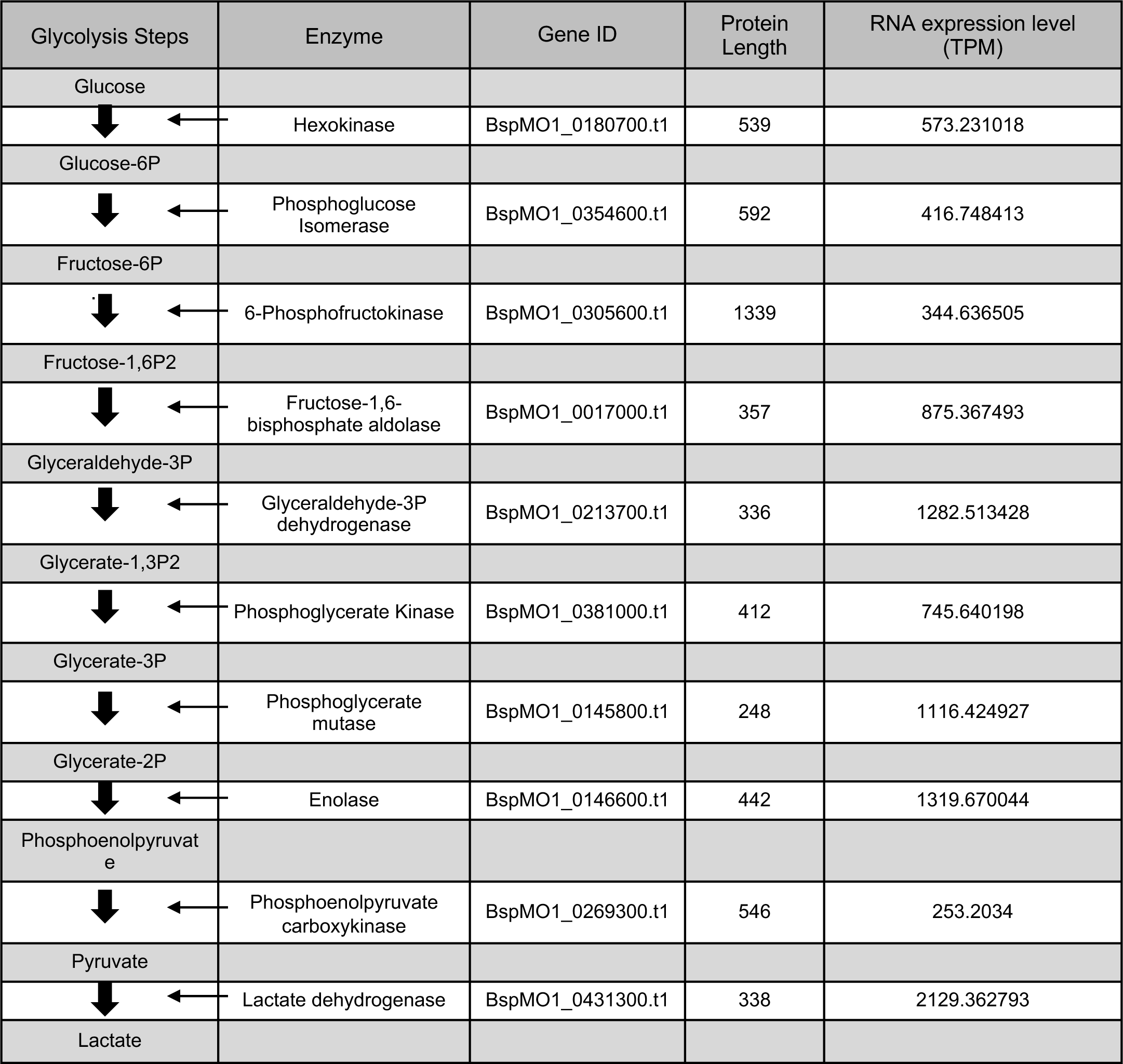
Predicted enzymes of the glycolytic pathway of *B.* MO1.

| Glycolysis Steps | Enzyme | Gene ID | Protein Length | RNA expression level (TPM) |
| --- | --- | --- | --- | --- |
| Glucose |  |  |  |  |
| ← | Hexokinase | BspMO1_0180700.t1 | 539 | 573.231018 |
| Glucose-6P |  |  |  |  |
| ← | Phosphoglucose Isomerase | BspMO1_0354600.t1 | 592 | 416.748413 |
| Fructose-6P |  |  |  |  |
| ← | 6-Phosphofructokinase | BspMO1_0305600.t1 | 1339 | 344.636505 |
| Fructose-1,6P2 |  |  |  |  |
| ← | Fructose-1,6-bisphosphate aldolase | BspMO1_0017000.t1 | 357 | 875.367493 |
| Glyceraldehyde-3P |  |  |  |  |
| ← | Glyceraldehyde-3P dehydrogenase | BspMO1_0213700.t1 | 336 | 1282.513428 |
| Glycerate-1,3P2 |  |  |  |  |
| ← | Phosphoglycerate Kinase | BspMO1_0381000.t1 | 412 | 745.640198 |
| Glycerate-3P |  |  |  |  |
| ← | Phosphoglycerate mutase | BspMO1_0145800.t1 | 248 | 1116.424927 |
| Glycerate-2P |  |  |  |  |
| ← | Enolase | BspMO1_0146600.t1 | 442 | 1319.670044 |
| Phosphoenolpyruvate |  |  |  |  |
| ← | Phosphoenolpyruvate carboxykinase | BspMO1_0269300.t1 | 546 | 253.2034 |
| Pyruvate |  |  |  |  |
| ← | Lactate dehydrogenase | BspMO1_0431300.t1 | 338 | 2129.362793 |
| Lactate |  |  |  |  |

**Table IV.** Predicted enzymes of the TCA cycle of *B.* MO1.

| Krebs Cycle steps | Enzyme | Predicted Gene ID | Protein Length | C/M/S | TM | RNA expression level (TPM) |
| --- | --- | --- | --- | --- | --- | --- |
| Citrate |  |  |  |  |  |  |
| ↓ | ← Aconitate Hydratase | BspMO1_0041000.t1 | 914 | M | - | 263.30365 |
| Isocitrate |  |  |  |  |  |  |
| ↓ | ← Isocitrate dehydrogenase | BspMO1_0098400.t1 /BspMO1_0139600.t1 | 519 / 455 | M | - | 102.51371 / 595.400269 |
| 2-oxaloglutarate |  |  |  |  |  |  |
| ↓ | ← 2-Oxoglutarate dehydrogenase | BspMO1_0276100.t1 | 952 | C | - | 251.601196 |
| Succinyl-CoA |  |  |  |  |  |  |
| ↓ | ← Succinyl-CoA synthetase | BspMO1_0145000.t1 /BspMO1_0407900.t1 | 461 | M | - | 324.873474 |
| Succinate |  |  |  |  |  |  |
| ↓ | ← Succinate dehydrogenase | BspMO1_0256000.t1 /BspMO1_0324700.t1 | 273 624 | M | - | 269.766235 / 139.289917 |
| Fumarate |  |  |  |  |  |  |
| ↓ | ← Fumarase | BspMO1_0281100.t1 | 468 | M | - | 156.482895 |
| Malate |  |  |  |  |  |  |
| ↓ | ← Malate dehydrogenase | BspMO1_0431300.t1 | 338 | M | - | 2129.362793 |
| Oxaloacetate |  |  |  |  |  |  |
| ↓ | ← Citrate synthase | BspMO1_0118500.t1 /BspMO1_0313600.t1 | 361 / 608 | C / M | - | 267.057251 / 214.3311 |

**Table V.** Predicted GPI-anchored proteins of *B.* MO1.

|  |  | Signal<br>P5.0 | PredGPI |  |  |  |
| --- | --- | --- | --- | --- | --- | --- |
| GPI-AP ID | Protein ID | Score | Specificity<br>Score | Probability | Protein<br>length (aa) | TPM Value |
| BMO1GPI1 | BspMO1_0016700.t1.1 | 0.9782 | 100 | Highy Probable | 459 | 102.585358 |
| BMO1GPI2 | BspMO1_0018700.t1.1 | 0.9763 | 100 | Highy Probable | 479 | 763.598877 |
| BMO1GPI2 | BspMO1_0001800.t1.1 | 0.9784 | 99.9 | Highy Probable | 134 | 10.512491 |
| BMO1GPI4 | BspMO1_0090300.t1.1 | 0.9243 | 100 | Highy Probable | 555 | 331.263947 |
| BMO1GPI5 | BspMO1_0096800.t1.1 | 0.6156 | 100 | Highy Probable | 619 | 403.405823 |
| BMO1GPI6 | BspMO1_0088800.t1.1 | 0.0071 | 99.9 | Highy Probable | 158 | 190.414551 |
| BMO1GPI7 | BspMO1_0090900.t1.1 | 0.751 | 99.9 | Highy Probable | 512 | 837.06189 |
| BMO1GPI8 | BspMO1_0119500.t1.1 | 0.972 | 99.9 | Highy Probable | 232 | 1594.880371 |
| BMO1GPI9 | BspMO1_0120200.t1.1 | 0.6151 | 99.9 | Highy Probable | 733 | 239.136703 |
| BMO1GPI10 | BspMO1_0126800.t1.1 | 0.0839 | 99.9 | Highy Probable | 357 | 258.695862 |
| BMO1GPI11 | BspMO1_0183500.t1.1 | 0.9153 | 100 | Highy Probable | 483 | 45.006592 |
| BMO1GPI12 | BspMO1_0213300.t1.1 | 0.7709 | 99.9 | Highy Probable | 139 | 528.32074 |
| BMO1GPI13 | BspMO1_0275300.t1.1 | 0.9899 | 99.9 | Highy Probable | 181 | 569.44812 |
| BMO1GPI14 | BspMO1_0277500.t1.1 | 0.9811 | 99.9 | Highy Probable | 579 | 7.428508 |
| BMO1GPI15 | BspMO1_0342900.t1.1 | 0.9442 | 100 | Highy Probable | 178 | 3517.976807 |
| BMO1GPI16 | BspMO1_0343500.t1.1 | 0.9454 | 100 | Highy Probable | 152 | 3709.012207 |
| BMO1GPI17 | BspMO1_0320700.t1.1 | 0.4404 | 99.9 | Highy Probable | 201 | 381.76651 |
| BMO1GPI18 | BspMO1_0407700.t1.1 | 0.0101 | 99.9 | Highy Probable | 448 | 186.392136 |
| BMO1GPI19 | BspMO1_0419800.t1.1 | 0.0032 | 99.9 | Highy Probable | 413 | 330.82 |
| BMO1GPI20 | BspMO1_0431600.t1.1 | 0.0466 | 99.9 | Highy Probable | 345 | 101.67 |

**Table VI.** Predicted AP2 proteins of *B.* MO1.

| Gene Name | Gene ID | Protein length (aa) | RNA Expression (TPM value) | Protein MW (kDa) | Domains |
| --- | --- | --- | --- | --- | --- |
| BMO1-AP2-1 | BspMO1_0177400.t1 | 629 | 4.00421 | 68.3 | AP2 |
| BMO1-AP2-2 | BspMO1_0027500.t1 | 680 | 52.212086 | 75.8 | AP2 |
| BMO1-AP2-3 | BspMO1_0036300.t1 | 742 | 384.286835 | 84.1 | AP2, RPT1 |
| BMO1-AP2-4 | BspMO1_0063100.t1 | 197 | 315.744598 | 23.3 | AP2 |
| BMO1-AP2-5 | BspMO1_000584-T1 | 74 | -- | 8.6 | AP2 |
| BMO1-AP2-6 | BspMO1_0136500.t1 | 375 | 46.351261 | 41.9 | AP2 |
| BMO1-AP2-7 | BspMO1_0155600.t1 | 401 | 722.822876 | 45.3 | AP2 |
| BMO1-AP2-8 | BspMO1_0164600.t1 | 581 | 179.783752 | 65.2 | AP2 |
| BMO1-AP2-9 | BspMO1_0196700.t1 | 932 | 456.914459 | 104 | AP2 |
| BMO1-AP2-10 | BspMO1_0206900.t1 | 513 | 178.840546 | 58.7 | AP2 |
| BMO1-AP2-11 | BspMO1_0279000.t1 | 488 | 388.687714 | 55.9 | AP2 |
| BMO1-AP2-12 | BspMO1_0297600.t1 | 475 | 1053.084351 | 53.8 | AP2 |
| BMO1-AP2-13 | BspMO1_0426700.t1 | 691 | 186.523087 | 75.3 | AP2 |
| BMO1-AP2-14 | BspMO1_0425100.t1 | 669 | 56.950947 | 75.6 | AP2. ACDC |
| BMO1-AP2-15 | BspMO1_0423600.t1 | 459 | 132.788696 | 52.2 | RPAP2_Rtr1 |
| BMO1-AP2-16 | BspMO1_003360-T1 | 794 | -- | 90.6 | AP2 |
| BMO1-AP2-17 | BspMO1_0103400.t1 | 261 | 159.928848 | 29.7 | AP2 |
| BMO1-AP2-18 | BspMO1_0109600.t1 | 214 | 107.731911 | 25.2 | AP2 |
| BMO1-AP2-19 | BspMO1_0112900.t1 | 148 | 20.662474 | 16.8 | AP2 |
| BMO1-AP2-20 | BspMO1_0138400.t1 | 408 | 84.778854 | 46.2 | PAP2_C |
| BMO1-AP2-21 | BspMO1_0377200.t1 | 541 | 77.83445 | 61.2 | AP2 |

### Comparative genomic and phylogenetic analyses of *Babesia* species reveal distinct genetic relationships and synteny patterns

The availability of genomic sequences from several Piroplasmids made it possible to conduct orthologous relationships between the genes of *B. divergens* Rouen 87, *B. divergens* 1802A, *B. bigemina, B. ovata, B*. MO1, *T. parva, B. duncani, B. bovis, B. microti,* and *B. sp.* Xinjiang. Our analysis identified 1,088 genes common to all species with a very high annotation percentage, 637 genes unique to *B.* MO1 aminly with unknown annotation, 223 genes unique to *B. divergens* 1802A strain, 188 genes unique to *B. divergens* Rouen 87 strain, and 516 genes shared among *B. divergens* 1802A, *B. divergens* Rouen 87, and *B.* MO1 (**Fig 2A**). Pairwise global alignment of the genomes showed the average nucleotide identity (ANI) between *B. divergens* 1802A and *B. divergens* Rouen 87 strains to be ∼99.1%, while the ANI between *B. divergens* Rouen 87 and *B.* MO1 is slightly lower at 96.7% (**Fig 2B**). A synteny analysis of *B.* MO1 against *B. duncani*, *T. parva*, *B. microti*, *B. divergens, B. bigemina, B. bovis* showed high synteny of *B.* MO1 with *B. divergens* Rouen 87, *B. bigemina*, and *B. bovis*, but lower synteny with *B. duncani*, *T. parva* and *B. microti* (**Fig 3**).

**Figure 2.**
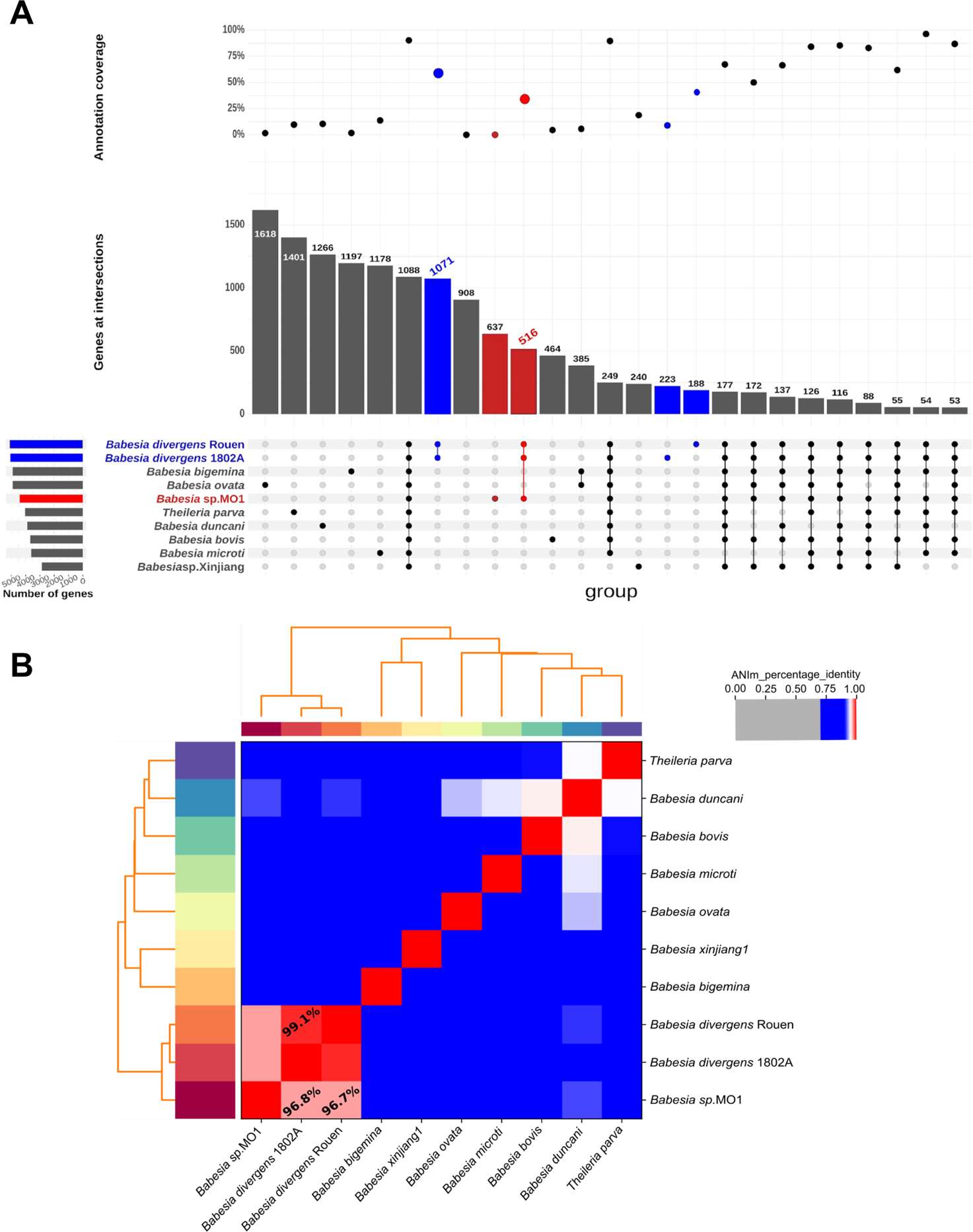
Evolutionary analysis of *Babesia* MO1 genome. **A.** Upset plot depicting orthogroups between *B.* MO1 and other apicomplexans. In the upper panel, the percentage of annotated proteins for shared or unique ones from a given organism is presented. In the middle panel, the total number of unique or shared proteins from a given organism is depicted. The lower panel represents the intersection or uniqueness of a given species with horizontal bars at the left side, representing the total number of genes for a given species. **B.** Heatmap of ANI values between *Babesia* species and *Theileria parva*. Higher values (red color) correspond to greater nucleotide similarity between the genomes.

**Figure 3.**
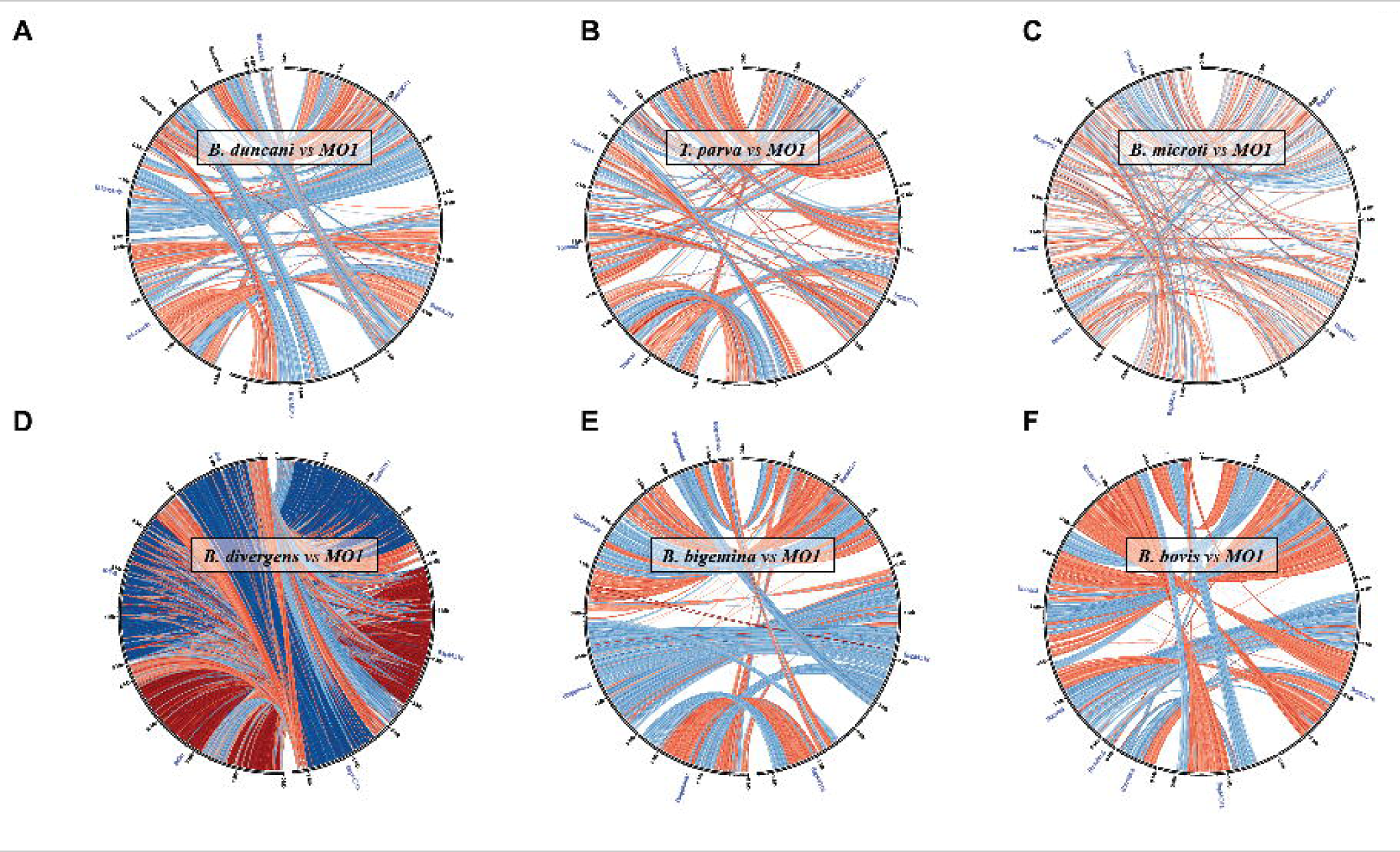
Circos synteny plots. The chromosomes of *B.* MO1 are illustrated on the right semicircle on all circular plots, and the chromosomes of the other organisms are on the left semicircle (**A**: *B. duncani*, **B**: *T. parva*, **C**: *B. microti*, **D**: *B. divergens* Rouen 87, **E:** *B. bigemina,* **F:** *B. bovis*); blue arcs indicate syntenies, red arcs indicate syntenies involved in a reversal; the intensity of the color is proportional to the level of collinearity; the number after the species’ name refers to the chromosome number (when chromosomes are broken into pieces, fragments).

Phylogenomic inference was employed to reconstruct the evolutionary history of *B. MO1* using supermatrix and supertree methods (details in Supplemental Methods). Two distinct sets of orthologous genes were considered in this study. Dataset 1 comprised ∼2500 orthologous groups, each with a single gene per isolate and a minimum of four sequences. Dataset 2 consisted of orthologous groups from dataset 1 but included only those with at least one one outgroup sequence. In cases where multiple outgroup sequences were available, it was imperative that they exhibited monophyly within the corresponding gene tree. Using Matrix Representation with Parsimony (MRP) method [32] on datasets 1 and 2, a single most parsimonious tree was generated. This tree received 100% support for each clade in dataset 2 and significant support for most clades in dataset 1 (**Fig 4A, Fig S5A & B**). The method confirmed that *B.* MO1 belongs to the *Babesia sensu stricto* clade VI, which includes *B. bigemina*, *B. bovis*, *B. caballi*, *B. divergens*, *B. ovata*, *B. ovis*, and *B. Xinjiang*. It also supported the placement of *B. MO1* outside the *B. divergens* subclade. To obtain confidence values for each branch, concordance factors were calculated from the source trees [33], providing 99% confidence for the clade containing *Babesia MO1*. More than 83% of the generated trees (85.5% for dataset 1 and 83% for dataset 2) supported the model indicating that *B. MO1* is likely a new *Babesia* species closely related to *B. divergens*. Consistent with previous studies, the MRP supertree method also confirmed *B. duncani* as a defining member of clade II [24]. Other computational approaches, such as PhySIC_IST, [34, 35] and Supermatrix, ran on dataset #2, further corroborated the distinct placement of *B.* MO1 from *B. divergens*, indicating their close yet distinct relationship and suggesting recent evolution (**Fig S5C and S5D**).

**Figure 4.**
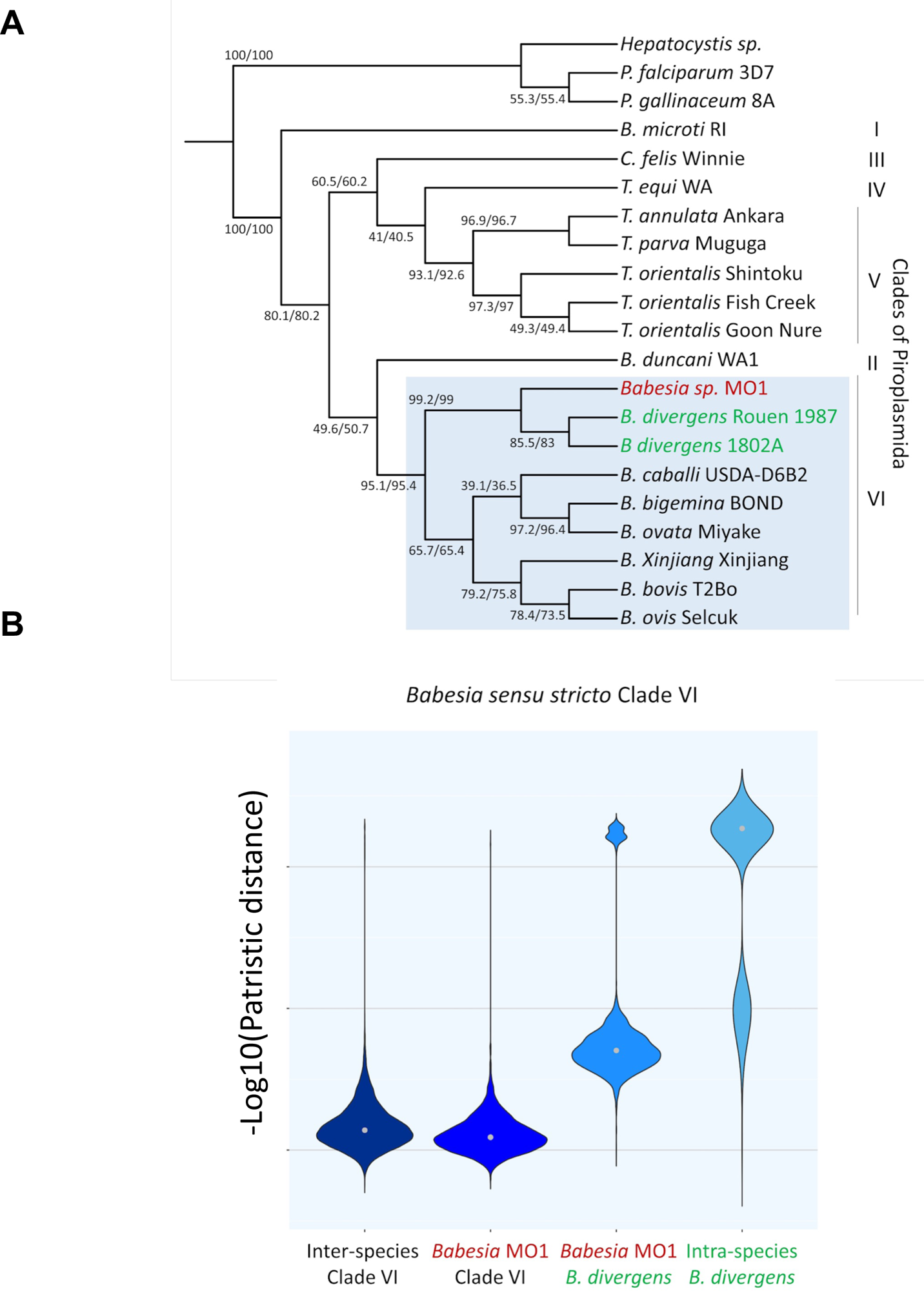
Piroplasmida species phylogeny inferred from phylogenomic analysis. **A.** Species phylogeny obtained by super matrix and super tree phylogenomic approaches. All bootstrap values with super matrix were at 100%. Displayed clade support values are estimated in the case of super tree methods by concordance factors from the source trees of dataset #1/dataset #2. The position of *Babesia* MO1 was analyzed in relation to the two *B. divergens* isolates (highlighted in green color in blue box). *B.* MO1 from the present study is in red (highlighted in blue box). *Hepatocystis* sp. (ex *Piliocolobus tephrosceles* 2019), *Plasmodium falciparum* 3D7 and *P. gallinaceum* 8A were taken as outgroup. **B.** Summary of the genetic exchanges between Piroplasmida species based on patristic distances. A matrix of patristic distances was calculated from the 2499 trees of dataset #1 for all pairs of species. Grey dot: median of the distribution. Comparisons between species of Clade VI, between *B.* MO1 and species of Clade VI, between *B.* MO1 and two strains *B. divergens*, and between two strains *B. divergens* are shown.

Patristic distances (PD) calculated from trees in dataset 1 were used to characterize the speciation between *Babesia MO1* and *B. divergens*, which appeared closely related in the species tree constructed through phylogenomic methods (**Fig 4B**). The distribution of – log10(PD) suggested recent evolution of *B.* MO1 from *B. divergens*, as the distances were greater between *B*. MO1 and *B. divergens* than between different isolates of *B. divergens*. The evidence for recent speciation was strengthened by the observation that the genetic distance between *B.* MO1 and *B. divergens* was shorter than the distance between *B. MO1* and other *Babesia* species belonging to Clade VI. Using PD values, *B. MO1* genes were then categorized into low, medium, and high groups (**Fig S6 and B**). Approximately 75 genes were found to evolve closely among 22 gene ontology (GO) identities (IDs). Genes in the low LD group (1e-04<PD<0.011) were associated with processes such as protein folding and quality control, particularly those occurring in the endoplasmic reticulum. Conversely, some genes in the high-distance genes (PD>0.29) were linked to mRNA maturation and degradation. This analysis further identified specific metabolic processes, such as pyrimidine and isoprenoid biosynthesis pathways, that show distinct evolution in both organisms, suggesting possible differences between *B. MO1* and *B. divergens* in their cellular metabolism and adaptation to host environments (**Fig. S13**)

### *B.* MO1 and *B. divergens* mitochondrial and apicoplast genomes

The mitochondrial and apicoplast genomes of *B.* MO1 were further analyzed and compared to those of *B. divergens*. The mitochondrial genome of *B.* MO1 is a linear molecule spanning 6.3 kb, while its apicoplast genome is circular, comprising 29.3 kb. The sizes of both mitochondrial and apicoplast genomes in *B. divergens* closely mirror those of *B.* MO1. The apicoplast genomes in both species are circular molecules measuring 29.3 kb for *B.* MO1 and 29.9 kb for *B. divergens*, with A+T content of 86.4% and 86.6%, respectively. Notably, the apicoplast genome of *B.* MO1 contains twenty-seven ORF genes, while *B. divergens* has twenty-six. The *B.* MO1 apicoplast genome includes sixteen ribosomal proteins, twenty-three tRNAs, two ribosomal RNAs (LSU and SSU), five RNA polymerases, and five additional proteins (ClpC1, ClpC2, and TufA) (**Fig 5A**). In contrast, the *B. divergens* apicoplast genome comprises seventeen ribosomal proteins, twenty tRNAs, two ribosomal RNAs (LSU and SSU), seven RNA polymerases, and five other proteins (ClpC1, ClpC2, hp3, hp5, and TufA). Some apicoplast-encoded transcripts in *B. divergens* are polycistronic, including rps2, rps3, RpoB, and RpoC1 (**Fig 5B**). The mitochondrial genomes of *B. MO*1 and *B. divergens* are characterized as monocistronic with sizes of 6326 bp and 6323 bp, respectively. Both mitochondrial genomes encode four genes (*cob, coxI, coxIII*, and *nad2*) and five tRNAs (**Fig 5C and D**). Additionally, the *B.* MO1 mitochondrial genome codes for seven rRNAs, while the *B. divergens* mitochondrial genome codes for six rRNAs (**Fig 5C and D**).

**Figure 5.**
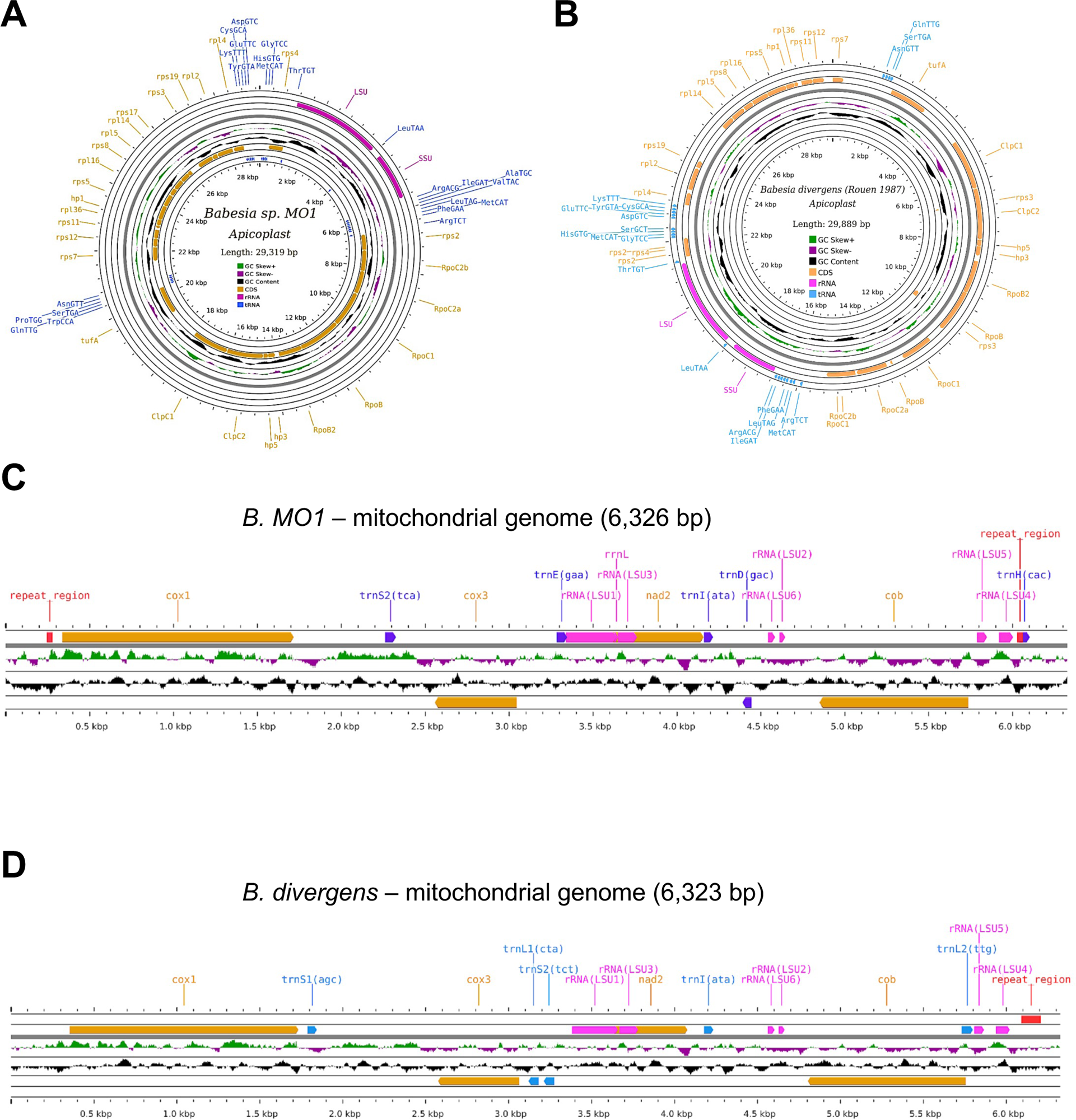
Apicoplast and mitochondrial genomes of *Babesia* MO1 and *B. divergens*. **A-B.** Graphical circular map of the apicoplast genome of *B.* MO1. and *Babesia divergens* Rouen 1987, respectively. **C-D.** Linear map of the mitochondrial genome of *B.* MO1 and B. divergens Rouen 1987, respectively. Orange arrows represent genes encoding proteins involved in the electron transport chain, including *cox1*, *cox3*, *nad2*, and *cob*. The genes encoding ribosomal RNA (rRNA) are depicted in pink color. Different tRNA encoding genes are displayed in purple color.

### Regulation of gene expression, epigenetics, and chromatin structure in *B*. MO1

To gain further insights into the biology of *B.* MO1, total mRNA was extracted for RNA-seq experiments for both F12 and B12 clones. After library preparation and sequencing, we mapped the resulting reads and calculated the expression (Transcripts Per Million (TPM)) of all genes from *B.* MO1 (both clones) and plotted them in **Fig 6 A and B**. We then binned average expression of genes across the 3 chromosomes in 50-kb bins, which were color-coded on average normalized gene expression values. (**Fig. 6C and 6D**). Similar to what was observed in several apicomplexan parasites including *B. duncani* that possess gene families involved in antigenic variation, a significant relationship between gene expression and the telomeres was detected with a significant decrease in the expression of genes localized near the telomeres [23, 24]. All telomeres except for the left end of chromosome II harbor several clusters of genes belonging to the *B.* MO1 MGF families (**Fig. 6B and 6C**) indicating that these genes may be repressed by a heterochromatin cluster allowing for possible mono-allelic expression of the MGF as described in *P. falciparum* [36]. Using the TPM expression values (see experimental procedures in Methods), we found that the total range of transcriptional activity captured using the continuous in vitro growth conditions varied by more than four orders of magnitude, from 0 to over 10,000 (See supplemental data). Overall, RNA-seq data captured 4540 (99.4%) of the 4569 predicted annotated genes in the assembled *B.* MO1 genome with greater than 0 TPM, and 4078 (89.3%) with greater than 10 TPM, indicating that most genes in both clones are expressed during the intraerythrocytic life cycle of the parasite and are potentially needed for parasite survival in the host red blood cells. Not surprisingly, among the most highly expressed genes were genes involved in translation most prominently, with many ribosomal proteins among those with the very highest TPM. Other highly expressed genes were those involved in the ubiquitin proteasome system, cell cycle, ATP hydrolysis-coupled proton transport, as well as histone core proteins indicating active metabolic activity and maintenance of the parasite by standard housekeeping genes. Amongst the 491 genes that were found to have fewer than 10 TPM (likely silenced during the intraerythrocytic life cycle in vitro), nearly all were genes that did not have an obvious match to other organisms by BLAST and are thus not currently assigned a specific function, although many (213 of 491, 43.4%) are members of the VESA1, VESA2, or UMGF multi-gene families idenfitied for *B. MO1.* These MGF genes with less than 10 TPM represent 50.6% of the 421 total MGF genes, and 347 (82.4%) have less than 50 TPM. Interestingly, 15 of the multi-gene family genes have over 300 TPM, placing them in the top 1000 most highly expressed genes, and perhaps indicative of an antigenic variation mechanism where only a small number of genes in these families are highly expressed at any given time. Although the above results are for clone F12, the same patterns seem to hold for B12 as well, with figures 6A-E showing that the two clones have similar expression patterns across the genome.

**Figure 6.**
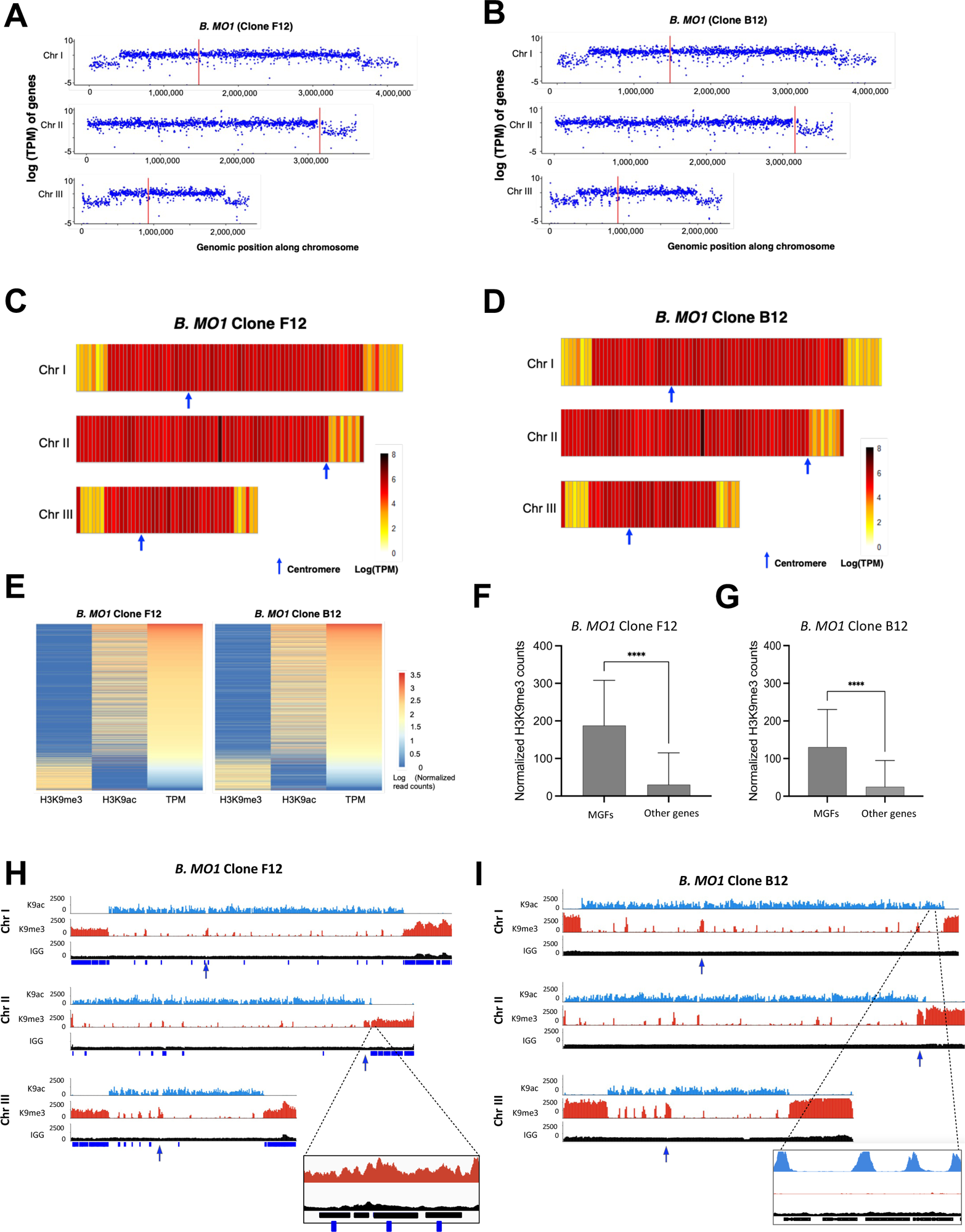
Transcriptomic profile and epigenomic landmarks of *B.* MO1. **A-B.** Logarithms of the TPM counts in *B.* MO1 clones F12 **(panel A**) and B12 (**panel B**) were used as expression values for each gene across the three chromosomes using the R package ggplot2. **C-D.** RNA-seq data of *B.* MO1 clones F12 **(panel C**) and B12 (**panel D**) as normalized heat maps across the three chromosomes. Chromosomes were divided into 50 kb bins and the average of the log TPM of genes within each bin was calculated. *n*LJ=LJ2 biologically independent samples. **E.** Comparison between epigenetic marks and gene expression in *B.* MO1 clones F12 and B12. Heat maps were built using normalized log_2_H3K9me3 and H3K9ac read counts in addition to the RNA-seq TPM levels of each gene. Read counts for H3K9me3 and H3K9ac were normalized to millions of mapped reads and gene length, whereas TPM was determined by Stringtie. Genes were ranked from high to low TPM highlighting the correlation and anti-correlation between transcript abundance and the H3K9ac3 and H3K9me3 marks, respectively. **F-G**. Normalized H3K9me3 counts in multigene families, and other genes encoded by *B.* MO1 clones F12 (**panel F**) and B12 (**panel G**) (unpaired t-test with Welch’s correction, P<0.0001) n = 2 biologically independent samples. **H-I**. Heterochromatin and euchromatin distribution across the three chromosomes of *B.* MO1 clones F12 (**panel H**) and B12 (**panel I**). Tracks correspond to H3K9ac3 ChIP (top), H3K9me3-ChIP (middle), and IgG control (bottom) and were normalized to millions of mapped reads. n=2 biologically independent samples.

This dataset was further examined in both clones to mine and identify additional molecular components that could be critical to the parasite life cycle progression. Of the reads that mapped against the *B.* MO1 genome, 78.6% mapped with at least 90% overlap to predicted protein-coding gene models, 7.02% mapped within 300-bp upstream of genes, and 7.47% mapped within 300-bp downstream of genes. In addition, 5.76% of reads fall entirely within intergenic regions only, outside of even upstream and downstream regions. The upstream and downstream mapped reads demarcate possible UTRs, while those mapped within intergenic regions only could represent long non-coding RNAs (lncRNAs). LncRNAs are non-protein coding transcripts that have been shown to play a critical role in biology including cell differentiation and sexual differentiation throughout changes in epigenetics and chromatin structures [37–39]. LncRNAs have also been implicated in the regulation of genes involved in antigenic variation in human malaria parasites [40, 41]. The RNAs mapping outside the annotated genes represent candidates for lncRNAs that can be explored in the future to complete the true transcriptome of *B.* MO1.

To further examine the possible relationship between epigenetics and gene expression, we conducted chromatin immunoprecipitation assays (ChIP) followed by next generation sequencing on both clones in duplicates to identify the localization of specific histone marks and their association with gene expression. ChIP was conducted using antibodies against tri-methylated histone 3 lysine 9 (H3K9me3) and acetylated histone 3 lysine 9 (H3K9ac) as markers for heterochromatin and euchromatin marks, respectively. The immune precipitated DNA and input used as a positive control were purified, amplified, and subjected to next-generation sequencing on the Illumina Novaseq sequencing platform. Reads were mapped to the *B.* MO1 clone F12 and B12 genomes and normalized per million of mapped reads for each sample. Very high Pearson correlation coefficients within each ChIP-seq pair of replicates confirm the reproducibility of our experiment (**Table S2A and S2B**). Negative correlation coefficients between H3K9me3 and H3K9ac samples, as well as genome-wide tracks showing mapping patterns, demonstrate that, similarly to what is observed in eukaryotes including apicomplexan parasites, euchromatin and heterochromatin marks are mutually exclusive (**Fig 6H** and **6I**). We also confirmed a large heterochromatin cluster near the telomeric and sub telomeric regions of all chromosomes surrounding multigene families. Transcription of multigene families where genes are repeated in tandem is responsible for the presence of GC-skew in these regions (**Fig S10**). Statistical analyses were used to determine if genes from multigene families were significantly enriched with H3K9me3 marks compared to other genes encoded in the *B.* MO1 genome for both clones. Our data demonstrate that like what was observed in *B. duncani*, genes that belong to multigene families (MGF) in the *Babesia* MO1 clone genomes are significantly enriched in H3K9me3 marks (**Fig 6F** and **6G**). Our analysis identified many genes marked by histone H3K9me3, most of them localized in the telomeric ends, annotated as hypothetical proteins with no homologs in other organisms. Considering their genomic localization and their strong enrichment in heterochromatin marks, these genes could also be involved in immune evasion. Additional histone H3K9me3 marks were also observed in genes throughout the genome (**Fig 6H** and **6I**) and correlate perfectly with genes not expressed during the erythrocytic stage that could be involved in either immune evasion or cell differentiation including sexual differentiation. The euchromatic mark, H3K9ac, on the other hand, is detected on all other chromosomal regions in both clones and found to be enriched in the promoters of active genes (**Fig 6H** and **6I)** and their intensity correlates with transcript abundance (**Fig 6E**). Our transcriptomic and epigenetic study further confirms that epigenetic marks correlate with gene expression and that silencing is associated with repressed genes either involved in sexual differentiation or multigene families most likely involved in antigenic variation.

To further investigate the effect the MGF such as *vesa g*enes have on the overall chromatin organization, we performed chromatin conformation capture (or Hi-C) experiments on the parasite chromatin for both *B.* MO1 clones. Hi-C libraries for each sample (F12 and B12) were prepared independently in duplicate as previously described [22, 24, 42] and sequenced to a mean depth of ∼98.4 million reads for clone F12 and 119 million reads for clone B12. The libraries were processed using HiC-Explorer [43] and resulted in ∼29.6 million valid interaction pairs for clone F12 and ∼49.6 million contacts for clone B12. To identify intrachromosomal and interchromosomal interactions, we selected to bin our reads at a 10-kb resolution. The contact map for *B.* MO1 from F12 and B21 clones are shown on supp Figs. S7A and S8A respectively. A close examination of the contact maps indicate that the genome assembly has no major large mis-joints or mis-assemblies in the chromosome cores, but many reads could not be mapped in the sub telomeric or highly repetitive regions and is consistent with what was also observed to a lesser extend in the *P. falciparum* genome [23]. When successfully mapped, all sub telomeric regions or regions mapped to potential multi gene families or heterochromatin marks were however detected as strongly interacting with each other confirming the formation of a possible heterochromatin cluster for most identified MGFs. This was further confirmed by overlapping our Hi-C and ChIP-seq data against the histone H3K9me3 (**Fig. 7A and 7B**). We also detected that the centromeres that exhibit acrocentric profile interact with each other and present a distinct pattern between *B.* MO1 (F12 and B12 clones) and *B. divergens* (see supp **Figs. S7, S8, and S9**). To confirm the genome-wide chromatin organization of *B.* MO1, a 3D model was constructed using PASTIS [44] from the Hi-C contact maps **(Fig. 7C and 7D**)). We also built a 3D model for Hi-C data generated for *B. divergens* **(supp Fig. 12**). In all models the three chromosomes of *B.* MO1 and *B. divergens* showed that the centromeres and heterochromatin/telomes cluster together in distinct regions within the nucleus, (**Fig. 7C and 7D**) an organization similar to what was reported in apicomplexan parasites including that of the *B. microti* and *B. duncani* genomes [24] The strong co-localization of genes with H3K9me3 marks that included most babesia MGFs confirming a tight control of *vesa* and MGF gene regions at the epigenetics and chromatin structure levels (**Fig 7A and 7B**).

**Figure 7.**
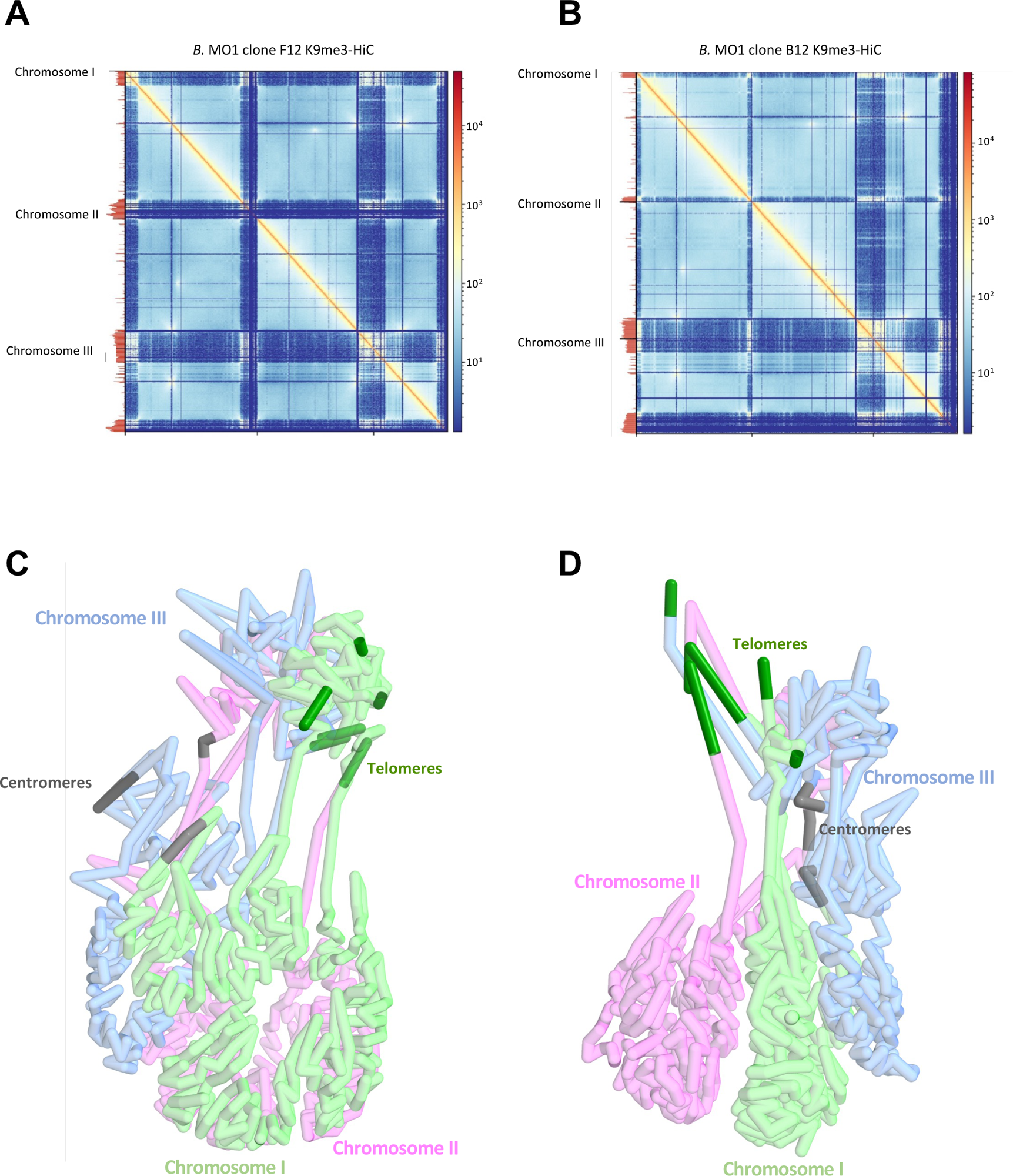
*Babesia* MO1 3D-genome. **A-B**. Hi-C contact maps coupled with H3K9me3 ChIP-seq tracks (left) of *B.* MO1 clones F12 and B12 (10-kb kb bins). Tracks are scaled to chromosome lengths. **C-D**. 3D genome structures of *B.* MO1 clones F12 and B12 derived from the contact map interactions. Chromosomes one, two, and three correspond to green, pink, and blue sections respectively. Dark green and grey represent the telomeric regions and centromeres.

**Figure 8.**
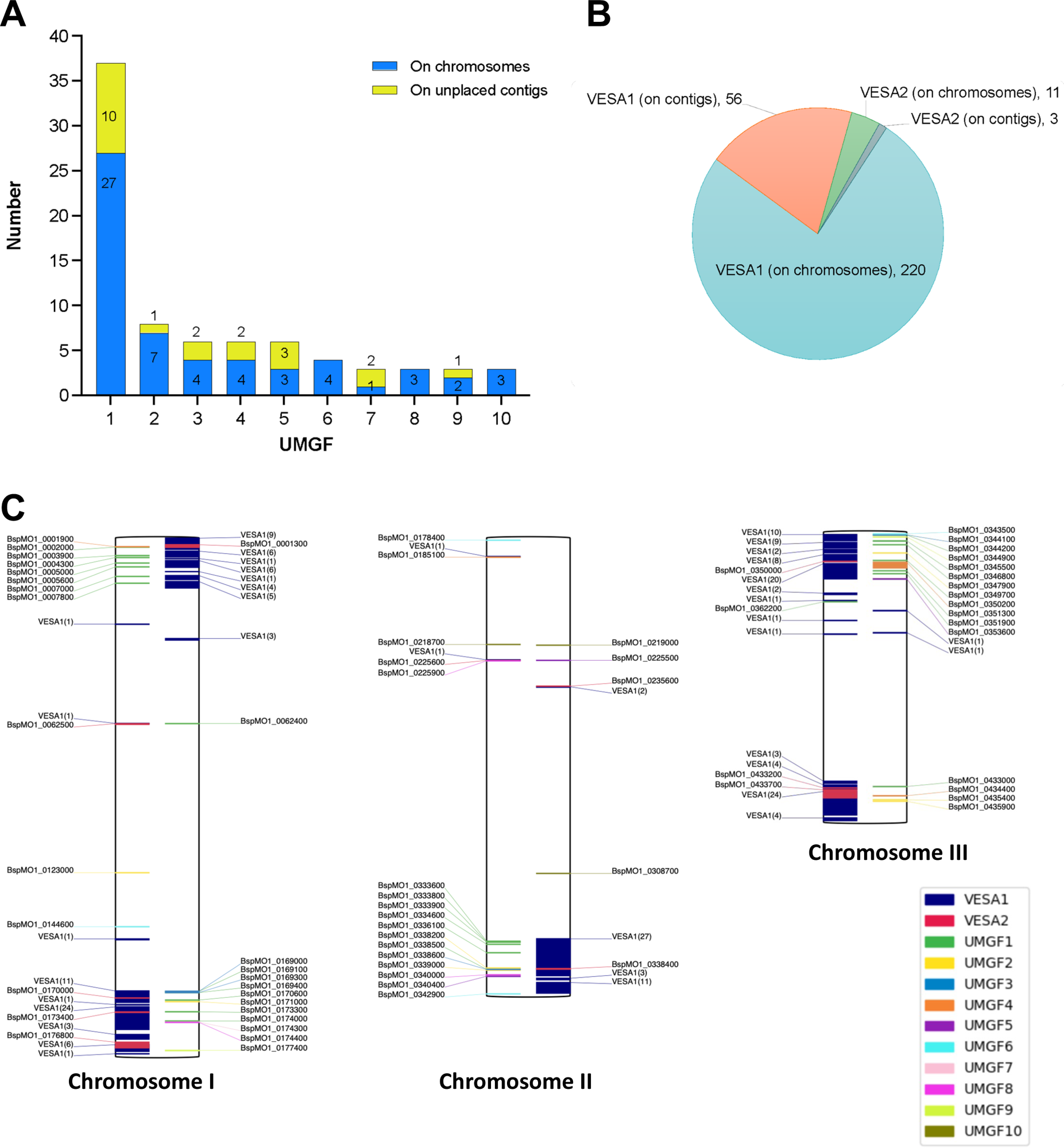
Multi-gene families of *B.* MO1 and their chromosomal localization. **A.** Plot depicting the unique multigene families (UMGFs) in *B.* MO1. The blue bars depict the genes localized on one of the three chromosomes, whereas the yellow bars denote the genes found on stray contigs. **B**. Distribution of *B.* MO1 *vesa*1 and *vesa*2 genes on either chromosomes or stray contigs. **C.** Localization of *vesa* genes and UMGFs members on the three *B.* MO1 chromosomes (genes localized on unplaced contigs are ignored). Genes denoted on the right side of a chromosome are on the positive strand, whereas those shown on the left side are on the negative strand.

### Evolution of multigene families in *B.* MO1 and *B. divergens*

A previous study in *B. divergens* identified 359 *ves* gene encompassing three subfamilies namely, *ves1* (n=202), *ves2a* (95), and *ves2b* (62) (**Table VII**) [18]. In our reannotated genome of *B. divergens* Rouen strain, we identified only 134 *vesa* genes. Interestingly, *B.* MO1 expresses 290 *vesa* genes: 276 of those had a C-terminal domain (*vesa1*) while the remaining 14 did not (*vesa2*). The *vesa* genes in *B.* MO1 encode proteins with an average of 617.1 aa (standard deviation 486.6 aa) for *vesa1* and an average of 295.8 aa (standard deviation 240.2 aa) for *vesa2*. In addition to this family of genes, our analysis identified 10 novel gene families (unique multigene families; UMGFs) with at least three members. Most members of these families localize to the highly repetitive telomeric regions, the largest of which, UMGF1 (unique multigene family 1), consists of 37 members, 27 of them successfully mapped to the telomeric regions of chromosomes I-III, and the remaining 10 mapped to unassembled contigs (**Fig 8A, 8B**). The second largest family, UMGF2, consists of 8 members, of which 7 members mapped to the telomeric regions of one of the three chromosomes; one was mapped to unassembled contigs (**Fig 8A, 8B**). No homologs of these proteins are found in other apicomplexan parasites, but their genome localization is reminiscent of the localization of gene families involved in antigenic variation in other parasites including the *var* genes in *P. falciparum* [21–23, 45] and or the VSG in *Trypanosoma brucei*) [46]. The role of these new gene families in parasite adaptation to its mammalian host and/or vector remains to be elucidated.

**Table VII.**
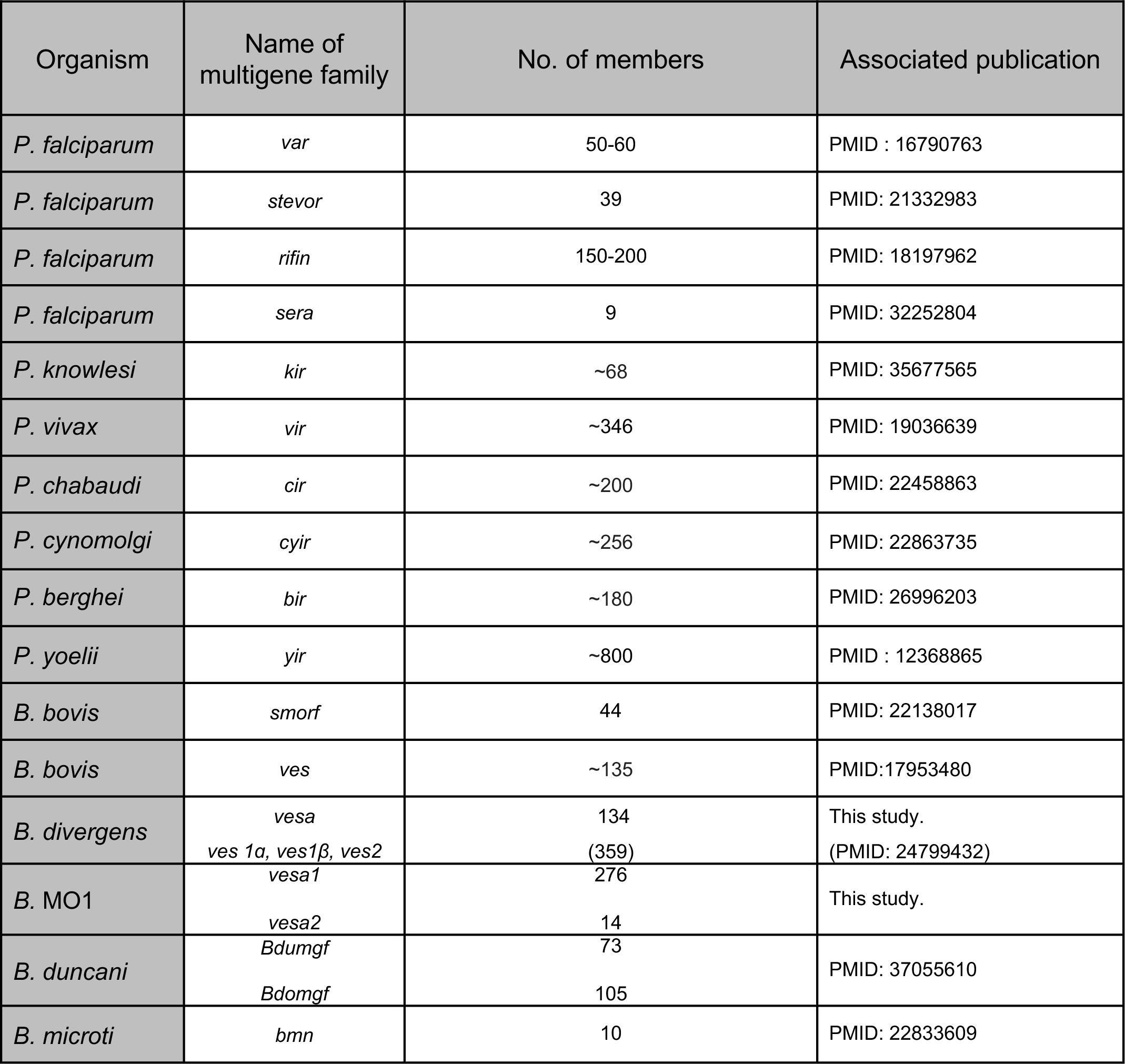
Multigene families in different apicomplexan parasites.

### *B.* MO1 and *B. divergens* display distinct susceptibility to antibabesial drugs

We conducted a comparative analysis of the susceptibility of *B.* MO1 and *B. divergens* to currently approved antibabesial drugs, including atovaquone, azithromycin, clindamycin, quinine, as well as antifolate drugs WR99210 and pyrimethamine. The data revealed that *B.* MO1 is ∼2.4-fold, ∼1.2-fold, 1.3-fold, and ∼2.9-fold less susceptible to atovaquone, azithromycin, clindamycin, and pyrimethamine, respectively, compared to the *B. divergens* Rouen 87 isolate (**Fig 9** and **Table VIII**). Conversely, *B.* MO1 displayed, 2.7-fold, and ∼160-fold greater sensitivity to quinine, and WR99210 than the *B. divergens* Rouen87 isolate. In various parasites, the mitochondrial-encoded *cyst* gene, and the nuclear-encoded genes *rpl6* and *dhfr-ts* have been established as the molecular targets for atovaquone, clindamycin, WR99210, and pyrimethamine, respectively. However, our analysis showed that the primary sequences of these enzymes are highly conserved between *B. divergens* and *B.* MO1, suggesting that polymorphism within their encoding genes might not account for the differences in drug susceptibility between the two species (**Fig. S11**). Interestingly, RNA sequencing analysis revealed significant differences in the expression levels of the genes encoding key enzymes involved in folate metabolism (**Table IX**). Notably, the expression levels of glutathione synthase (GS) showed an approximately 10-fold difference, and dihydropteroate synthase (DHPS) exhibited an approximately 12-fold difference (**Table IX**). These differences in gene expression levels might thus contribute to the differences in drug susceptibility observed between the two species.

**Figure 9.**
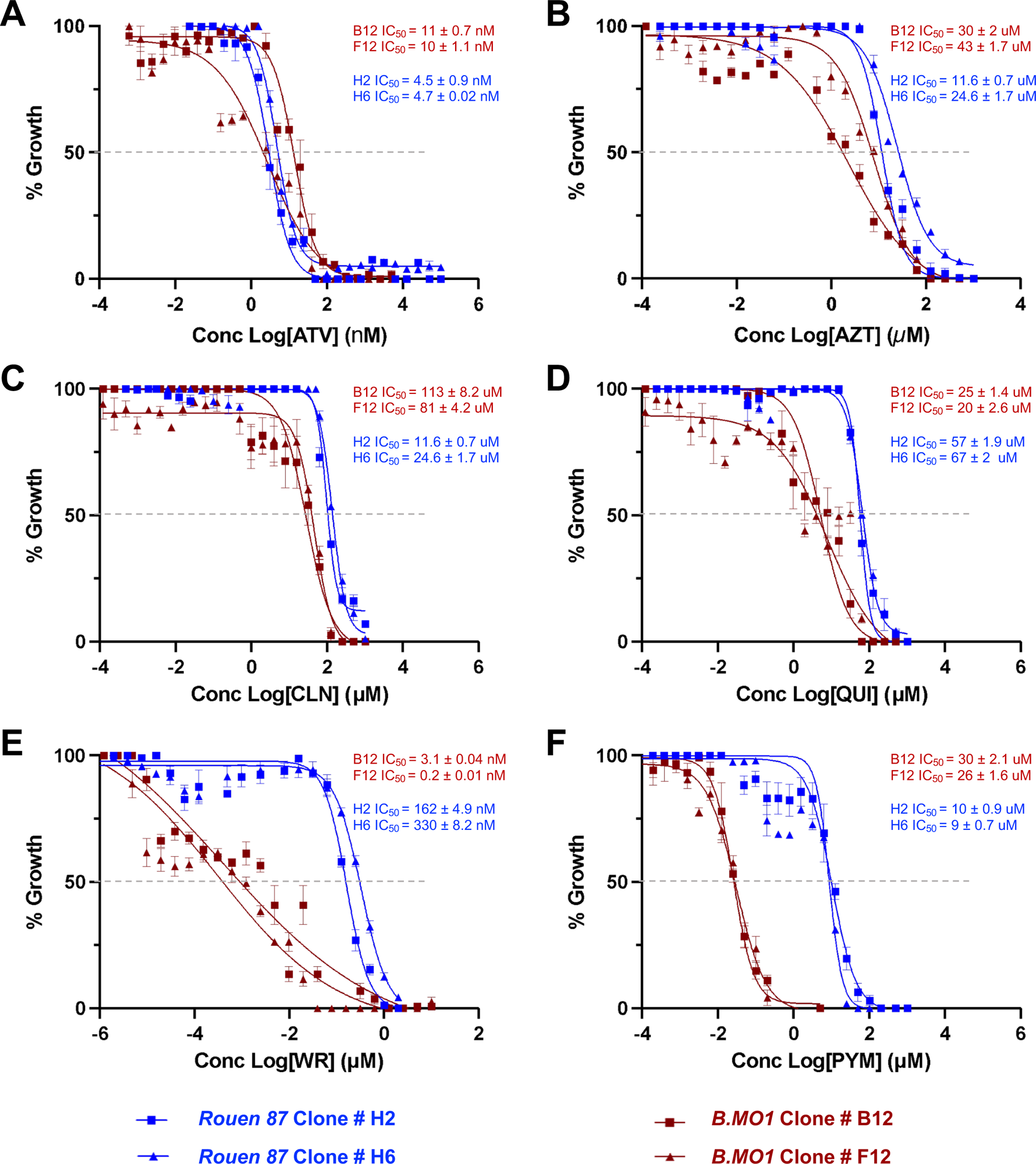
In vitro efficacy of current antibabesial compounds against *B.* MO1 and *B. divergens* Rouen 87. **A-F.** Potency and IC_50_ determination of Atovaquone (ATV), Azithromycin (AZT), Quinine [46], Clindamycin (CLN), WR99210 (WR), and Pyrimethamine (PYM) against *B. divergens* Rouen 87 clones H2 and H6, and *B.* MO1 clones B12 and F12. Data presented as mean ± SD of three independent experiments performed in biological triplicates.

**Table VIII.** Comparison of half minimal inhibitory concentration (IC_50_) of various antiparasitic drugs between clones of *B.* MO1 and *B. divergens* Rouen 87.

| Antiparasitic drug<br>(Target) | *B. MO1* B12 | *B. MO1* F12 | *B. divergens*<br>Rouen87<br>Clone H2 | *B. divergens*<br>Rouen87<br>Clone H6 | Fold<br>difference |
| --- | --- | --- | --- | --- | --- |
| Atovaquone (Cyt-b) | 11 ± 0.7 nM | 10 ± 1.1 nM | 4.5 ± 0.9 nM | 4.7 ± 0.02 nM | 2.4 |
| Azithromycin (RPL6) | 30 ± 2 uM | 43 ± 1.7 uM | 11.6 ± 0.7 uM | 24.6 ± 1.7 uM | 1.2 |
| Clindamycin | 113 ± 8.2 uM | 81 ± 4.2 uM | 11.6 ± 0.7 uM | 24.6 ± 1.7 uM | 1.3 |
| Quinine | 25 ± 1.4 uM | 20 ± 2.6 uM | 57 ± 1.9 uM | 67 ± 2 uM | 2.7 |
| WR99210 (DHFR-TS) | 3.1 ± 0.04 nM | 0.2 ± 0.01 nM | 162 ± 4.9 nM | 330 ± 8.2 nM | 164 |
| Pyrimethamine( DHFR-TS) | 30 ± 2.1 uM | 26 ± 1.6 uM | 10 ± 0.9 uM | 9 ± 0.7 uM | 2.9 |

**Table IX.** RNA -seq TPM values of folate metabolism genes.

| Gene Name | *B. MO1*<br>Clone B12 | *B. MO1*<br>Clone F12 | *B. divergens*<br>Rouen 87 |
| --- | --- | --- | --- |
| Serine hydroxymethyltransferase-1 (SHMT) | 290.896484 | 255.062485 | 108.65 |
| S-adenosylmethionine synthase-2 (SAMS) | 87.162872 | 81.135635 | 336.04 |
| Glutathione synthetase (GS) | 224.737961 | 232.131134 | 22.56 |
| Dihydrofolate reductase thymidylate synthase (DHFR-TS) | 297.84668 | 304.414795 | 382.95 |
| Adenosyl homocysteinase (AHC) | 238.004715 | 240.822906 | 328.56 |
| Dihydropteroate synthase (DHPS) | 214.073288 | 199.632187 | 16.59 |

## Discussion

The results presented in this study provide valuable insights into the biology, genomics, and epigenetics of both *B.* MO1 and its close relative, *B. divergens*. These findings reveal striking differences in the replication rates, transmission dynamics, genomic characteristics, and susceptibility to antibabesial drugs between these two pathogens. The data, which substantiate the notion that these organisms are distinct but closely related species, underscore the critical importance of understanding the intricacies of these parasites, particularly in the context of their evolution and the potential for zoonotic transmission to humans.

First, we found that the two organisms display major differences in replication rates and dynamics under similar experimental growth conditions. Differences in the strategies between *B.* MO1 and *B. divergens* to produce daughter parasites during each intraerythrocytic life cycle in human RBCs also suggest that *B. divergens* is better adapted to these host cells compared to *B.* MO1. These variations could have implications for the severity of infection and the potential for these parasites to proliferate within their respective host populations. The differing transmission pathways, involving different tick vectors (*Ixodes dentatus* for *B.* MO1 and *I. ricinus* for *B. divergens*) and animal reservoirs (cottontail rabbits for *B.* MO1 and cattle for *B. divergens*) highlight the complex ecological interactions shaping the epidemiology of these parasites, and suggest niche specialization. Understanding these host-vector relationships and transmission cycles is crucial for devising effective control measures and assessing the risk of human infections.

Second, at the genomic level, our analysis revealed differences in chromosomal organization, both within and between *B.* MO1 and *B. divergens* isolates. While the genome size and chromosome numbers are consistent between the two organisms, the patterns observed in pulse field gel electrophoresis demonstrated varying chromosome sizes, suggesting chromosomal rearrangements. Interestingly, differences between the parental isolates and clones generated from single infected erythrocytes were also observed, indicating that both *B.* MO1 and *B. divergens* undergo dynamic polymorphism during their asexual development, likely the result of extensive mitotic recombination events.

Third, the genome assembly of *B.* MO1 and *B. divergens*, while achieving a high-level resolution, presented challenges, especially in fully assembling repetitive telomeric ends, despite the use of long read sequencing and optical mapping technologies. This emphasizes the need for improved methods to capture and assemble repetitive genomic regions accurately. Our analysis of the genomes of *B.* MO1 and *B. divergens* highlighted telomeric regions as primary source of chromosome size variation observed in PFEG, genetic variation and the location of several genomic rearrangements. Furthermore, our analysis of Average Nucleotide Identity (ANI) values and the number of orthologous proteins between *B.* MO1 and *B. divergens* strains provides compelling evidence in support of classifying *B.* MO1 as a distinct species. Genome relatedness indices, such as ANI, offer a rapid and readily applicable means of comparing genomes to delineate species boundaries. In prokaryotes, a 95% cutoff value is well-established for grouping genomes of the same species, but of ANI distribution and cutoff values for eukaryote species delimitation have not yet been fully defined. Nevertheless, the ANI value between *B. divergens* strains (99.1%) significantly exceeds the values observed between any *B. divergens* strain and *B.* MO1 (96.8% or 96.7%, respectively). Additionally, the number of orthologs shared between *B. divergens* strains (1,071 proteins) is higher than the count shared with *B.* MO1 (516 proteins). The sequence divergence between *B.* MO1 and *B. divergens* results in several proteins that are unique to each organism (637 proteins in *B.* MO1 and 223 or 188 in *B. divergens* strains), likely tied to their specific evolution and adaptation to their respective hosts. Furthermore, our genome assemblies were crucial in exploring the evolution and function of unique proteins encoded by multigene families, such as the previously described members of the *vesa* gene family found in both *B.* MO1 and *B. divergens*. However, several multigene families remain with unknown functions and need further experimental characterization to elucidate their role in each parasite. Altogether these findings highlight the genetic diversity within these parasites and offer insights into potential genetic adaptations to specific host niches.

Fourth, RNA-seq, ChIP-Seq and Hi-C analyses revealed important differences in gene expression and regulation between *B.* MO1 and *B. divergens*. For example, most of the multigene families were found to be transcriptionally silent and maintained in a large heterochromatin structure, a profile similar to that of other genes involved in antigenic variation from other apicomplexan parasites. These differences in chromosomal organization were further corroborated at the epigenetics and chromatin structure levels (**Fig 6 and 7**), suggesting that recombination events within heterochromatin clusters may have facilitated sub telomeric variations and the potential expansion and evolution of *vesa* genes in the analyzed clones and strains. Previous research has already noted a high incidence of mutations and sub telomeric instability in highly variable genes, such as *var* genes in the human malaria parasite, *P. falciparum* [47].

Another important finding in this study, is the finding of major differences drug susceptibility between *B.* MO1 and *B. divergens*. The differences emphasize the necessity of considering species-specific variations when designing therapeutic interventions.

In conclusion, this comprehensive study significantly advances our understanding of the biology and genomics of *B.* MO1 and *B. divergens*. The findings have implications for public health, emphasizing the need for tailored approaches to prevent and manage infections caused by these parasites. Additionally, the identification of potential new species and the exploration of drug susceptibility contribute valuable knowledge to the broader field of parasitology and infectious diseases. Future research should further investigate the molecular mechanisms underlying the observed differences and explore the ecological factors influencing the epidemiology of these *Babesia* species.

## Materials and Methods: (*Additional methods are in Supplemental Methods*)

### Ethics statement

*Babesia* MO1*, B. divergens* Rouen 87 and a *B. divergens* clinical isolate from Spain were cultured using human A^+^ blood obtained from healthy volunteer donors [6]. The blood was sourced from the American red cross (US), the Interstate Blood Bank (US), or the Blood Transfusion Center (Spain), adhering to approved protocols and in compliance with the relevant institutional guidelines and regulations.

### Continuous *in vitro* culture of *B.* MO1 *and B. divergens* in human red blood cells

*B.* MO1 parasites were initially obtained from BEI Resources (BEI Resources, NR-50441) and cultured in the HL1 medium (Lonza, 344017). Subsequently, we discovered that, similar to *B. duncani* [48], the parasite can be continuously propagated in complete DMEM/F12 medium or RPMI 1640 medium. The medium consists of either DMEM/F12 and RPMI1640 media (Lonza, BE04-687F/U1; Gibco-Life Technology, 11875093) supplemented with 20% heat-inactivated FBS (Sigma, F4135) or 0.5% albumax I (Thermofisher Scientific, 11020021), 2% 50X HT Media Supplement Hybrid-MaxTM (Sigma, H0137), 1% 200 mM L-Glutamine (Gibco, 25030-081), 1% 100X Penicillin/Streptomycin (Gibco, 15240-062) and 1% 10 mg/mL Gentamicin (Gibco, 15710-072)) in 5% hematocrit A^+^ RBCs. The parasite cultures were maintained at 37°C under a 2% O_2_ / 5% CO_2_ / 93% N_2_ atmosphere in a humidified chamber. Culture medium was changed every 24 h, and parasitemia was monitored by examining Giemsa-stained blood smears using a light microscope. *B. divergens* parasites (Bd Rouen 1987 strain and the clinical isolate from Spain) were cultured *in vitro* in human A^+^ RBCs and complete medium consisting of RPMI 1640 (Gibco-Life Technology, 11875093) supplemented with 0.5% Albumax II (Gibco, 11021037), 7.5% (w/v) sodium bicarbonate solution (Lonza Group Ltd, Basel, Switzerland, 144-55-8), and 100 μmol/L hypoxanthine (Sigma-Aldrich Corporation, St Louis, MO, H9377) at a pH of 7.3 at 37°C in a humidified atmosphere of 5% CO_2_ [21]. The culture medium was replaced every 24 h, and parasitemia was monitored by examining Giemsa-stained blood smears by a light microscope.

### Gene prediction and annotation of *B.* MO1 and *B. divergens*

The *Babesia* MO1 genome was processed using the gene annotation pipeline FunAnnotate v1.8.9 (https://github.com/nextgenusfs/funannotate) and PAP (https://github.com/kjestradag/PAP) pipelines. FunAnnotate was supplied with the MO1 IsoSeq isoforms computed above, along with protein sets of *B. bigemina, B. bovis, B. microti, P. falciparum, T. gondii, T. orientalis, T. parva* and all UniProt/SwissProt protein models. Functional annotations were obtained using InterProScan v5.55-88 with default parameters. For *B. divergens* Rouen 87, gene annotations were transferred to the improved assembly presented here using the PATT pipeline (https://github.com/kjestradag/PATT). Gene models form *B.* MO1 were constructed based on annotations of evolutionarily-related species and further refined using PacBio Iso-seq data specific to *B.* MO1.

### Phylogenetic and phylogenomics analyses

Phylogenomic analysis was conducted using protein sequences from PiroplasmaDB plus *Babesia* sp. MO1 and *B. divergens* Rouen 87 genome annotation from present study, *B. duncani* WA1 [24] and three outgroup genomes, namely *Hepatocystis* sp. (ex *Piliocolobus tephrosceles* 2019), *Plasmodium falciparum* (strain 3D7) and *P. gallinaceum* (strain 8A) from PlasmoDB.

Protein sequences were compared by selecting OrthoMCL groups (supplementary method). Dataset #1 contains the 2,499 orthologous groups having a unique gene per isolate and at least four sequences. Dataset #2 contains the 1,361 orthologous groups from Dataset #1. Each group of orthologous sequences was aligned using the following procedure. First, the orthologous sequences were aligned using Muscle v5.1 [49], with default parameters. Second, the resulting alignment was filtered using HMMCleaner v1.8 [50], with default parameters. Finally, gap-only sequences and gap-only sites were removed using the splitAlignment subprogram of MACSE v2.07 [51]. For each filtered alignment of an orthologous group, we inferred a gene tree by the maximum likelihood criterion using IQ-TREE [52–55] (details given in supplemental material). PhySIC_IST and SuperTriplets require rooted trees, thus we rooted the gene trees by resorting to the outgroup method (see supplemental material for more details). We used three different supertree methods, namely MRP [32], PhySIC_IST [34] and SuperTriplets [35]. The two later require rooted trees as input, thus could only be run on Dataset #2, while MRP could analyze both datasets #1 and #2 (see supplementary data for more details).

We carried out a supermatrix analysis, both on Dataset #1 and #2 by concatenating all alignments of the orthologous groups composing a dataset. We thus obtained a supermatrix of 1,109,333 characters x 21 taxa containing only 34% of missing data for Dataset #1 and 541,931 characters x 18 taxa with 18% missing data for Dataset #2. We then estimated the most likely species tree according to each of these matrices separately, thanks to the IQ-TREE version 2 software. We used the edge-linked partition model to analyze the supermatrix [52, 53], allowing each gene family to have its own evolutionary rate though all families shared the same branch lengths. We obtained branch support with the ultrafast bootstrap [54] by resampling partitions then sites within partitions [56, 57].

### In vitro growth rate determination of *B.* MO1 clones and *B. divergens* clones in different culture media

In vitro cultures of the *B.* MO1 clones B12 and F12 and *B. divergens* Rouen 87 clones H2, and H6 were initiated at 1% parasitemia in human RBCs at 5% hematocrit and sustained in RPMI medium supplemented with either 20% fetal bovine serum or 0.5% albumax. The parasite cultures in the aforementioned media were maintained for four days without subculturing. The respective culture media was replaced daily, and parasite growth was monitored after every 24 h by examination of Giemsa-stained blood smears using a light microscope.

### RNA-seq processing for gene-expression analysis

RNA-seq data were assessed for quality using FastQC v0.11.8. Adapter sequences as well as the first 11 bp of each read were trimmed using Trimmomatic v0.39. Tails of reads were trimmed using Sickle with a Phred base quality threshold of 25, and reads shorter than 18 bp were removed. Reads were then aligned to the *B.* MO1 F12 genome assembly using HISAT2 v2.2.1. Only properly paired reads were retained, with filtering done using Samtools v1.11. Non—uniquely mapped reads were retained due to highly repetitive regions. PCR duplicates were removed with PicardTools MarkDuplicates v2.18.0 (Broad Institute). StringTie v2.2.1 was run with the -e parameter to estimate the abundance of each gene in TPM (transcripts per million).

## ChIP-seq sample preparation

Approximately 20 million *B.* MO1 parasites per sample/per condition were pelleted and crosslinked with formaldehyde, then quenched with glycerine, and followed by a series of washes with PBS. The resulting pellet was resuspended in 1mL nuclear extraction buffer (10 mM HEPES, 10 mM KCl, 0.1 mM EDTA, 0.1 mM EGTA, 1 mM DTT, 0.5 mM AEBSF, 1X Roche protease inhibitor, 1X Roche phosphatase inhibitor) followed by a 30 min incubation on ice. 10% Igepal CA-630 was added to each sample, homogenized by passing through a 26G × 1⁄2 needle and centrifuged at 5,000 rpm to obtain the nuclear pellet. The nuclear pellets were resuspended in shearing buffer (0.1% SDS, 1 mM EDTA, 10 mM Tris-HCl pH 7.5, 1X Roche protease inhibitor, and 1X Roche phosphatase inhibitor) and transferred into 130uL Covaris tubes (PN 520045). Samples were then sonicated using a Covaris S220 (under following settings: 5 min, duty cycle 5%, intensity 140 W, 200 cycles/burst, 6°C) before adding equal volumes of ChIP dilution buffer (30 mM Tris-HCl pH 8, 3 mM EDTA, 0.1% SDS, 30 mM NaCl, 1.8% Triton X-100, 1X protease inhibitor, 1X phosphatase inhibitor). Samples were centrifuged at 13,000 rpm for 10 min at 4°C. For each sample, 13 μL protein A agarose/salmon sperm DNA beads were washed 3 times with ChIP dilution buffer without inhibitors. The washed beads were added to the diluted chromatin for 1 hr at 4°C with agitation to pre-clear the samples. ∼10% of each sample by volume was set aside as input; to the remaining, 2μL of antibodies anti-H3K9me3 (Abcam ab8898), anti-H3K9ac (Diagenode C15410004), or IgG(Abcam ab46540) were added for overnight rotation at 4°C. To each sample, 25 μl of washed protein A agarose/salmon sperm DNA beads with ChIP buffer were blocked with 1 mg/ml BSA for 1 hr at 4°C, re-washed, and added to each sample for 1 hr rotation at 4°C. The bead/antibody/protein complexes were washed a total of 8 times 15 min intervals per wash): twice with low salt buffer (1% SDS,1% Triton X-100, 2 mM EDTA, 20 mM Tris-HCl pH 8, 150 mM NaCl), twice with high salt buffer (1% SDS,1% Triton X-100, 2 mM EDTA, 20 mM Tris-HCl pH 8, 500 mM NaCl), twice with LiCl buffer (0.25 M LiCl, 1% NP-40, 1% Na-deoxycholate,1 mM EDTA, 10 mM Tris-HCl, pH 8.1), and twice with TE (10 mM Tris-HCl pH 8, 1 mM EDTA) buffer. DNA was then eluted from the beads with two 250 μl washes of elution buffer (1% SDS, 0.1 M sodium bicarbonate) and added NaCl (55ul of 5M) to reverse crosslink overnight at 45°C. RNAse A (15 μl of 20 mg/mL) and proteinase K (2 μl 20 mg/mL) were subsequently added to the samples, incubated at 37°C and 45°C, respectively, followed by a DNA extraction via phenol/chloroform and ethanol precipitation. After precipitation, the samples were centrifuged at 13,000 rpm for 30 min at 4°C, forming pelleted DNA, washed with 80% ethanol, re-pelleted, and resuspended the DNA in 50 μl nuclease-free water. The DNA was purified with AMPure XP beads and prepared Illumina sequencing libraries using a KAPA Hyperprep kit (KK8504), followed by the NovaSeq 6000 sequencing platform (Illumina).

### ChIP-seq Analysis

Read quality was analyzed using FastQC (https://www.bioinfor-matics.babraham.ac.uk/projects/fastqc/) and trimmed adapters and low-quality bases using Trimmomatic (http://www.usadellab.org/cms/?page=trimmomatic) and Sickle (https://github.com/najoshi/sickle). Reads were mapped against the *B.* MO1 F12 and B12 assemblies using Bowtie2 v2.4.4 (https://doi.org/10.1038/s41564-023-01360-8) while keeping non-uniquely mapped fragments and retained only correctly paired reads using Samtools (v1.11) (http://samtools.sourceforge.net). PCR duplicates were removed with PicardTools MarkDuplicates v2.18.0 (Broad Institute). To obtain per nucleotide coverage and generate browser tracks, we used BedTools v2.27.1 and custom scripts, normalizing counts by millions of mapped reads. Chromosome tracks were viewed using IGV (Broad Institute). To compare H3K9me3 levels between MGF genes and other genes, read counts for H3K9me3 (and IgG control) were calculated within each gene body using bedtools multicov. Counts were normalized to millions of mapped reads per library and gene length in kb. The background signal from the IgG control was subtracted from H3K9me3 counts, setting negative values to 0. H3K9ac read counts were also generated by bedtools multicov, but including 300 bp upstream of genes as acetylation is often in promoter regions. Heatmaps were generated using normalized H3K9me3, H3K9ac, and RNA-seq TPM counts for each gene to compare histone modifications with gene expression, sorting genes by TPM. The heatmap used log-scaled counts and sorted genes from high to low TPM.

### Data availability

All datasets generated for the current study are accessible in the NCBI/SRA repository under Bioproject PRJNA1032622 (reviewer <u>link</u>). Specifically, the datasets include PacBio HiFi reads (SRA accession number <u>SRR26661633</u>), *B.* MO1 genome, RNA-Seq (SRA accession number <u>SRR26661632</u>), Hi-C reads (SRA accession number <u>SRR26661630</u>, <u>SRR26661631</u>), ChIP-Seq reads (SRA accession number <u>SRR26661627</u>, <u>SRR26661629</u>, <u>SRR26661626</u>, <u>SRR26661628</u>, <u>SRR26661625</u>).

## Acknowledgements

C.B.M.’s research was supported by grants from the National Institutes of Health (AI097218, GM110506, AI123321 and R43AI136118), the Steven and Alexandra Cohen Foundation (Lyme 62 2020), and the Global Lyme Alliance. KLR’s research is supported by the National Institutes of Allergy and Infectious Diseases (R01 AI136511, R01 AI142743-01 and R21 AI142506-01) and the University of California, Riverside (NIFA-Hatch-225935). EM and LMG research is supported by a grant from the Instituto de Salud Carlos III, Spain (PI20CIII-00037).

## Supplementary Figure Legends

**Figure S1.** Chromosomal organization of *Babesia divergens* clinical isolates from France and Spain by PFGE and subsequent Southern blot analyses using a *Plasmodium berghei* telomeric probe. **A**. PFGE (lines 1-5) and Southern-blot (lines 1*-5*) show the number and approximate sizes of chromosomes of *B. divergens* clinical isolates from France. **B**. PFGE (lane 6) and Southern-blot (line 6*) show the number and approximate sizes of chromosomes of the *B. divergens* clinical isolate from Spain. *Schizosaccharomyces cerevisiae* (Sc), *Hansenula wingei* and *Schizosaccharomyces pombe* (Sp) DNA chromosomes were used as DNA markers. The manufacture’s estimates of the sizes of chromosomes are indicated in Megabase pairs on the right and left of Panel A and on the left of panel B [13]. The Table shows epidemiologic and genomic features of the *B. divergens* clinical isolates. [33] [13].

**Figure S2A.** Visualization of the alignment of the *B.* MO1 clone F12 assembly against the Bionano optical map. The green lines represent the optical map molecules, the blue lines represent assembled contigs (1 is ChrI, 2 is Chr2, 3 is Chr3, while the others are unplaced contigs); vertical lines indicate matching positions during the restriction enzyme mapping.

**Figure S2B.** Visualization of the alignment of the *B.* MO1 clone B12 assembly against the Bionano optical map. The green lines represent the optical map molecules, the blue lines represent assembled contigs (1 is Chr I, 2 is Chr II, 3 is Chr III, while the others are unplaced contigs); vertical lines indicate matching positions during the restriction enzyme mapping.

**Figure S3.** Synteny analysis of *B.* MO1 clone B12 (blue), *B.* MO1 clone F12 (orange), and the parental *B.* MO1 (green); gray shaded areas indicated synteny; the length of insertions is annotated; “ITS” indicate the presence of interstitial telomeric sequence in the assembly.

**Figure S4A.** Dot-plot alignment between *B.* MO1 clone F12 and *B.* MO1 clone B12 assembly; the three largest blocks correspond to chromosomes I-III; the dot-plot includes unplaced contigs.

**Figure S4B.** Dot-plot alignment between *B.* MO1 clone F12 and the parental *B.* MO1; the three largest blocks correspond to chromosomes I-III; the dot-plot includes unplaced contigs.

**Figure S5. Phylogenomic analysis. A.** Species phylogeny proposed by Matrix Representation Parsimony (MRP) supertree phylogenomic approaches. Displayed clade support values are estimated by bootstrap on dataset #1. The tree obtained with dataset #2 was identical. All bootstraps were at 100% with dataset #2. *Hepatocystis sp.* (ex *Piliocolobus tephrosceles* 2019), *Plasmodium falciparum* 3D7 and *P. gallinaceum* 8A were taken as outgroup. **B.** Species phylogeny proposed by Super Triplets super tree phylogenomic approaches. The tree was obtained from dataset #2. Displayed clade are confidence value (from 0 to 100) computed by the method with respect to the input trees and then considering only the clades with confidence value above 50. *Hepatocystis sp.* (ex *Piliocolobus tephrosceles* 2019), *Plasmodium falciparum* 3D7 and *P. gallinaceum* 8A were taken as outgroup. **C.** Species phylogeny proposed by super matrix phylogenomic approaches. using Dataset #1’ (see supplementary method)**. D.** Species phylogeny proposed by super matrix phylogenomic approaches based on Dataset #2. All bootstraps were at 100%. *Hepatocystis sp.* (ex *Piliocolobus tephrosceles* 2019), *Plasmodium falciparum* 3D7 and *P. gallinaceum* 8A were taken as outgroup.

**Figure S6. Functional analysis of *Babesia* MO1 gene depending on patristic distances.** Patristic distances were calculated from the trees of dataset #1 for all *Babesia* sp. MO1-*B. divergens* isolates pairs. OUT trees support the position of *Babesia* sp. MO1 as a new species. MIX trees places Babesia sp. MO1 between the two *B. divergens* isolates. **A**. Cumulative distribution of patristic distances among OUT and MIX trees. The X-axis is defined as –log10(patistitic distance). Higher distances are on the left part of the graph. Threshold values (vertical dashed lines) between High, medium, and Low set of genes were the lower and upper quartile of the value that were below 4. Genes with values higher than 4 were considered as non-significant (NS), which means too close to *B. divergens* genes to support any phylogenetic inference. All genes from MIX trees were considered as NS. **B.** GO term enrichment among the four sets of genes. The hypergeometric law was used to evaluate the p-value. GO terms were selected when more than two genes match the term in the subset and p-value was below 0.125. GO terms were ordered from top to bottom by descendant value of the median of patristic distance of all genes matching the terms in a subset. The color intensity is according to the p-value, red being the most significant.

**Figure S7.** Hi-C contact map of *B.* MO1 clone F12; the panels at the bottom are the contact maps for individual chromosomes; green circles/squares indicate the putative location of the centromeres.

**Figure S8.** Hi-C contact map of *B.* MO1 clone B12; the panels at the bottom are the contact maps for individual chromosomes.

**Figure S9.** Hi-C contact map of *B.* divergens Rouen 87; the panels at the bottom are the contact maps for individual chromosomes.

**Figure S10.** GC skew plots for *B.* MO1 clone F12, *B.* MO1 clone B12 and *B.* divergens Rouen 87 obtained using SkewIT.

**Figure S11.** Sequence alignment of DHFR-TS from different *Babesia* and *Plasmodium* species.

**Figure S12.** 3D genome structures of *B. divergens* Rouen 87 derived from the contact map interactions (Fig. S9). Chromosomes one, two, and three correspond to green, pink, and blue sections respectively. Dark green and grey represent the telomeric regions and centromeres.

**Figure S13.** Evolution of *B. MO1* and *B. divergens*. **A.** Phylogenetic tree constructed using 18S rRNA from different apicomplexan species, including *T. gondii*, *P. falciparum*, *B. duncani*, *B. microti*, *B. bovis*, *B. ovata*, *B. bigemina*, *B. divergens*, *B.* MO1 and *T. parva*. **B.** Phylogenetic tree constructed based on mitochondrial genome sequences from different *Babesia* species.

## Supplementary Methods

### Cloning of *B. MO1* isolate

*B. MO1* in vitro culture was initiated in A^+^ human RBCs in DMEM/F12 medium at 0.5% parasitemia and 5% hematocrit (HC). The parasite culture was allowed to grow for four days and the parasitemia was measured by Giemsa-stained blood smears. The culture was subjected to serial dilution to obtain 30 parasites in 20 ml (5% HC) and 200 μl of this parasite suspension was plated per well in a 96-well plate. The culture medium of the cloning plate was replaced with fresh medium every 3^rd^ day for 21 days. On day 22, SYBR Green-I assay was performed to determine the parasite positive wells of the cloning plate. Briefly, 25 μl of culture per well from the cloning plate was transferred to a black bottom 96-well plate (Stellar Scientific, IP-DP35F-96-BLK) and mixed with 25 μl of SYBR Green-I lysis buffer (20 mM Tris, pH 7.4, 5 mM EDTA, 0.008% saponin, 0.08% Triton X-100 and 1X SYBR Green-I (Molecular Probes, 10,000X solution in DMSO, Eugene, OR, USA)) and incubated for 30 min in dark at 37°C. In addition, the uninfected human RBCs (5% HC, 25 μl volume) were used as a negative control. Following the incubation, the SYBR Green-I measurement was performed on BioTek Synergy MX fluorescence plate reader with an excitation of 497 nm and emission of 520 nm. The readings from uninfected human RBCs were used as background and subtracted from the readings of the cloning plate wells in order to determine wells positive for parasites (higher SYBR Green-I readings in comparison to the negative control). Following identification of parasite positive wells using SYBR Green-I assay, the same wells were used to prepare smears for Giemsa staining and presence of parasites was confirmed using light microscopy. Six clones from parasite positive the 96-well plate were picked and expanded to 1 ml cultures and allowed to grow to 2% parasitemia before expanding them to 5 ml cultures. Two of the six clones (*B. MO1* clone B12 and clone F12) were used in this study.

### DNA preparation for Oxford Nanopore and Illumina sequencing for *B. divergens* Rouen 87

Genomic DNA (gDNA) was isolated from asynchronous *B. divergens in vitro* cultures with 40% of parasitemia. The gDNA was prepared using pellets of infected RBCs. Pellets were lysed with 0.15% Saponin (Sigma-Aldrich) for 30 minutes and centrifuge at 2000 x g and 4°C for 10 minutes. The final pellets were incubated in lysis buffer (0.1 M NaCl, 50 mM Tris-HCl, pH 7.5, 1 mM EDTA, sodium dodecyl sulfate [SDS; 0.5% by volume], and 100 μg ml^-1^ of proteinase K (Sigma-Aldrich) for 16 h at 56°C. Nucleic acid was recovered by phenol-chloroform extraction, followed by ethanol precipitation. RNA was removed by RNase digestion (Roche Diagnostic GmbH, Germany) and DNA was subjected to a further round of phenol-chloroform extraction and ethanol precipitation.

### DNA preparation for Bionano Optical Map for *B. MO1*

*B. MO1* was cultured in vitro in human RBCs to attain a parasitemia of 8-10% at 5% haematocrit (total 100 ml). The parasite pellet was generated by centrifuging the cultures at 500 x g and used to isolate ultra-high molecular weight (HMW) genomic DNA for use in genomic optical mapping (Histogenetics) using the Bionano Prep Blood and Cell Culture DNA Isolation kit (Bionano Genomics, 80004). The DNA was quantified using Qubit dsDNA BR Assay kit. Around 0.8g of HMW DNA was labelled using the Bionano Prep direct label and stain method (Bionano Genomics, 80005) and loaded onto a flow cell to run on the Saphyr optical mapping system (Bionano Genomics). Around 1.2 Gb of data were generated per run. Raw optical mapping of molecules in the form of BNX files were run through a preliminary bioinformatics pipeline that filtered out molecules less than 150 kb in size and less than 9 motifs per molecule to generate a *de novo* assembly of the genome maps.

### Genome Sequencing and Assembly of *B. MO1* isolate F12 and B12

DNA for clones F12 and B12 were sequenced at the Yale Center for Genome Analysis using PacBio HiFi (CCS). HiFi reads for clone F12 totaled 31.2 B bases, which translated to a ∼2600x coverage of the *B. MO1* genome (assuming a genome of 12Mb). Given the abundance of sequencing data, hifiasm v0.19.6 [32] and HiCanu v.2.2 [24] were tested on (1) the entire 2600x-coverage data, (2) the 250 thousand longest HiFi reads (511x coverage, average read length = 21,043 bp), and (3) the 100 thousand longest HiFi reads (228x coverage, average read length = 27,362 bp). These six assemblies were aligned to the Bionano optical map using Bionano RefAligner Solve v3.7 to detect possible mis-joins. Based on assembly statistics, comparison with the optical map and BUSCO completeness, it was determined that the best assembly of clone F12 was obtained using hifiasm on the 100 thousand longest HiFi reads. This assembly was used as the reference *B. MO1* genome in the rest of this study. HiFi reads for clone B12 totaled 33.8 B bases, which translated to a ∼2800x coverage of the *B. MO1* genome (assuming a genome of 12Mb). The same assembly strategy used for F12 was used for clone B12. The best assembly of clone B12 was obtained again using hifiasm on the 100 thousand longest HiFi reads (about 200x coverage, average read length = 24,041 bp).

### Genome Sequencing and Assembly of *B. divergens* Rouen 87

DNA from a *B. divergens* culture was used for Oxford Nanopore sequencing. Sequencing libraries were prepared using the SQK-LSK109 kit with a 1µg of DNA input following the vendor’s protocol. Sequencing was performed using a MinION flow cell (v9.4). The base-calling was carried out using the software Guppy v4.0.14 with default parameters and a high accuracy error model (dna_r9.4.1_450bps_hac.cfg). A *de novo* assembly was performed using Oxford Nanopore long reads and Canu v.1.9 [24] assembler with default parameters. This assembly was corrected using Illumina reads from the already previous *B. divergens* assembly [19] and three iterations of Pilon v.1.23 [33].

### PacBio IsoSeq processing

PacBio IsoSeq data was mapped to the *B. MO1* genome using Minimap2 with options ‘splice:hq -uf –secondary=no -C5’. The resulting alignments were fed into the PacBio cDNA_Cupcake pipeline (https://github.com/Magdoll/cDNA_Cupcake) using the script ‘collapse_isoforms_by_sam.py’ to obtain non-redundant transcript isoforms. The isoform sequences were used in the gene finding pipeline below.

### Comparative genomics

Comparative genomics between different species of *Babesia* was performed by running OrthoMCL on the genome data obtained from PiroplasmaDB and PlasmoDB release 58. *Babesia bigemina* strain BOND*, Babesia bovis* T2Bo, *Babesia divergens* strain 1802A, *Babesia duncani* strain WA1, *Babesia microti* strain RI, *Babesia ovata* strain Miyake, *Babesia sp. Xinjiang Xinjiang*, and *Theileria parva* strain Muguga genomes were used in this analysis. OrthoMCL was run on these eight species, as well as the newly assembled genomes of *B. divergens* Rouen 87 and *B.* MO1. The UpSet plot was generated using R.

The pairwise comparisons between the genome of *Babesia* species was performed. First, the synteny between assemblies using the web server Genies (http://dgenies.toulouse.inra.fr/) with the Minimap2 aligner was calculated. The average nucleotide identity (ANI) between all genome pairs was calculated with PyANI v.0.2.10 (https://github.com/widdowquinn/pyani). Synteny circos plots in Fig 3 were obtained using mummer2circos v1.4.2 (https://github.com/ metagenlab/mummer2circos) that uses the promer algorithm in conjunction with CIRCOS. CIRCOS plots in Fig 4 were generated using the circoletto.pl script v.07.09.16 that uses CIRCOS v2.43.0 underneath. The used options for circoletto were: --out_size 2000 --e_value 1e-3 --untangling_off (https://github.com/infspiredBAT/Circoletto). We obtained orthologous proteins between *B. divergens* Rouen and B. MO1, *B. microti* and *B. bovis*, we used the ProteinOrtho v.6.0.24 software using default parameters and proteins from each genome. The genome from the mitochondrion and apicoplast organelles for *Babesia divergens* Rouen and *B.* MO1 were compared against other species (*B. ovata, B. microti, B. bovis* and *B. bigemina*) by performing a multiple alignment with MAFFT v.7.453 with the following parameters: --reorder --maxiterate 1000 --threadit 0 --retree 1. A phylogenetic tree was generated with a maximum likelihood approach by using first jmodeltest-2.1.10 to select the best tree model and then PhyML version 3.3.3:3.3.20190909-1 to generate the tree.

Gene localization plots in Fig 5 were produced using our tool GFViewer (https://github.com/sakshar/gene-localization-tool).

GC-skew plots in Fig S13 were obtained using SkewIT (https://jenniferlu717.shinyapps.io/SkewIT/) [PMID: 33275607]

### Phylogenetic analyses

To infer the species phylogeny, a phylogenomic analysis was conducted using protein sequences from PiroplasmaDB plus *Babesia sp.* MO1 and *B. divergens* Rouen 1987 genome annotation from present study, *B. duncani* WA1 [27] and three outgroup genomes, namely *Hepatocystis s*p. (ex *Piliocolobus tephrosceles* 2019), *Plasmodium falciparum* (strain 3D7) and *P. gallinaceum* (strain 8A) from PlasmoDB. Pseudogenes and genes encoding peptides below 100 amino acids were removed. CH-HIT was used to removed duplicated genes with following for loop for f in *.fasta; do b=$(basename $f .fasta); ../../BABESIA_2022/soft/CDHIT/cd-hit-v4.8.1-2019-0228/cd-hit -i $f -o ../CDHIT_results/${b}_noDup.fasta -c 1.00 -t 1 > ../CDHIT_results/${b}_noDup.log;done

For the analysis of orthology groups, *B. sp.* MO1*, B. divergens* Rouen 1987 and *B. duncani* genes were assigned to OrthoMCL (https://OrthoMCL.org) groups using the orthology assignment tool available through the VEuPathDB (https://VEuPathDB.org) Galaxy workspace. Proteins in FASTA format were assigned to groups based on the OG6r15 BLAST database using the default settings. Output files generated by the OrthoMCL pipeline included a mapping file between gene IDs and OrthoMCL v.6 group IDs. VEuPathDB resources including PlasmoDB.org and PiroplasmaDB.org provided OrthoMCL v.6 group IDs. A matrix containing the number of genes per OrthoMCL group was generated with a custom R script.

Protein sequences were compared by selecting OrthoMCL groups. Each group of orthologous sequences was aligned using the following procedure. First, the orthologous sequences were aligned using Muscle v5.1 [34], with default parameters. Second, the resulting alignment was filtered using HMMCleaner v1.8 [35], with default parameters. Finally, gap-only sequences and gap-only sites were removed using the splitAlignment subprogram of MACSE v2.07 [36].

For data set generation, we selected only a subset of these alignments for phylogenomic analysis. Indeed, inferring a species tree from families containing both orthologous and paralogous sequences is error prone. Thus, we only considered families with at most one sequence per taxa, maximizing the probability to consider only orthologous sequences. We restricted ourselves to gene families spanning at least four taxa (there is only one possible unrooted tree topology for three taxa). Phylogenomic inference was done using supermatrix and supertree methods. Some supertree methods require rooted trees as input. Overall, we considered two datasets: Dataset #1 contains the 2,499 orthologous groups having a unique gene per isolate and at least four sequences. Dataset #2 contains the 1,361 orthologous groups from Dataset #1 that additionally contained at least one outgroup sequence and such that the outgroup sequences were monophyletic in the corresponding gene tree (when several outgroup sequences were present).

A tree showing has been inferred by maximum likelihood through the IQ-TREE version 2 software for each gene family [37–40] with the command:

iqtree2 -s OG6_100089_filtered.aln --seqtype AA -b 100 -mset LG,WAG,JTT,Blosum62 - cmax 4 --prefix OG6_100089_iqtree --quiet

where OG6_100089 is the gene family considered here.

The matrices of patristic distances (distance from one leaf to another in a phylogeny) was calculated for our 2499 trees with the following command:

for c in $(cat ../cog.list); do nw_distance -n -m m ALIPHY_DETAILS/${c}/${c}_iqtree.treefile > patristiDistances/${c}.pdist; done

The maximum likelihood inference detailed above gave unrooted gene trees. We rooted each of them by placing the root node on the branch separating the outgroup taxa from the other ones. The outgroups in this analysis are *Hepatocystis* sp., *Plasmodium falciparum* 3D7 and *P. gallinaceum* 8A. When a gene family contained no outgroup, it could not be rooted.

The rooting was performed by the version 0.1.3 of the bpp-reroot utility from Bio++ (Dutheil et al 2006). For instance, for the OG6r15_117499 orthologous group we used the following command:

./bppReRoot input.list.file=OG6r15_117499_iqtree.treefile outgroups.file=outgroup.txt output.trees.file= OG6r15_117499.bppReRoot.nwk print.option=false

Graphic representation was performed using ggplot2 in R.

PhySIC_IST and SuperTriplet require rooted trees, thus we rooted the gene trees by resorting to the outgroup method (see supplemental material for more details). Here outgroup taxa are the two *Plasmodium* isolates together with *Hepatocystis* sp. sequences.

We inferred a piroplasma phylogeny from datasets #1 and #2. We performed both a supermatrix and a supertree analysis. We used three different supertree methods: MRP [26], PhySIC_IST [28] and SuperTriplets [29]. The two latter require rooted trees as input, thus could only be run on Dataset #2, while MRP could analyze both datasets #1 and #2.

The analysis with MRP method was conducted by using the BuM program, available online at http://nuvem.ufabc.edu.br/bum. We obtained a binary character matrix encoding the source trees for datasets #1 and #2 separately after trimming all branch lengths and clade support values according to the program manual. For both datasets we produced a most parsimonious tree for the character matrix by the TnT software. The analyzing script asked TnT to perform an exact search of the most parsimonious tree, which is feasible for such a small number of taxa. Below is the precise script used for analyzing Dataset #1:

log ds1_optimal.log;

mxram 1000;

nstates NOGAPS;

taxname=;

p ds1_treefiles_topo.ss;

hold 1000;

ienum;

export - ds1_optimal_MRP.tre;

quit;

The computations on datasets #1 and #2 ended up proposing only one single most parsimonious tree (Figure 1 in main paper). We then relaunched the parsimony analysis of the matrices, this time asking for bootstrap support, using the following script:

log ds1_boot.log;

mxram 1000;

nstates NOGAPS;

taxname=;

p all_OG_1Copy_4spe_bpp_could_root.ss;

hold 1000;

rseed 0;

collapse 0;

ienum;

export - ds1_initial_intensive.best;

resample boot rep 1000 freq savetrees [mult=rep 1 hold 1];

export - ds1.intensive.boottrees;

log/; quit;

PhySIC_IST offers the possibility to detect and correct outlier clades among the source trees. We can mainly set two parameters for this method: i) a confidence threshold b above which the clades of the source trees should be considered (in our case, this confidence value was inferred for each source tree by bootstrap from the alignment of the corresponding orthologous group); ii) a correction threshold c of strictness in correcting outlier clades form the source trees.

The analysis with the PhySIC_IST method was conducted for a large number of combinations of the STC (-c flag) and confidence (-b flag) parameters: from 0 to 1 varying by 0.1. The confidence support allows to account only for branches of the input trees having a support (e.g., bootstrap) above a given threshold. The STC parameter allows to change the behavior of the method from a purely optimization method (lower values of STC) to a strict consensus method (STC set to 1.0). More precisely, increasing STC (up to 100%) allows a smaller and smaller minority of trees to put a veto to proposed clades that contradict some of their triplets. Hence, ultimately, when set at 1.0, for any clade in the proposed supertree, all triplets induced by this clade must be present or induced by the input trees and, moreover, not contradicted by any of them. A typical command line to run PhySIC_IST was:

./PhySIC_IST-newMac.v1.1.0 -s ds2.tre -b $B -c $C -o physicist-b${B}-c${C}.tre -f newForest-b${B}-c${C}.tre > phys-b${B}-c${C}.out

where $B and $C are values for the confidence and STC parameters respectively, ds2.tre contains the gene trees of dataset #2, newForest-b${B}-c${C}.tre is the set of input trees modified to only keep branches with a threshold at least $B

The analysis with the superTriplets method was conducted as following:

java -jar -Xmx600m SuperTriplets_v1.1.jar rootedTress.tre superTripletSupportedClades.tre and lead to the binary phylogeny. The reliability of each clade is based on the percentage of triplets of the input trees in agreement/disagreement with the clade (a triplet is a subtree connecting three given leaves. Any rooted input tree on n leaves can be equivalently represented by its set of O(n3) triplets). Note that superTriplets branch supports are more conservative than traditional bootstrap values. They mostly reflect the percentage of gene trees supporting the clade (independently of the number of considered gene trees).

We carried out a supermatrix analysis, both on Dataset #1 and #2 by concatenating all alignments of the orthologous groups composing a dataset. We thus obtained a supermatrix of 1,109,333 characters x 21 taxa containing only 34% of missing data for Dataset #1 and 541,931 characters x 21 taxa with 18% missing data for Dataset #2. We then estimated the most likely species tree according to each of these matrices separately, thanks to the IQ-TREE version 2 software. We used the edge-linked partition model to analyze the supermatrix [37, 38], allowing each gene family to have its own evolutionary rate though all families shared the same branch lengths. We obtained branch support with the ultrafast bootstrap [39] by resampling partitions then sites within partitions [41, 42].

We met a technical problem with the IQ-TREE method when analyzing Dataset #1, as distances between some taxa were too important (>3), which stopped the program at an intermediary inference step. To tackle the problem of studying too distant taxa, we temporarily removed the three outgroups (*Hepatocystis* sp., *Plasmodium falciparum* 3D7 and *P. gallinaceum* 8A) from the 2499 alignments, as *B. microti* was consistently found at the root of remaining taxa in the previous analyses (see above) and this could be used to root the obtained phylogeny. We discarded the alignments where less than 3 taxa remained. We thus obtained a data set (denoted #1’) of 2,381 alignments on 18 taxa.

Tree samples:

- MRP

From dataset #1 ((BdunW,((CfelW,(TequW,((ToriS,(ToriF,ToriG)),(TannA,TparM)))),(BmicR,(Hpil2,(Pgal8, Pf3D7))))),(((BxinX,(BoviS,BbovF)),(BcabD,(BovaM,BbigB))),(Bmo1F,(Bdiv1,BdivR))));

With PhyML Bootstrap

(ToriF:0.01902491,ToriG:0.01481576,(ToriS:0.00000001,((TannA:0.00142389,TparM:0.001 02794)100:0.07953690,(TequW:0.01282136,(CfelW:0.01651942,((BdunW:0.00609445,((B moGF:0.00000001,(BdivR:0.00284104,BdivG:0.00689378)100:0.05757303)100:0.10101826,((BxinX:0.00215897,(BbovF:0.01250604,BoviS:0.00651609)100:0.06222349)100:0.095625 10,(BcabD:0.01180556,(BbigB:0.00126298,BovaM:0.00108992)100:0.11136155)100:0.030 87336)100:0.06988525)100:0.13588391)100:0.04311891,(BmicR:0.00442233,(Hpil2:0.0000 0001,(Pgal8:0.01592997,Pf3D7:0.01586667)100:0.04094167)100:0.13248621)100:0.082298 58)100:0.07498309)100:0.03831524)100:0.11464958)100:0.10066643)100:0.03154716);

- PhySIC_IST with confident factor from dataset 2
(((Hpil2,(Pgal8,Pf3D7)55.4)100,(BmicR,((CfelW,(TequW,((TannA,TparM)96.7,(ToriS,(Tori G,ToriF)49.4)97)92.6)40.5)60.2,(BdunW,((Bmo1F,(Bdiv1,BdivR)83)99,((BcabD,(BovaM,B bigB)96.4)36.5,(BxinX,(BoviS,BbovF)73.5)75.8)65.4)95.4)50.7)80.2)100):0.0000000000;
- SuperTriplets with support
(((Pf3D7,Pgal8)55,Hpil2)100,((((((BdivR,Bdiv1)83,Bmo1F)99,((BovaM,BbigB)98,BcabD,((BoviS,BbovF)75,BxinX)82)73)97,BdunW)58,(TequW,((TparM,TannA)97,(ToriG,ToriF,Tor iS)98)96,CfelW)74)83,BmicR)100);
- Super matrix with dataset #1

(ToriS:0.0706113988,(((((((((BoviS:0.2031976164,BbovF:0.3980474090):0.1147599795,Bxi nX:0.2494624721):0.1472209562,((BovaM:0.0780112597,BbigB:0.0857584247):0.2816494 470,BcabD:0.3099744240):0.0454836912):0.1709173511,(Bmo1F:0.0180586829,(Bdiv1:0.0 015619949,BdivR:0.0008510062):0.0103423142):0.4030948603):0.7437210735,BdunW:1.0 050350068):0.1732543377,BmicR:2.5743476758):0.2769205668,CfelW:0.6911774661):0.1 108373463,TequW:0.5273225306):0.6369920638,(TannA:0.0966684769,TparM:0.0926800 225):0.3383463649):0.3648904977,(ToriF:0.0550107601,ToriG:0.0451767588):0.02127602 59);

- Super matrix with dataset #2

(Pgal8:0.1064314407,(((((((TannA:0.0674276368,TparM:0.0629092051)100:0.2144271461,(ToriS:0.0448211435,(ToriF:0.0360463184,ToriG:0.0299960787)100:0.0143531254)100:0.2 287617944)100:0.3864559684,TequW:0.3322368963)100:0.0698171744,CfelW:0.43284095 90)100:0.1623446465,(((Bmo1F:0.0126225224,(Bdiv1:0.0011523844,BdivR:0.0008050027) 100:0.0074767120)100:0.2490952175,(((BovaM:0.0519324115,BbigB:0.0557209125)100:0. 1743791334,BcabD:0.1961292999)100:0.0311203505,(BxinX:0.1618722380,(BbovF:0.2494 914854,BoviS:0.1323962495)100:0.0718957894)100:0.0929663209)100:0.1099934141)100: 0.4411543291,BdunW:0.6223490019)100:0.1157227747)100:0.5131302484,BmicR:1.01525 66998)100:1.8324573530,Hpil2:0.1903734173)100:0.0676408189,Pf3D7:0.1325278584);

### *In vitro* drug efficacy

The inhibitory effect of currently used anti-babesial drugs including atovaquone, clindamycin, azithromycin, quinine and an antifolate drug WR99210 on the intra-erythrocytic development of *B. MO1* parental isolates and clones B12 and F12 were tested and IC_50_ determination was performed using a previously reported protocol. Briefly, *B. MO1* parental isolate as well as two clones were cultured *in vitro* in human RBCs at 5% hematocrit (HC) in complete DMEM/F12 medium (Lonza, BE04-687F/U1). The parasite cultures (0.5% parasitemia, 5% HC in complete DMEM/F12 medium) were treated with decreasing concentrations of the compound of interest in a 96-well plate for 72 h. Following this, the parasitemia determination was performed using SYBR Green-I assay [31]. Briefly, 100 μl of the drug treated, or control parasite cultures were mixed with 100 μl of lysis buffer (0.008% saponin, 0.08% Triton-X-100, 20 mM Tris-HCl (pH = 7.5) and 5 mM EDTA) containing SYBR Green-I (0.01%) and incubated at 37°C for 1h in the dark. The fluorescence was measured at 480nm (excitation) and 540 nm (emission) by using a BioTek Synergy™ Mx Microplate Reader. The background fluorescence (uninfected RBCs in complete DMEM/F12 medium) was subtracted from each concentration and 50% inhibitory concentration (IC_50_) of the drug was determined by plotting sigmoidal dose-response curve fitting with drug concentration and percent parasite growth in the Graph Pad prism 9.4.1 from three independent experiments performed in triplicates. Data are shown as mean ± SD.

### DNA preparation for PacBio sequencing

In vitro cultures of *B. MO1* clones B12 and F12 were initiated in human RBCs at 1% parasitemia, 5% HC (50 ml each) and cultured to attain 10% parasitemia. The cultures were harvested, and genomic DNA was isolated from both the clones using DNasy Blood and Tissue kit (Qiagen, Cat. No. 69506), The concentration determination and quality control was assessed using nanodrop and qubit, respectively. DNA integrity was determined using Blue Pippin pulse gel and following this, the DNA was used for library preparation using Pacific Biosciences SMRTbell Express template Prep Kit 2.0 (Cat. No. PN: 100-938-900) according to the manufacturer’s instructions. Loading concentration and proper stoichiometric measurements were determined using the Pacific Biosciences Smart Link software. Following this, the gDNA library was annealed to the Pacific Biosciences V5 primer for 1h at 20°C. The annealed library was then bound to polymerase using Pacific Biosciences Polymerase 2.2 for 1-4h at 30°C and was loaded on to the Sequel II Instrument as an adaptive sequencing run. At least one smart cell was sequenced for each genomic DNA library with a movie time of 30h and a pre-extension of 2h. After the DNA library sequencing was complete, the loading metrics were evaluated by mean read length, polymerase read length, data yield and P1 values to ensure the sample ran as expected and data had met Yale’s gold standards (polymerase read length between 50-60kb, data yield (HiFi) around 2-4 million reads of total 10-20Gb, and P1 between 60-70%).

### DNA preparation for Hi-C

*In vitro* cultures of *B. MO1* clones B12 and F12 were initiated in human RBCs at 1% parasitemia, 5% HC (100 mL) and cultured to attain 10% parasitemia. The cultures were centrifuged, and the parasite pellets were cross-linked with 1.25% formaldehyde for 25 min at 37°C. Cross-linking reaction was quenched by the addition of 150mM (final concentration) glycine and incubation for 15 min at 37°C followed by a 15 min incubation at 4°C. This was followed by the lysis of parasite pellets by resuspension in lysis buffer (10 mM Tris-HCl, pH 8.0, 10 mM NaCl, 2 mM 4-(2-aminoethyl) benzenesulfonyl fluoride HCl (AEBSF), 0.25% Igepal CA-360 (v/v), and EDTA-free protease inhibitor cocktail (Roche)) and incubation for 30 min on ice. Nuclei were isolated after homogenization by 15 needle passages. *In situ* Hi-C protocol was conducted as described by Rao and colleagues [32]. Briefly, 0.5% sodium dodecyl sulfate (SDS) was used to permeabilize the nuclei. Subsequently, the DNA was digested using 100 units of Mbol (NEB), the ends of restriction fragments were filled using biotinylated nucleotides and ligated using T4 DNA ligase (NEB). After reversal of crosslinks, ligated DNA was purified and sheared to a length of ∼300-500 bp using the Covaris ultrasonicator S220 (settings: 10% duty factor, 200 cycles per burst and a peak incident power of 140). Ligated fragments were pulled down using streptavidin beads (Invitrogen) and prepped for Illumina sequencing by subsequent end-repair, addition of A-overhangs and adapter ligation. Libraries were amplified for a total of 12 PCR cycles (45 sec at 98°C, 12 cycles of 15 sec at 98°C, 30 sec at 55°C, 30 sec at 62°C and a final extension of 5 min at 62°C) and sequenced with the NOVASeq platform (Illumina), generating 100 bp paired-end sequence reads at the UCSD core facility.

### RNA preparation for Illumina RNA-seq

*B. MO1* clones B12 and F12 were cultured to a parasitemia of 8% at 5% HC (10mL culture volume per clone). Total RNA was isolated from clones B12 and F12 using five volumes of Trizol LS Reagent (Life Technologies, Carlsbad, CA, USA) and following manufacturer’s instructions. Total RNA was subjected to DNA-free DNA removal kit (ThermoFisher; AM1906) for removal of contaminating DNA. Following this, mRNA was purified from total RNA using NEBNext Poly(A) mRNA Magnetic Isolation Module (NEB, E7490S), and RNA-seq library was constructed using NEBNext Ultra II RNA-library preparation kit (NEB, E7770S) according to the manufacturer’s instructions. The RNA-libraries were amplified for 15 PCR cycles (45s at 98°C followed by 15 cycles of [15s at 98°C, 30s at 55°C, 30s at 62°C], 5 min 62°C). Next, the libraries were sequenced at 150 bp paired-end sequenced on the Illumina Novaseq platform (Illumina, San Diego, CA) at the UCSD and Yale core facility.

### Oxford Nanopore Sequencing

DNA from *B. divergens* Rouen 87 and *Babesia* MO1 was not sheared and was used directly from purification for library construction. An ONT genomic DNA library was prepared by Ligation using the kit SQK-LSK109 following the vendor’s protocol. A size-selection step was done at the last purification step after adapter ligation using Large Fragment Buffer (LFB) to wash AMpure XP beads, just before loading the library in the MinION R9.4.1 flow-cell. Base calling was performed with the Guppy software requesting High Accuracy Calling on a laptop with Graphic Processing Units (GPU’s).

### DNA preparation for Bionano optical map

Exactly 3 ml packed frozen pellets of *B. MO1* in human RBCs were used to isolate ultra-high molecular weight (uHMW) genomic DNA for use in genomic optical mapping by Histogenetics (Ossining, NY) using the Bionano Prep™ Blood and Cell Culture DNA Isolation Kit (Bionano Genomics, cat No. 80004). Following this, DNA was quantified using Qubit™ dsDNA BR Assay Kit. A total of 0.75 ug of HMW DNA was then labeled using the Bionano Prep direct label and stain (DLS) method (Bionano Genomics, cat No. 80005) and loaded onto a flow cell to run on the Saphyr optical mapping system (Bionano Genomics). Approximately 1,177 Gb of data was generated per run. Raw optical mapping molecules in the form of BNX files were run through a preliminary bioinformatic pipeline that filtered out molecules less than 150 kb in size with and less than 9 motifs per molecule to generate a *de novo* assembly of the genome maps.

### Illumina sequencing

Extracted DNA passed standard quantity, quality and purity assessments via determination of the 260/280nm for values of 1.7-2.0, and 260/230 absorbance ratios for values ≥ and 1% agarose gel electrophoresis to ensure that the gDNA is neither degraded nor displays RNA contamination. The library preparation started with 0.5ug of well quantified gDNA and underwent enzymatic fragmentation, end-repair and “A” base in a single reaction using Lotus DNA Library Prep kit (IDT, Part#10001074). The adapters with appropriate dual multiplexing indices, xGen UDI-UMI Adapters (IDT, Part #10005903), were ligated to the ends of the DNA fragments for hybridization to the flow-cell for cluster generation. Size of the final library construct was determined on Caliper LabChip GXsystem and quantification was performed by qPCR SYBR Green reactions with a set of DNA standards using the Kapa Library Quantification Kit (KAPA Biosystems, Part#KK4854). For sequencing, the sample concentrations were normalized to 2nM and loaded onto Illumina NovaSeq 6000 S4 flow cells at a concentration that yields the requested number of passing filter data per lane. Samples were sequenced using 151 bp paired-end sequencing reads according to Illumina protocols.

### PacBio Iso-Seq library preparation and sequencing of *Babesia* MO1

TRIzol reagent (Life Technologies, Carlsbad, CA, USA, No. 15596–026) was used to isolate total RNA from 100 ml *in vitro* culture of *B.* MO1 (15% parasitemia and 5% hematocrit) according to the manufacturer’s protocol. 1 µg of total RNA was used for the synthesis and amplification of cDNA using a combination of NEBNext Single Cell/Low Input cDNA Synthesis & Amplification module (Cat. No. E6421S), NEBNext High-Fidelity 2X PCR Master Mix (Cat. No. M0541S), Iso-Seq Express Oligo Kit (Cat. No. PN 101-737-500), and elution buffer (Cat. No. PN 101-633-500). SMRTbell libraries were constructed according to the Iso-Seq Express Template Protocol (Pacific Biosciences). Primer annealing and polymerase binding were performed following the SMRT Link v8.0 Sample Setup instructions and 90 pM of the SMRTbell templates were loaded for sequencing. One SMRT Cell 8M was used for each sample and sequencing was performed using the Sequel II system.

### Illumina RNA-Seq library preparation and sequencing of *B. divergens* Rouen 87

Free merozoites and intraerythrocytic parasites were collected from two highly parasitized independent asynchronous *B*. *divergens* cultures, 75 ml each at parasitemias of 40% Total RNA from *B. divergens* free merozoites and intraerythrocytic parasites was prepared using Trizol LS Reagent (Life Technologies, Carlsbad, CA, USA, No. 15596–026) and chloroform extraction. Libraries were prepared using the Illumina Kit (Illumina) following the manufacturer’s protocol. High quality RNA samples from three biological replicates of free merozoites and from intraerythrocytic stages were used to prepare three independent libraries for each stage. The libraries were sequenced using the Illumina HiSeq platform with a paired-end configuration.

### PacBio HiFi sequencing

Genomic DNA was isolated from 100 ml *in vitro* culture of *B.* MO1 (15% parasitemia and 5% hematocrit) using DNasy Blood and Tissue kit (Qiagen; Cat. No. 69506), and quality control along with concentration determination was performed by using nanodrop and qubit. DNA integrity was evaluated using Blue Pippin pulse gel and the DNA was then used for library preparation using Pacific Biosciences SMRTbell Express template Prep Kit 2.0 (Cat. No. PN: 100-938-900) according to the manufacturer’s instructions. The Pacific Biosciences Smart Link software was used to determine loading concentration and proper stoichiometric measurements. The gDNA library was then annealed to the Pacific Biosciences V5 primer for 1h at 20°C. The annealed library was then bound to polymerase using Pacific Biosciences Polymerase 2.2 for 1-4h at 30°C and was loaded on to the Sequell II Instrument as an adaptive sequencing run. At least one smart cell was sequenced for each genomic DNA library with a movie time of 30h and a pre-extension of 2h. After the DNA library sequencing was complete, the loading metrics were evaluated by mean read length, polymerase read length, data yield and P1 values to ensure the sample ran as expected and data had met Yale’s gold standards.

### Hi-C data processing

Illumina reads were mapped using BWA MEM 0.7.17 [57] Contact maps were produced using HiC-Explorer v3.7.2 [43].

### Three-dimensional modeling

Three-dimensional coordinate matrices were generated from the HiCexplorer output matrices using PASTIS [44]. The coordinate matrices were then converted to PDB format and visualized as 3D chromatin models in ChimeraX [58] and 10-kb bins containing telomeres and the approximate location of centromeres were highlighted.

## Pulse field gel electrophoresis (PFGE)

Cultures of *B. divergens* MO1 and *B.* MO1 clones (B12, H1, F12, H6, A3 and F1), *B. divergens* Rouen 87 and *B. divergens* clones (H2, H6, C1, C7, A6 and H10) and the *B. divergens* clinical isolate from Spain were centrifuged at 1.300 x g for 5 min to yield pellets containing intact cells. Pellets, were embedded in 1% (w/v) SeaKem Gold Agarose (Lonza, Rockland, ME, USA) to an approximately concentration of 1×10^8^ infected RBCs/ml. The resultant agarose plugs were incubated in lysis solution (100mM EDTA, pH8.0, 0.2% sodium deoxycholate, 1% sodium lauryl sarcosine) supplemented with 1 mg/ml of proteinase K (Thermo Fisher Scientific, Vilnius, Lithuania) for 24 h at 50°C. Finally, plugs were washed 4 times for 30 min each in wash buffer (20 mM Tris, pH 8.0, 50 mM EDTA). Intact chromosomes were separated on a 0.8% Megabase Agarose gel (Bio-Rad Labs Inc., Hercules, CA, USA) in 1X TAE buffer chilled at 14°C for 48 h for *B. divergens* MO1 and *B. MO1* clones and 72 h for *B. divergens* Rouen 87, *B. divergens* clones and the *B. divergens* clinical isolate from Spain on a CHEF MapperTM XA pulsed field electrophoresis system (Bio-Rad). The switch time was 20 min-40 min-23 sec at 2V/cm with an include angle of 106L. The agarose gel was stained with GelRed (Biotium, Fremont, CA, USA) and visualized under ultraviolet transilluminator.

## Southern Blot Analysis

Telomeric ends of *B. divergens* clinical isolate form Spain chromosomes were analyzed by Southern Blot using a nucleotide repeat sequence (CCCTGAACCCTAAA) of the telomeric ends of *Plasmodium berghei* chromosomes. The telomeric probe was labeled using the DIG Oligonucleotide Tailing Kit, 2nd Generation (Cat. No. 03353383910, Roche, Mannheim, Germany).

After PFGE and before transfer, DNA from agarose gels were depurinated (20 min in 0.25 M HCL), denatured (2 × 20 min in 0.5N NaOH; 1.5 M NaCl) and neutralized (2 × 20 min in 0.5 M Tris HCl, pH 7.5; 1.5 M NaCl). Southern blotting was done on nylon membrane, positively charged (Cat. No. 1417240, Roche) using 10X SSC and followed by UV crosslinking of transferred DNA.

A membrane was hybridized overnight at 26LC with the telomeric probe and washed twice in 2X SSC and 0.1% SDS for 5 min. Then, the membrane was washed twice in 0.5X SSC and 0.1% SDS at 26°C for 20 min.

Bound probe was detected with disodium-2-chloro-5(4 methoxyspiro (1,2-dioxetane-3.2’-[5-chloro]tricycle[3.3.1.1.3.7 55] decan)-4-yl)-1-phenyl phosphate (CDP-StarTM, Cat. No.12041677001, Roche) according to the manufacturer’s instructions. All membranes were visualized using an Amersham ImageQuant 800 58 system (GE Healthcare Bio-Science AB, Uppsala, Sweden.

## Notes

### Competing Interest Statement

The authors have declared no competing interest.

## References

1. Amos B, Aurrecoechea C, Barba M, Barreto A, Basenko EY, Bazant W, et al. VEuPathDB: the eukaryotic pathogen, vector and host bioinformatics resource center. Nucleic Acids Res. 2022;50(D1):D898–D911. doi: 10.1093/nar/gkab929. PubMed PMID: 34718728; PubMed Central PMCID: PMCPMC8728164.

2. Rosenberg R, Lindsey NP, Fischer M, Gregory CJ, Hinckley AF, Mead PS, et al. Vital Signs: Trends in Reported Vectorborne Disease Cases - United States and Territories, 2004-2016. MMWR Morb Mortal Wkly Rep. 2018;67(17):496–501. Epub 20180504. doi: 10.15585/mmwr.mm6717e1. PubMed PMID: 29723166; PubMed Central PMCID: PMCPMC5933869.

3. Wikel SK. Ticks and Tick-Borne Infections: Complex Ecology, Agents, and Host Interactions. Vet Sci. 2018;5(2). Epub 20180620. doi: 10.3390/vetsci5020060. PubMed PMID: 29925800; PubMed Central PMCID: PMCPMC6024845.

4. Cornillot E, Hadj-Kaddour K, Dassouli A, Noel B, Ranwez V, Vacherie B, et al. Sequencing of the smallest Apicomplexan genome from the human pathogen Babesia microti. Nucleic Acids Res. 2012;40(18):9102–14. Epub 20120724. doi: 10.1093/nar/gks700. PubMed PMID: 22833609; PubMed Central PMCID: PMCPMC3467087.

5. Hildebrandt A, Zintl A, Montero E, Hunfeld KP, Gray J. Human Babesiosis in Europe. Pathogens. 2021;10(9). Epub 20210909. doi: 10.3390/pathogens10091165. PubMed PMID: 34578196; PubMed Central PMCID: PMCPMC8468516.

6. Asensi V, Gonzalez LM, Fernandez-Suarez J, Sevilla E, Navascues RA, Suarez ML, et al. A fatal case of Babesia divergens infection in Northwestern Spain. Ticks Tick Borne Dis. 2018;9(3):730–4. Epub 20180221. doi: 10.1016/j.ttbdis.2018.02.018. PubMed PMID: 29496491.

7. Gonzalez LM, Rojo S, Gonzalez-Camacho F, Luque D, Lobo CA, Montero E. Severe babesiosis in immunocompetent man, Spain, 2011. Emerg Infect Dis. 2014;20(4):724–6. doi: 10.3201/eid2004.131409. PubMed PMID: 24656155; PubMed Central PMCID: PMCPMC3966382.

8. Kumar A, O’Bryan J, Krause PJ. The Global Emergence of Human Babesiosis. Pathogens. 2021;10(11). Epub 20211106. doi: 10.3390/pathogens10111447. PubMed PMID: 34832603; PubMed Central PMCID: PMCPMC8623124.

9. Martinot M, Zadeh MM, Hansmann Y, Grawey I, Christmann D, Aguillon S, et al. Babesiosis in immunocompetent patients, Europe. Emerg Infect Dis. 2011;17(1):114–6. doi: 10.3201/eid1701.100737. PubMed PMID: 21192869; PubMed Central PMCID: PMCPMC3204631.

10. Schlogl KS, Hiesel JA, Wolf R, Kopacka I, Wagner P, Kastelic J, et al. Spatiotemporal cluster and incidence analysis of cattle mortality caused by bovine babesiosis in Styria, Austria, between 1998 and 2016. Parasitol Res. 2020;119(3):1117–23. Epub 20200225. doi: 10.1007/s00436-020-06604-8. PubMed PMID: 32100102; PubMed Central PMCID: PMCPMC7075847.

11. Herwaldt BL, Caccio S, Gherlinzoni F, Aspock H, Slemenda SB, Piccaluga P, et al. Molecular characterization of a non-Babesia divergens organism causing zoonotic babesiosis in Europe. Emerg Infect Dis. 2003;9(8):942–8. doi: 10.3201/eid0908.020748. PubMed PMID: 12967491; PubMed Central PMCID: PMCPMC3020600.

12. Herwaldt B, Persing DH, Precigout EA, Goff WL, Mathiesen DA, Taylor PW, et al. A fatal case of babesiosis in Missouri: identification of another piroplasm that infects humans. Ann Intern Med. 1996;124(7):643–50. doi: 10.7326/0003-4819-124-7-199604010-00004. PubMed PMID: 8607592.

13. Beattie JF, Michelson ML, Holman PJ. Acute babesiosis caused by Babesia divergens in a resident of Kentucky. N Engl J Med. 2002;347(9):697–8. doi: 10.1056/NEJM200208293470921. PubMed PMID: 12200568.

14. Herwaldt BL, de Bruyn G, Pieniazek NJ, Homer M, Lofy KH, Slemenda SB, et al. Babesia divergens-like infection, Washington State. Emerg Infect Dis. 2004;10(4):622–9. doi: 10.3201/eid1004.030377. PubMed PMID: 15200851; PubMed Central PMCID: PMCPMC3323086.

15. Holman PJ, Spencer AM, Droleskey RE, Goethert HK, Telford SR, 3rd. In vitro cultivation of a zoonotic Babesia sp. isolated from eastern cottontail rabbits (Sylvilagus floridanus) on Nantucket Island, Massachusetts. J Clin Microbiol. 2005;43(8):3995–4001. doi: 10.1128/JCM.43.8.3995-4001.2005. PubMed PMID: 16081941; PubMed Central PMCID: PMCPMC1233898.

16. Holman PJ, Spencer AM, Telford SR, 3rd, Goethert HK, Allen AJ, Knowles DP, et al. Comparative infectivity of Babesia divergens and a zoonotic Babesia divergens-like parasite in cattle. Am J Trop Med Hyg. 2005;73(5):865–70. PubMed PMID: 16282295.

17. Cuesta I, Gonzalez LM, Estrada K, Grande R, Zaballos A, Lobo CA, et al. High-Quality Draft Genome Sequence of Babesia divergens, the Etiological Agent of Cattle and Human Babesiosis. Genome Announc. 2014;2(6). Epub 20141113. doi: 10.1128/genomeA.01194-14. PubMed PMID: 25395649; PubMed Central PMCID: PMCPMC4241675.

18. Jackson AP, Otto TD, Darby A, Ramaprasad A, Xia D, Echaide IE, et al. The evolutionary dynamics of variant antigen genes in Babesia reveal a history of genomic innovation underlying host-parasite interaction. Nucleic Acids Res. 2014;42(11):7113–31. Epub 20140505. doi: 10.1093/nar/gku322. PubMed PMID: 24799432; PubMed Central PMCID: PMCPMC4066756.

19. Gonzalez LM, Estrada K, Grande R, Jimenez-Jacinto V, Vega-Alvarado L, Sevilla E, et al. Comparative and functional genomics of the protozoan parasite Babesia divergens highlighting the invasion and egress processes. PLoS Negl Trop Dis. 2019;13(8):e0007680. Epub 20190819. doi: 10.1371/journal.pntd.0007680. PubMed PMID: 31425518; PubMed Central PMCID: PMCPMC6715253.

20. Rezvani Y, Keroack CD, Elsworth B, Arriojas A, Gubbels MJ, Duraisingh MT, et al. Comparative single-cell transcriptional atlases of Babesia species reveal conserved and species-specific expression profiles. PLoS Biol. 2022;20(9):e3001816. Epub 20220922. doi: 10.1371/journal.pbio.3001816. PubMed PMID: 36137068; PubMed Central PMCID: PMCPMC9531838.

21. Ay F, Bunnik EM, Varoquaux N, Bol SM, Prudhomme J, Vert JP, et al. Three-dimensional modeling of the P. falciparum genome during the erythrocytic cycle reveals a strong connection between genome architecture and gene expression. Genome Res. 2014;24(6):974–88. Epub 20140326. doi: 10.1101/gr.169417.113. PubMed PMID: 24671853; PubMed Central PMCID: PMCPMC4032861.

22. Bunnik EM, Cook KB, Varoquaux N, Batugedara G, Prudhomme J, Cort A, et al. Changes in genome organization of parasite-specific gene families during the Plasmodium transmission stages. Nat Commun. 2018;9(1):1910. Epub 20180515. doi: 10.1038/s41467-018-04295-5. PubMed PMID: 29765020; PubMed Central PMCID: PMCPMC5954139.

23. Bunnik EM, Venkat A, Shao J, McGovern KE, Batugedara G, Worth D, et al. Comparative 3D genome organization in apicomplexan parasites. Proc Natl Acad Sci U S A. 2019;116(8):3183–92. Epub 20190205. doi: 10.1073/pnas.1810815116. PubMed PMID: 30723152; PubMed Central PMCID: PMCPMC6386730.

24. Singh P, Lonardi S, Liang Q, Vydyam P, Khabirova E, Fang T, et al. Babesia duncani multi-omics identifies virulence factors and drug targets. Nat Microbiol. 2023;8(5):845–59. Epub 20230413. doi: 10.1038/s41564-023-01360-8. PubMed PMID: 37055610; PubMed Central PMCID: PMCPMC10159843.

25. Muller LSM, Cosentino RO, Forstner KU, Guizetti J, Wedel C, Kaplan N, et al. Genome organization and DNA accessibility control antigenic variation in trypanosomes. Nature. 2018;563(7729):121–5. Epub 20181017. doi: 10.1038/s41586-018-0619-8. PubMed PMID: 30333624; PubMed Central PMCID: PMCPMC6784898.

26. Zintl A, Mulcahy G, Skerrett HE, Taylor SM, Gray JS. Babesia divergens, a bovine blood parasite of veterinary and zoonotic importance. Clin Microbiol Rev. 2003;16(4):622–36. doi: 10.1128/CMR.16.4.622-636.2003. PubMed PMID: 14557289; PubMed Central PMCID: PMCPMC207107.

27. Smit A, Hubley R, Green P. RepeatMasker Open-4.0. 2013–2015. 2015.

28. Goel M, Sun H, Jiao WB, Schneeberger K. SyRI: finding genomic rearrangements and local sequence differences from whole-genome assemblies. Genome Biol. 2019;20(1):277. Epub 20191216. doi: 10.1186/s13059-019-1911-0. PubMed PMID: 31842948; PubMed Central PMCID: PMCPMC6913012.

29. Goel M, Schneeberger K. plotsr: visualizing structural similarities and rearrangements between multiple genomes. Bioinformatics. 2022;38(10):2922–6. doi: 10.1093/bioinformatics/btac196. PubMed PMID: 35561173; PubMed Central PMCID: PMCPMC9113368.

30. Koren S, Walenz BP, Berlin K, Miller JR, Bergman NH, Phillippy AM. Canu: scalable and accurate long-read assembly via adaptive k-mer weighting and repeat separation. Genome Res. 2017;27(5):722–36. Epub 20170315. doi: 10.1101/gr.215087.116. PubMed PMID: 28298431; PubMed Central PMCID: PMCPMC5411767.

31. Manni M, Berkeley MR, Seppey M, Simao FA, Zdobnov EM. BUSCO Update: Novel and Streamlined Workflows along with Broader and Deeper Phylogenetic Coverage for Scoring of Eukaryotic, Prokaryotic, and Viral Genomes. Mol Biol Evol. 2021;38(10):4647–54. doi: 10.1093/molbev/msab199. PubMed PMID: 34320186; PubMed Central PMCID: PMCPMC8476166.

32. Baum BR, Ragan MA. The MRP method. Phylogenetic supertrees: combining information to reveal the Tree of Life. 2004:17–34.

33. Minh BQ, Hahn MW, Lanfear R. New Methods to Calculate Concordance Factors for Phylogenomic Datasets. Mol Biol Evol. 2020;37(9):2727–33. doi: 10.1093/molbev/msaa106. PubMed PMID: 32365179; PubMed Central PMCID: PMCPMC7475031.

34. Scornavacca C, Berry V, Lefort V, Douzery EJ, Ranwez V. PhySIC_IST: cleaning source trees to infer more informative supertrees. BMC Bioinformatics. 2008;9:413. Epub 20081004. doi: 10.1186/1471-2105-9-413. PubMed PMID: 18834542; PubMed Central PMCID: PMCPMC2576265.

35. Ranwez V, Criscuolo A, Douzery EJ. SuperTriplets: a triplet-based supertree approach to phylogenomics. Bioinformatics. 2010;26(12):i115–23. doi: 10.1093/bioinformatics/btq196. PubMed PMID: 20529895; PubMed Central PMCID: PMCPMC2881381.

36. Lopez-Rubio JJ, Mancio-Silva L, Scherf A. Genome-wide analysis of heterochromatin associates clonally variant gene regulation with perinuclear repressive centers in malaria parasites. Cell Host Microbe. 2009;5(2):179–90. doi: 10.1016/j.chom.2008.12.012. PubMed PMID: 19218088.

37. Jachowicz JW, Strehle M, Banerjee AK, Blanco MR, Thai J, Guttman M. Xist spatially amplifies SHARP/SPEN recruitment to balance chromosome-wide silencing and specificity to the X chromosome. Nat Struct Mol Biol. 2022;29(3):239–49. Epub 20220317. doi: 10.1038/s41594-022-00739-1. PubMed PMID: 35301492; PubMed Central PMCID: PMCPMC8969943.

38. Quinodoz S, Guttman M. Long noncoding RNAs: an emerging link between gene regulation and nuclear organization. Trends Cell Biol. 2014;24(11):651–63. Epub 20141023. doi: 10.1016/j.tcb.2014.08.009. PubMed PMID: 25441720; PubMed Central PMCID: PMCPMC4254690.

39. Rinn JL, Chang HY. Long Noncoding RNAs: Molecular Modalities to Organismal Functions. Annu Rev Biochem. 2020;89:283–308. doi: 10.1146/annurev-biochem-062917-012708. PubMed PMID: 32569523.

40. Amit-Avraham I, Pozner G, Eshar S, Fastman Y, Kolevzon N, Yavin E, et al. Antisense long noncoding RNAs regulate var gene activation in the malaria parasite Plasmodium falciparum. Proc Natl Acad Sci U S A. 2015;112(9):E982–91. Epub 20150217. doi: 10.1073/pnas.1420855112. PubMed PMID: 25691743; PubMed Central PMCID: PMCPMC4352787.

41. Epp C, Li F, Howitt CA, Chookajorn T, Deitsch KW. Chromatin associated sense and antisense noncoding RNAs are transcribed from the var gene family of virulence genes of the malaria parasite Plasmodium falciparum. RNA. 2009;15(1):116–27. Epub 20081126. doi: 10.1261/rna.1080109. PubMed PMID: 19037012; PubMed Central PMCID: PMCPMC2612763.

42. Andreoli TE, Schafer JA, Troutman SL. Perfusion rate-dependence of transepithelial osmosis in isolated proximal convoluted tubules: estimation of the hydraulic conductance. Kidney Int. 1978;14(3):263–9. doi: 10.1038/ki.1978.118. PubMed PMID: 723152.

43. Ramirez F, Bhardwaj V, Arrigoni L, Lam KC, Gruning BA, Villaveces J, et al. High-resolution TADs reveal DNA sequences underlying genome organization in flies. Nat Commun. 2018;9(1):189. Epub 20180115. doi: 10.1038/s41467-017-02525-w. PubMed PMID: 29335486; PubMed Central PMCID: PMCPMC5768762.

44. Varoquaux N, Ay F, Noble WS, Vert JP. A statistical approach for inferring the 3D structure of the genome. Bioinformatics. 2014;30(12):i26–33. doi: 10.1093/bioinformatics/btu268. PubMed PMID: 24931992; PubMed Central PMCID: PMCPMC4229903.

45. Deitsch KW, Dzikowski R. Variant Gene Expression and Antigenic Variation by Malaria Parasites. Annu Rev Microbiol. 2017;71:625–41. Epub 20170711. doi: 10.1146/annurev-micro-090816-093841. PubMed PMID: 28697665.

46. Jackson AP, Berry A, Aslett M, Allison HC, Burton P, Vavrova-Anderson J, et al. Antigenic diversity is generated by distinct evolutionary mechanisms in African trypanosome species. Proc Natl Acad Sci U S A. 2012;109(9):3416–21. Epub 20120213. doi: 10.1073/pnas.1117313109. PubMed PMID: 22331916; PubMed Central PMCID: PMCPMC3295286.

47. Dharia NV, Plouffe D, Bopp SE, Gonzalez-Paez GE, Lucas C, Salas C, et al. Genome scanning of Amazonian Plasmodium falciparum shows subtelomeric instability and clindamycin-resistant parasites. Genome Res. 2010;20(11):1534–44. Epub 20100909. doi: 10.1101/gr.105163.110. PubMed PMID: 20829224; PubMed Central PMCID: PMCPMC2963817.

48. Singh P, Pal AC, Mamoun CB. An Alternative Culture Medium for Continuous In Vitro Propagation of the Human Pathogen Babesia duncani in Human Erythrocytes. Pathogens. 2022;11(5). Epub 20220520. doi: 10.3390/pathogens11050599. PubMed PMID: 35631120; PubMed Central PMCID: PMCPMC9146245.

49. Edgar RC. Muscle5: High-accuracy alignment ensembles enable unbiased assessments of sequence homology and phylogeny. Nat Commun. 2022;13(1):6968. Epub 20221115. doi: 10.1038/s41467-022-34630-w. PubMed PMID: 36379955; PubMed Central PMCID: PMCPMC9664440.

50. Di Franco A, Poujol R, Baurain D, Philippe H. Evaluating the usefulness of alignment filtering methods to reduce the impact of errors on evolutionary inferences. BMC Evol Biol. 2019;19(1):21. Epub 20190111. doi: 10.1186/s12862-019-1350-2. PubMed PMID: 30634908; PubMed Central PMCID: PMCPMC6330419.

51. Ranwez V, Douzery EJP, Cambon C, Chantret N, Delsuc F. MACSE v2: Toolkit for the Alignment of Coding Sequences Accounting for Frameshifts and Stop Codons. Mol Biol Evol. 2018;35(10):2582–4. doi: 10.1093/molbev/msy159. PubMed PMID: 30165589; PubMed Central PMCID: PMCPMC6188553.

52. Nguyen LT, Schmidt HA, von Haeseler A, Minh BQ. IQ-TREE: a fast and effective stochastic algorithm for estimating maximum-likelihood phylogenies. Mol Biol Evol. 2015;32(1):268–74. Epub 20141103. doi: 10.1093/molbev/msu300. PubMed PMID: 25371430; PubMed Central PMCID: PMCPMC4271533.

53. Chernomor O, von Haeseler A, Minh BQ. Terrace Aware Data Structure for Phylogenomic Inference from Supermatrices. Syst Biol. 2016;65(6):997–1008. Epub 20160426. doi: 10.1093/sysbio/syw037. PubMed PMID: 27121966; PubMed Central PMCID: PMCPMC5066062.

54. Hoang DT, Chernomor O, von Haeseler A, Minh BQ, Vinh LS. UFBoot2: Improving the Ultrafast Bootstrap Approximation. Mol Biol Evol. 2018;35(2):518–22. doi: 10.1093/molbev/msx281. PubMed PMID: 29077904; PubMed Central PMCID: PMCPMC5850222.

55. Kalyaanamoorthy S, Minh BQ, Wong TKF, von Haeseler A, Jermiin LS. ModelFinder: fast model selection for accurate phylogenetic estimates. Nat Methods. 2017;14(6):587–9. Epub 20170508. doi: 10.1038/nmeth.4285. PubMed PMID: 28481363; PubMed Central PMCID: PMCPMC5453245.

56. Gadagkar SR, Rosenberg MS, Kumar S. Inferring species phylogenies from multiple genes: concatenated sequence tree versus consensus gene tree. J Exp Zool B Mol Dev Evol. 2005;304(1):64–74. doi: 10.1002/jez.b.21026. PubMed PMID: 15593277.

57. Seo TK, Kishino H, Thorne JL. Incorporating gene-specific variation when inferring and evaluating optimal evolutionary tree topologies from multilocus sequence data. Proc Natl Acad Sci U S A. 2005;102(12):4436–41. Epub 20050311. doi: 10.1073/pnas.0408313102. PubMed PMID: 15764703; PubMed Central PMCID: PMCPMC555482.

