## Supplementary tables for "Multiomics analysis reveals *B.* MO1 as a distinct *Babesia* species and provides insights into its evolution and virulence"

**Table S1:** Molecule size for the optical map for *B.* MO1

| Optical Molecule # | Length (bp) |
|---|---|
| 1 | 4,416,462 |
| 2 | 3,532,684 |
| 3 | 2,290,926 |
| 4 | 507,370 |
| 5 | 272,473 |
| 6 | 230,786 |
| 15 | 137,680 |
| 7 | 83,877 |
| SUM | 11,472,258 |

**Table S2A:** Genome-wide read count Pearson correlations- ChIP-Seq analysis on *B.* MO1 clone F12

| | | | | | |
|---|---|---|---|---|---|
| H3K9me3_rep1 | 1 | | | | |
| H3K9me3_rep2 | 0.970557561 | 1 | | | |
| H3K9ac_rep1 | -0.226618949 | -0.221437881 | 1 | | |
| H3K9ac_rep2 | -0.225918015 | -0.22061916 | 0.993004902 | 1 | |
| IgG | 0.407501319 | 0.407465025 | 0.459774564 | 0.458195753 | 1 |
| | H3K9me3_rep1 | H3K9me3_rep2 | H3K9ac_rep1 | H3K9ac_rep2 | IgG |

**Table S2B:** Genome-wide read count Pearson correlations- ChIP-Seq analysis on *B.* MO1 clone B12

| | | | | | |
|---|---|---|---|---|---|
| H3K9me3_rep1 | 1 | | | | |
| H3K9me3_rep2 | 0.958261066 | 1 | | | |
| H3K9ac_rep1 | -0.227423493 | -0.228878465 | 1 | | |
| H3K9ac_rep2 | -0.227801951 | -0.229025069 | 0.992019183 | 1 | |
| IgG | 0.333857894 | 0.30096601 | 0.133843448 | 0.126827877 | 1 |
| | H3K9me3_rep1 | H3K9me3_rep2 | H3K9ac_rep1 | H3K9ac_rep2 | IgG |
