## Supplementary figures for "Multiomics analysis reveals *B.* MO1 as a distinct *Babesia* species and provides insights into its evolution and virulence"

### Figure S1

**A**

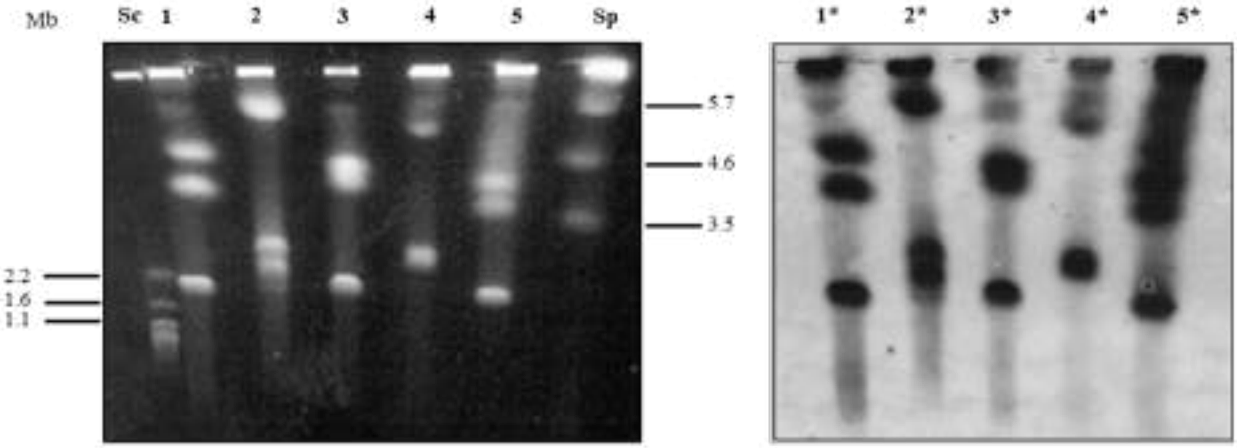


**B**

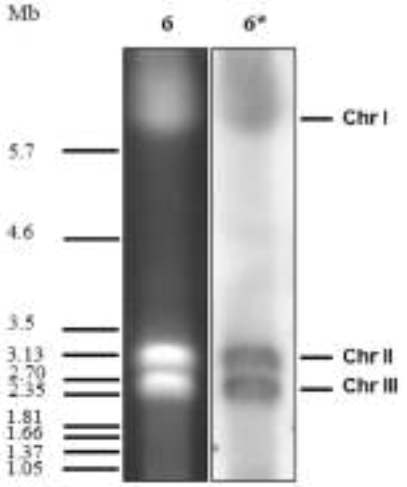


| SAMPLE | 1 | 2 | 3 | 4 | 5 | 6 |
|---|---|---|---|---|---|---|
| HOST | Bovine | Bovine | Human | Bovine | Bovine | Human |
| ISOLATE ORIGIN | 5008B Manche (Fr) | 6303E Puy de Dome (Fr) | Rouen87 Seine-Maritime (Fr) | Y5 Ireland | W8843 Scotland | Asturias Spain |
| Chr I (Mb) | 4.8 Mb | 5.7 Mb | 4.7 Mb | 5.0 Mb | 4.6 Mb | 5.7 Mb |
| Chr II (Mb) | 3.5 Mb | 2.8 Mb | 4.1 Mb | 2.9 Mb | 3.7 Mb | 3.13 |
| Chr III (Mb) | 2.2 Mb | 2.3 Mb | 2.2 Mb | 2.9 Mb | 2.2 Mb | 2.7 |
| GENOME SIZE | 10.5 Mb | 10.8 Mb | 11.0 Mb | 10.8 Mb | 10.5 Mb | 11.55 |

### Figure S2 A

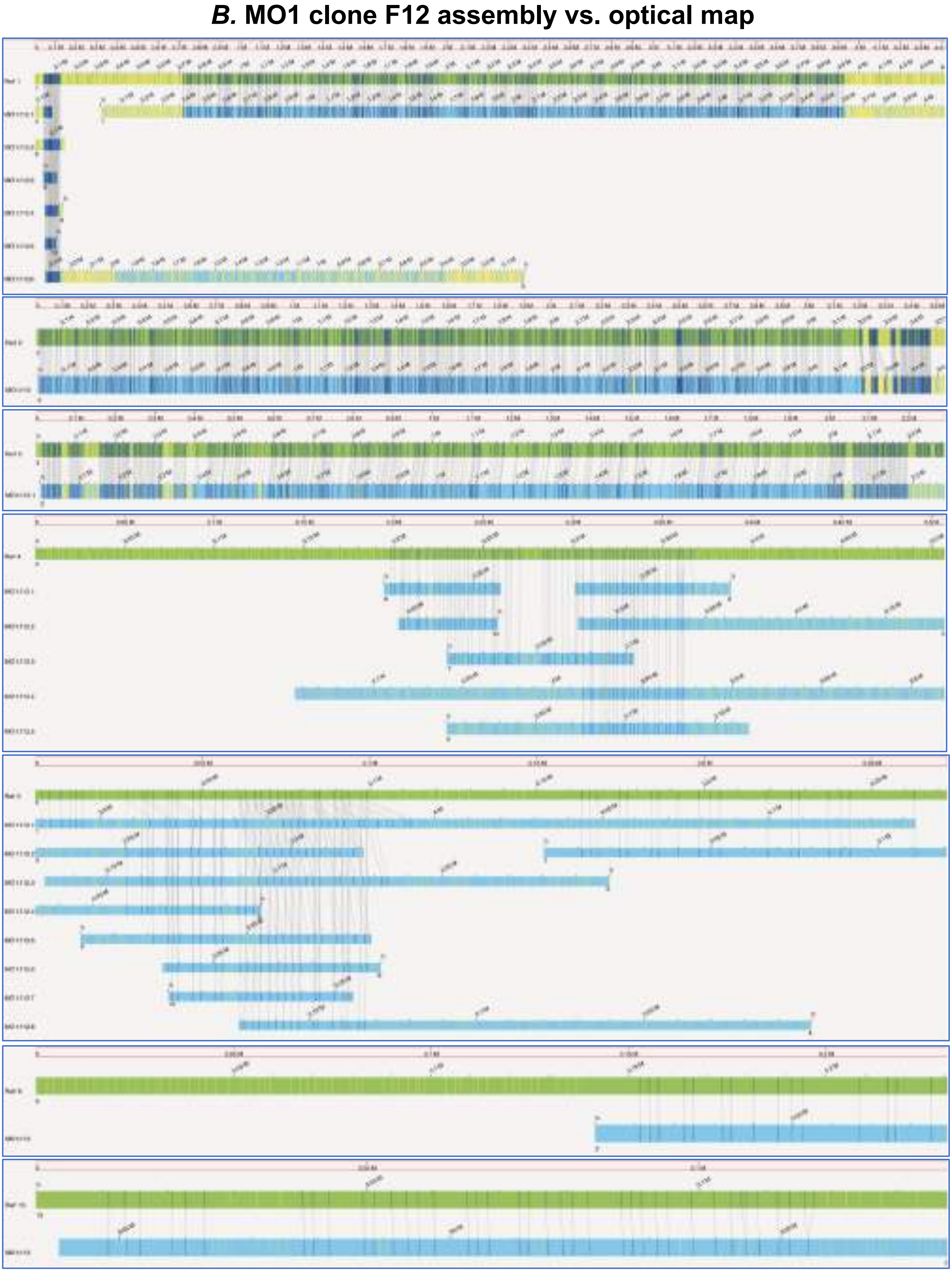

### Figure S2 B

***B.* MO1 clone B12 assembly vs. optical map**

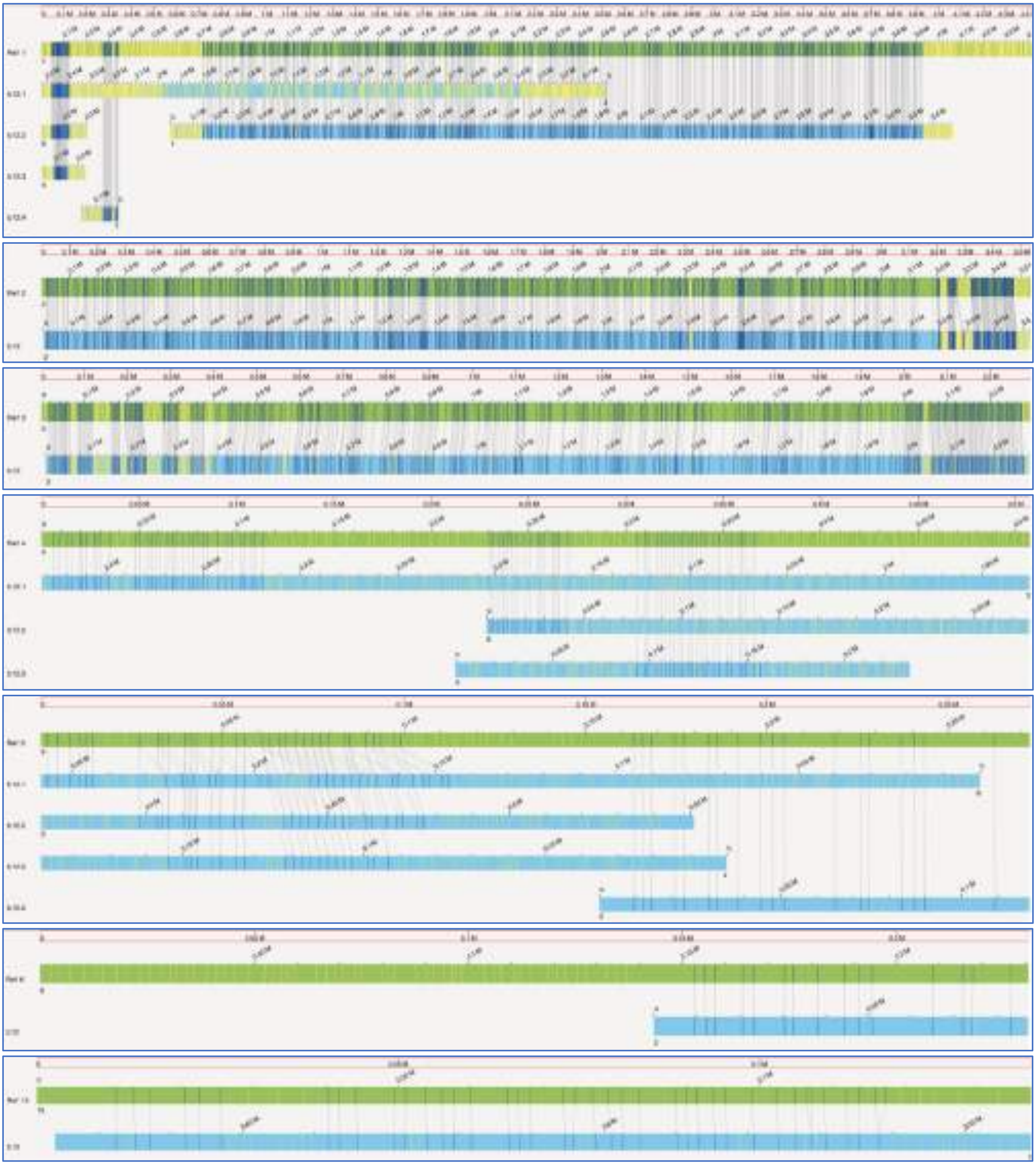

### Figure S3

#### Synteny plot (clone F12, clone B12, parental *B.* MO1)

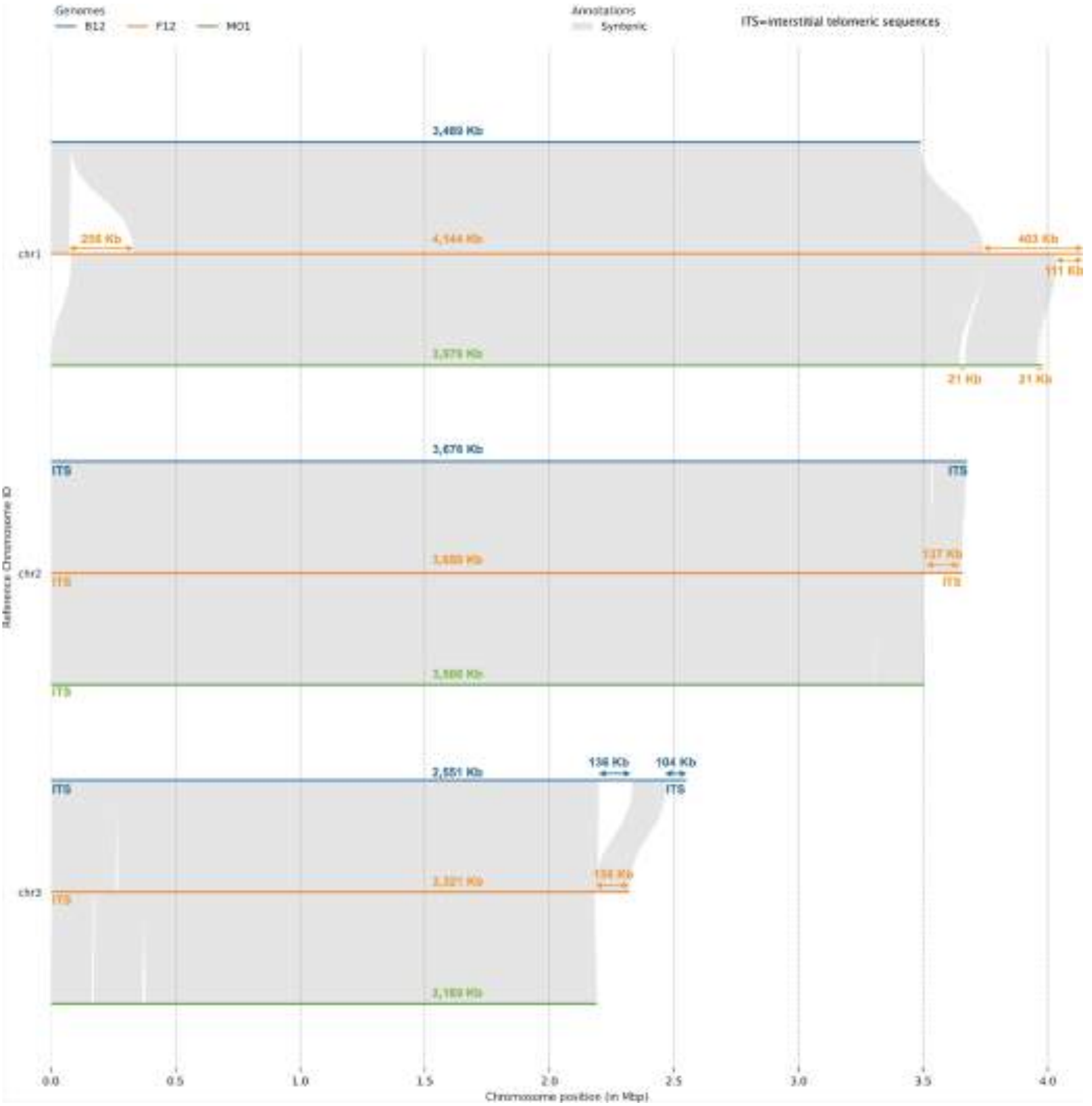

### Figure S4A

#### *B.* MO1 clone F12 vs clone B12

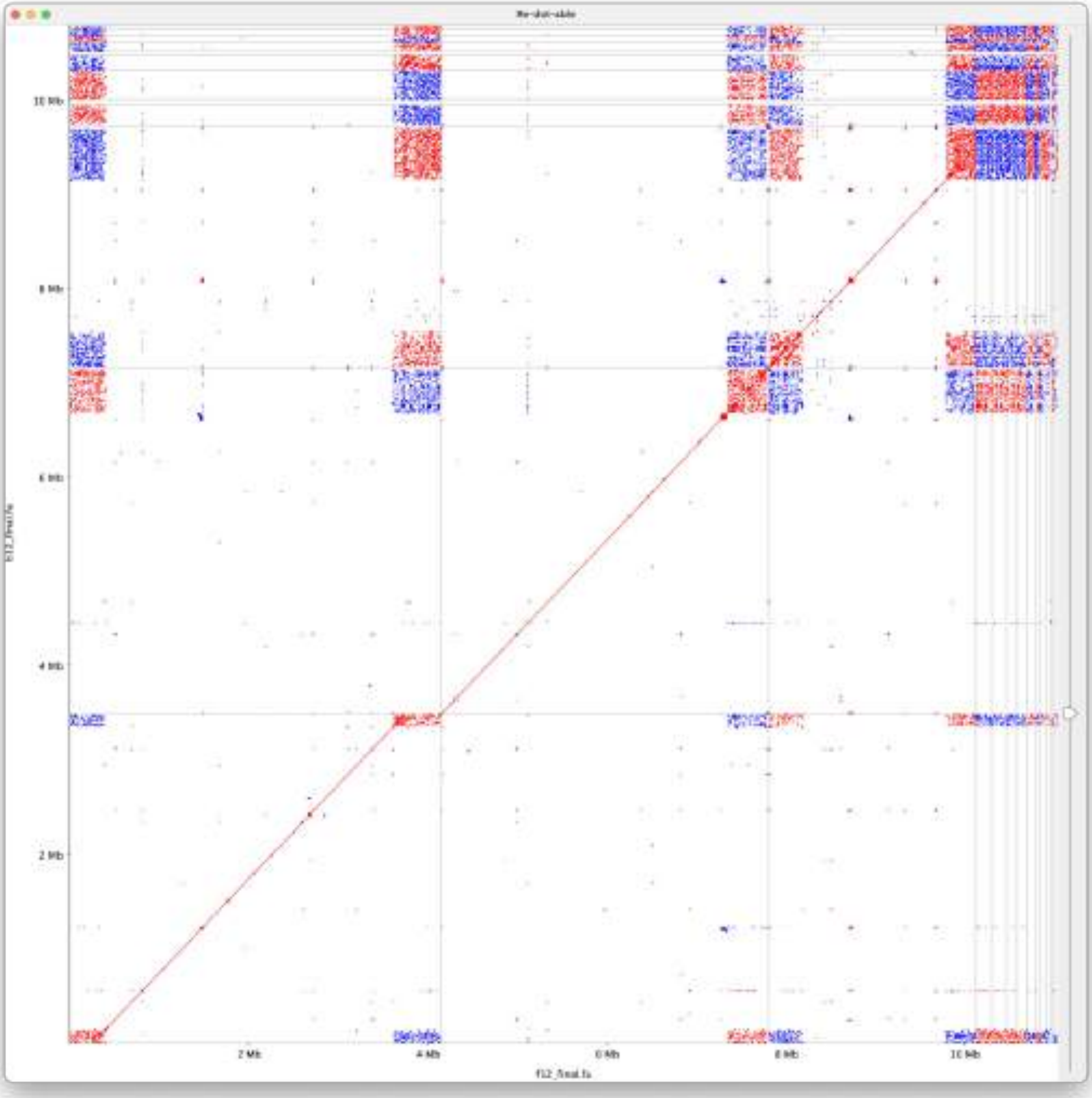

**Figure S4B**

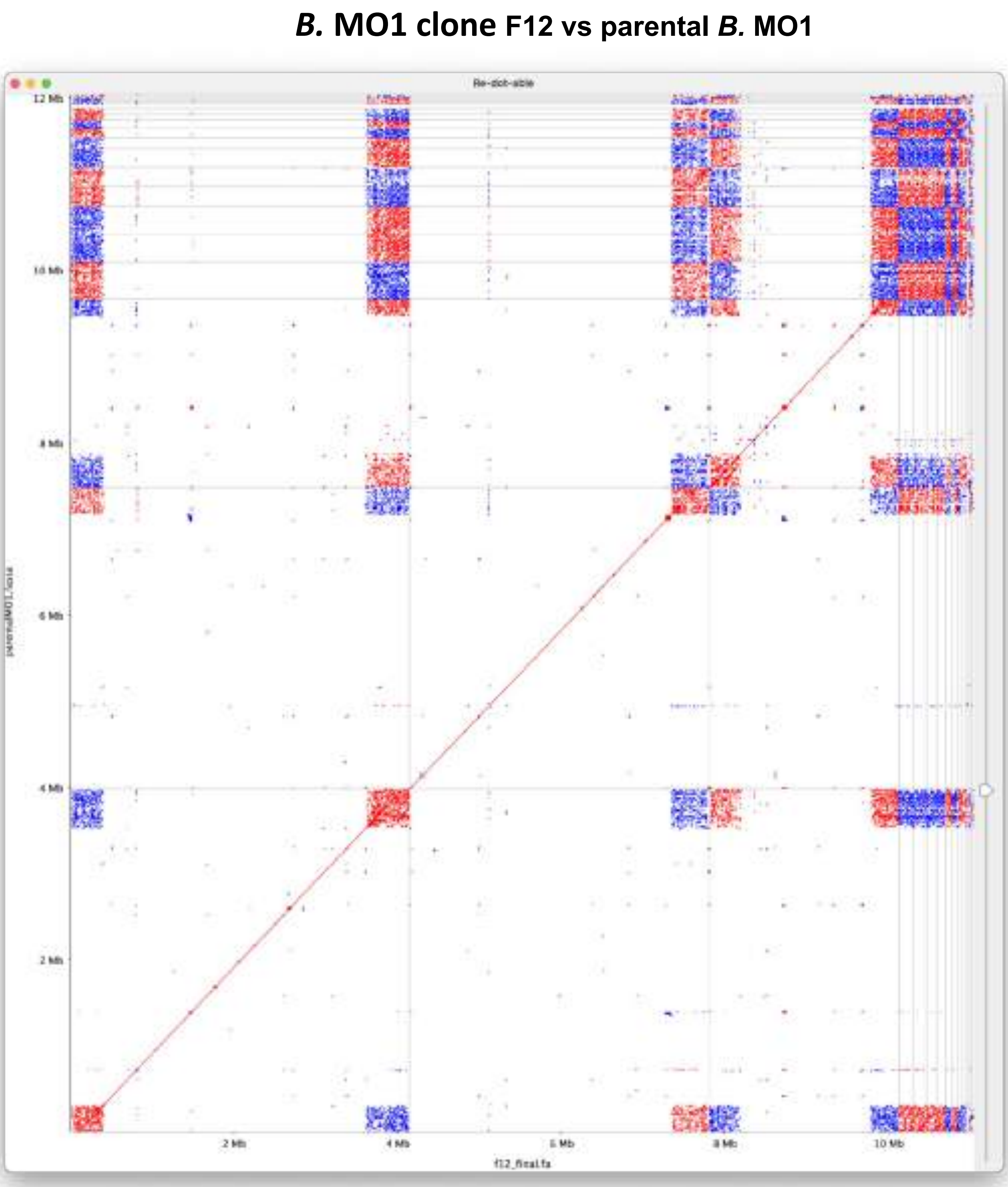
*B.* MO1 clone F12 vs parental *B.* MO1
Re-dot-able
12 Mb
10 Mb
8 Mb
6 Mb
4 Mb
2 Mb
2 Mb
4 Mb
6 Mb
8 Mb
10 Mb
f12_final.fa

### Figure S5

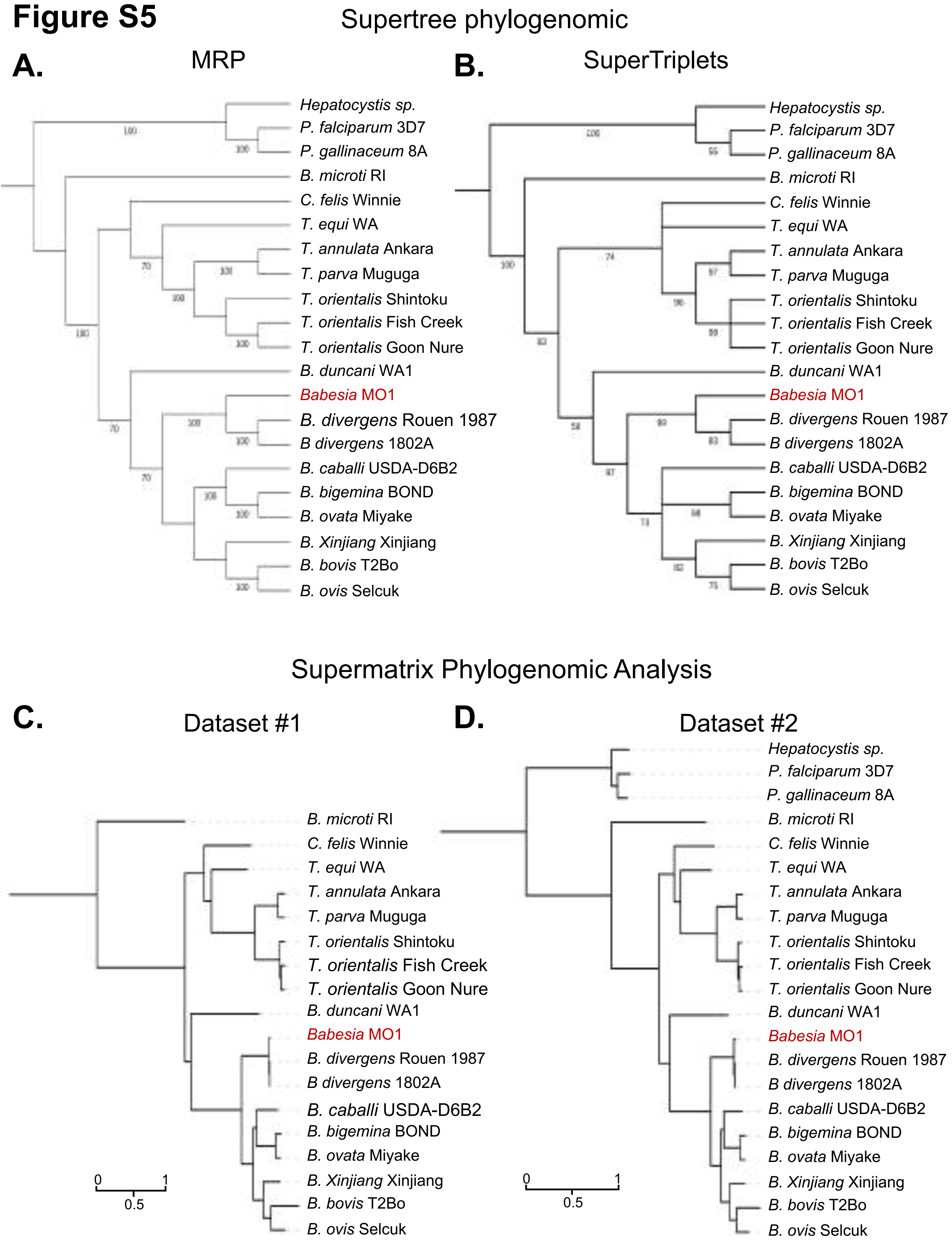
Supertree phylogenomic
A.
MRP
Hepatocystis sp.
P. falciparum 3D7
P. gallinaceum 8A
B. microti RI
C. felis Winnie
T. equi WA
T. annulata Ankara
T. parva Muguga
T. orientalis Shintoku
T. orientalis Fish Creek
T. orientalis Goon Nure
B. duncani WA1
Babesia MO1
B. divergens Rouen 1987
B divergens 1802A
B. caballi USDA-D6B2
B. bigemina BOND
B. ovata Miyake
B. Xinjiang Xinjiang
B. bovis T2Bo
B. ovis Selcuk
B.
SuperTriplets
Hepatocystis sp.
P. falciparum 3D7
P. gallinaceum 8A
B. microti RI
C. felis Winnie
T. equi WA
T. annulata Ankara
T. parva Muguga
T. orientalis Shintoku
T. orientalis Fish Creek
T. orientalis Goon Nure
B. duncani WA1
Babesia MO1
B. divergens Rouen 1987
B divergens 1802A
B. caballi USDA-D6B2
B. bigemina BOND
B. ovata Miyake
B. Xinjiang Xinjiang
B. bovis T2Bo
B. ovis Selcuk
Supermatrix Phylogenomic Analysis
C.
Dataset #1
B. microti RI
C. felis Winnie
T. equi WA
T. annulata Ankara
T. parva Muguga
T. orientalis Shintoku
T. orientalis Fish Creek
T. orientalis Goon Nure
B. duncani WA1
Babesia MO1
B. divergens Rouen 1987
B divergens 1802A
B. caballi USDA-D6B2
B. bigemina BOND
B. ovata Miyake
B. Xinjiang Xinjiang
B. bovis T2Bo
B. ovis Selcuk
0
0.5
1
D.
Dataset #2
Hepatocystis sp.
P. falciparum 3D7
P. gallinaceum 8A
B. microti RI
C. felis Winnie
T. equi WA
T. annulata Ankara
T. parva Muguga
T. orientalis Shintoku
T. orientalis Fish Creek
T. orientalis Goon Nure
B. duncani WA1
Babesia MO1
B. divergens Rouen 1987
B divergens 1802A
B. caballi USDA-D6B2
B. bigemina BOND
B. ovata Miyake
B. Xinjiang Xinjiang
B. bovis T2Bo
B. ovis Selcuk
0
0.5
1

#### Figure S6

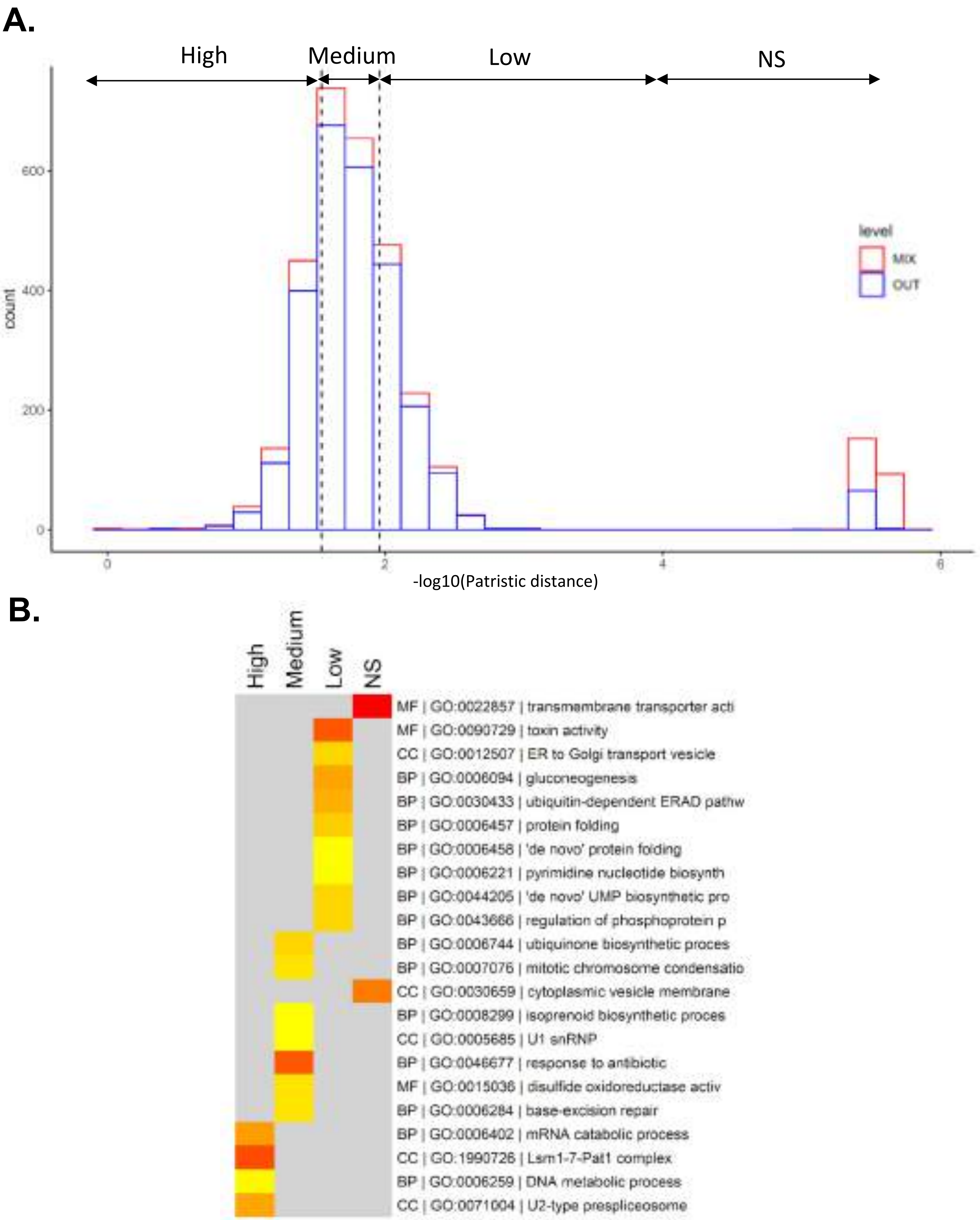
A.
High
Medium
Low
NS
level
MIX
OUT
count
600
400
200
0
0
2
4
6
-log10(Patristic distance)
B.
High
Medium
Low
NS
MF | GO:0022857 | transmembrane transporter acti
MF | GO:0090729 | toxin activity
CC | GO:0012507 | ER to Golgi transport vesicle
BP | GO:0006094 | gluconeogenesis
BP | GO:0030433 | ubiquitin-dependent ERAD pathw
BP | GO:0006457 | protein folding
BP | GO:0006458 | 'de novo' protein folding
BP | GO:0006221 | pyrimidine nucleotide biosynth
BP | GO:0044205 | 'de novo' UMP biosynthetic pro
BP | GO:0043666 | regulation of phosphoprotein p
BP | GO:0006744 | ubiquinone biosynthetic proces
BP | GO:0007076 | mitotic chromosome condensatio
CC | GO:0030659 | cytoplasmic vesicle membrane
BP | GO:0008299 | isoprenoid biosynthetic proces
CC | GO:0005685 | U1 snRNP
BP | GO:0046677 | response to antibiotic
MF | GO:0015036 | disulfide oxidoreductase activ
BP | GO:0006284 | base-excision repair
BP | GO:0006402 | mRNA catabolic process
CC | GO:1990726 | Lsm1-7-Pat1 complex
BP | GO:0006259 | DNA metabolic process
CC | GO:0071004 | U2-type prespliceosome

#### Figure S7

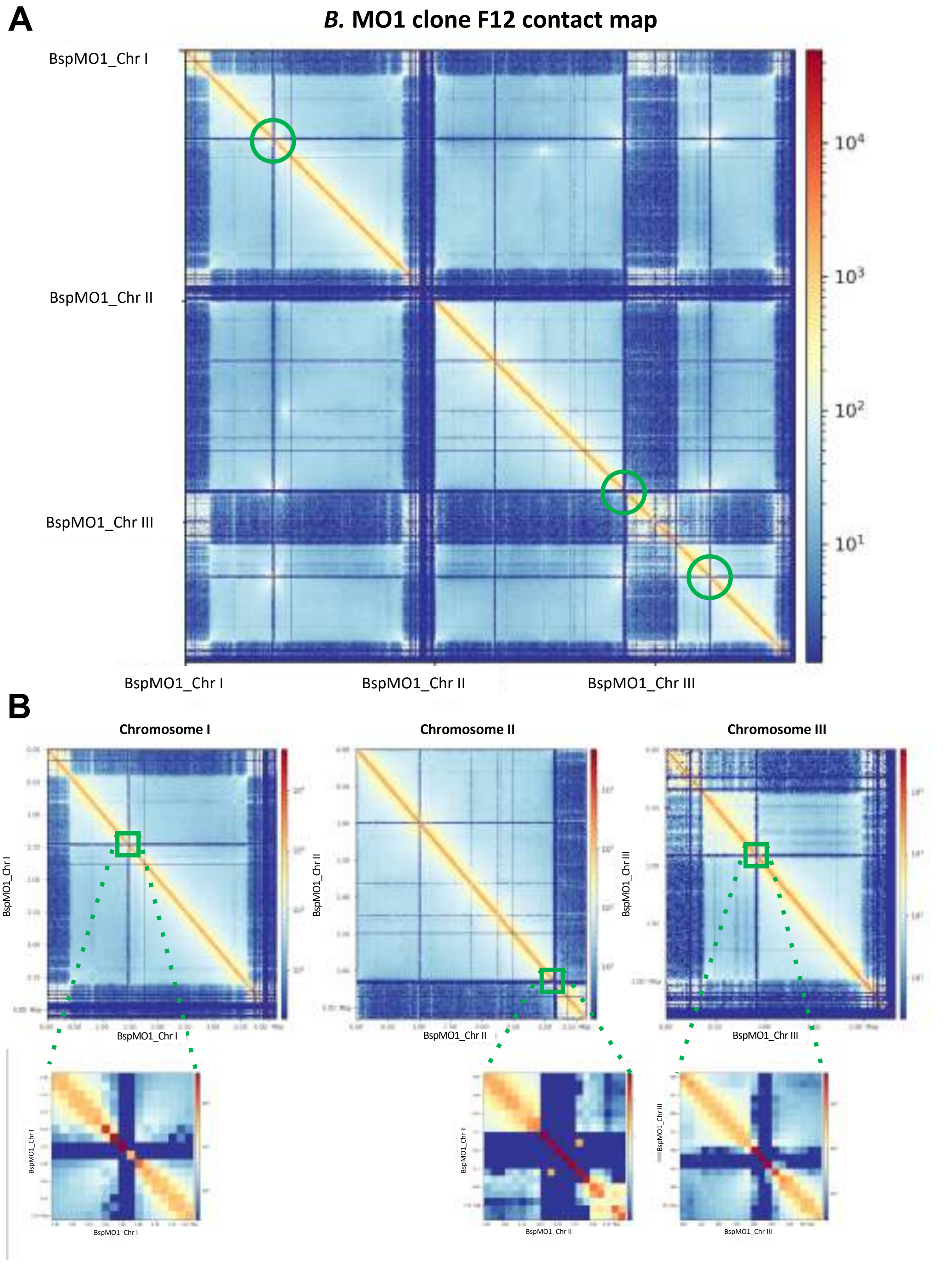
A
B. MO1 clone F12 contact map
BspMO1_Chr I
BspMO1_Chr II
BspMO1_Chr III
BspMO1_Chr I
BspMO1_Chr II
BspMO1_Chr III
$10^4$
$10^3$
$10^2$
$10^1$
B
Chromosome I
Chromosome II
Chromosome III
BspMO1_Chr I
BspMO1_Chr II
BspMO1_Chr III

### Figure S8

A

***B.* MO1 clone B12 contact map**

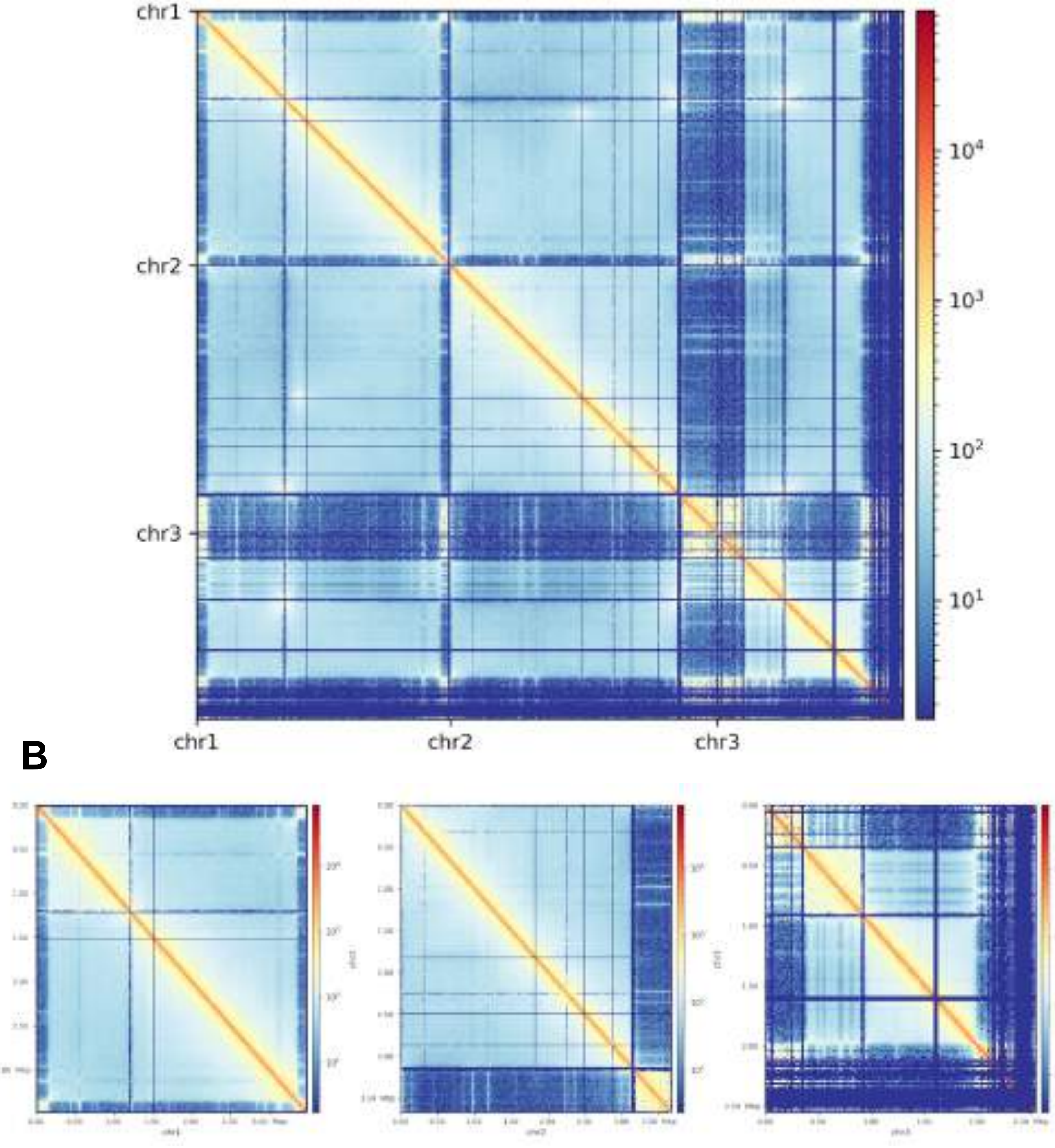


B

**Figure S9**

**A**

***B. divergens*** **ROUEN 87 contact map**

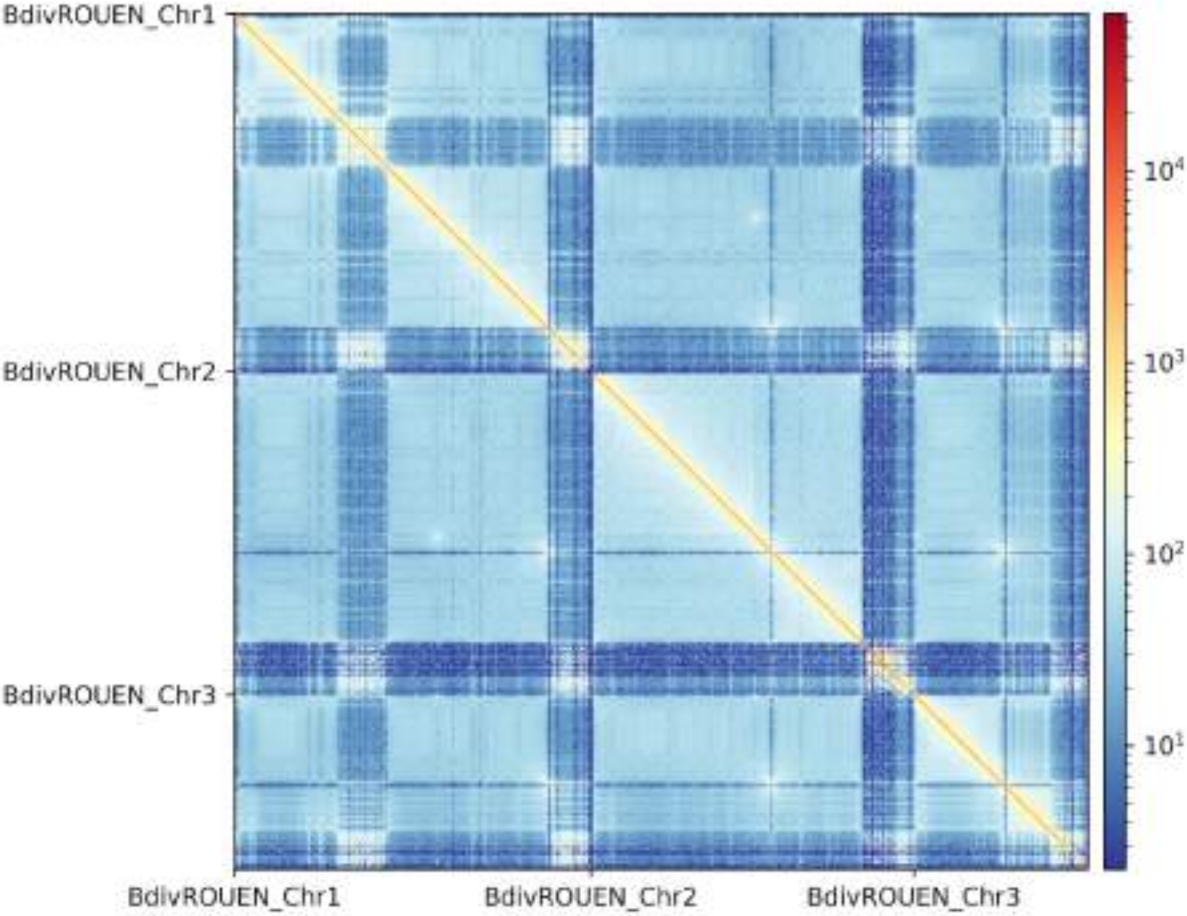


**B**

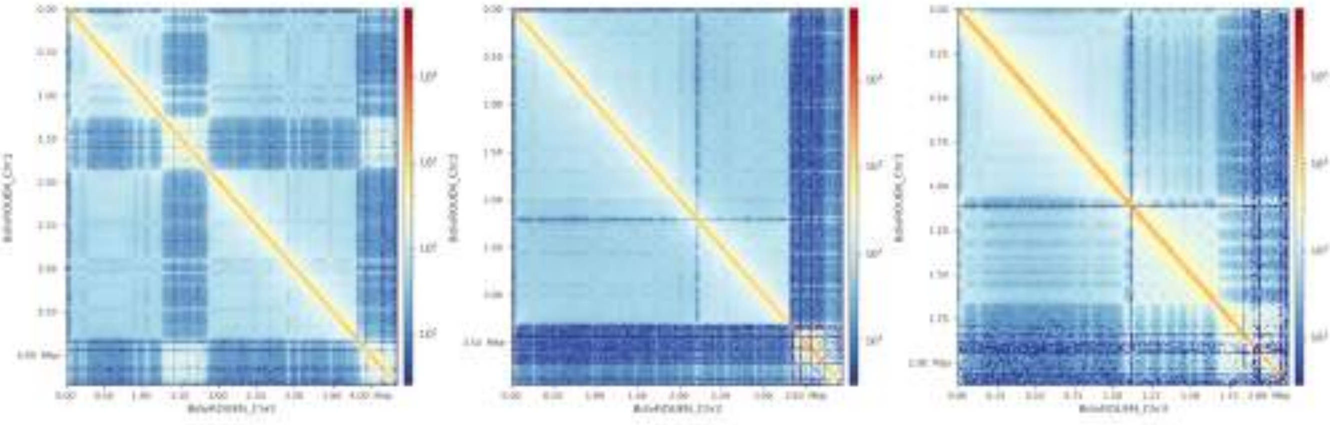

### Figure S10

#### GC skew plots

##### *B.* MO1 clone F12

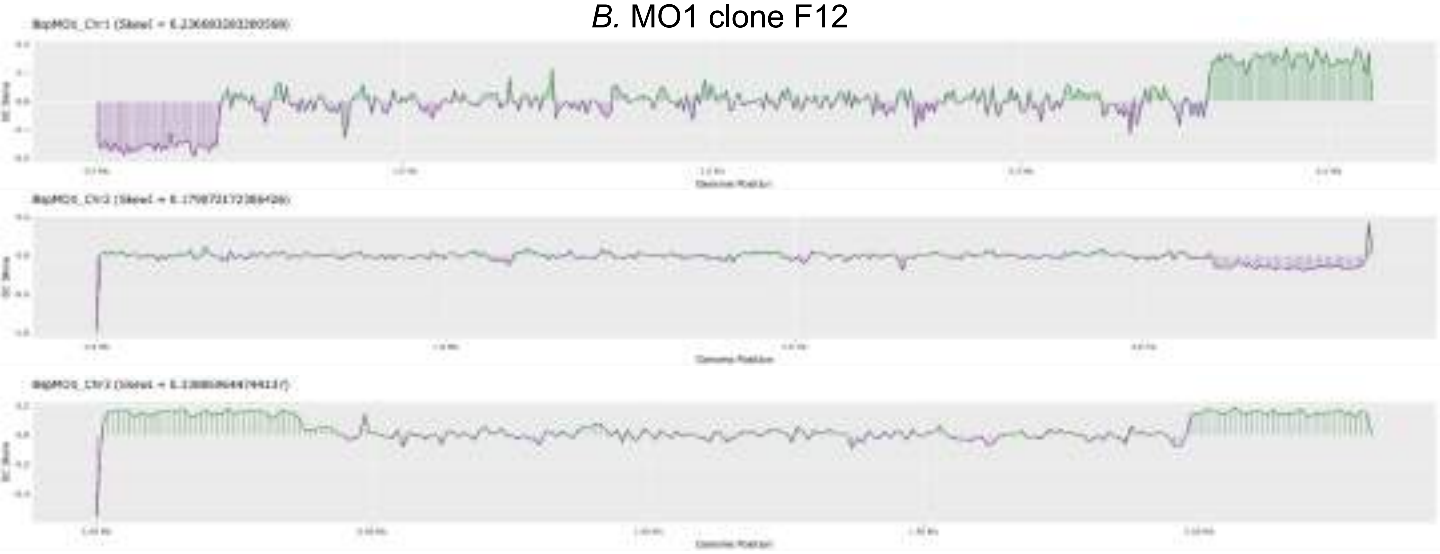

##### *B.* MO1 clone B12

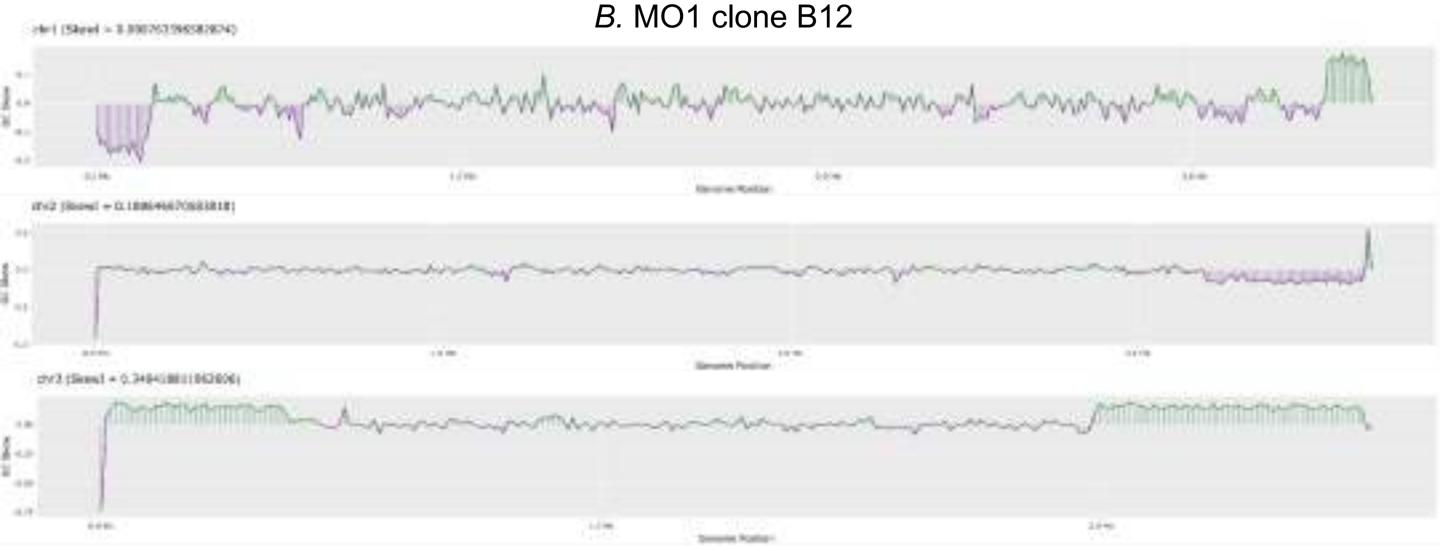

##### *B. divergens* ROUEN 87

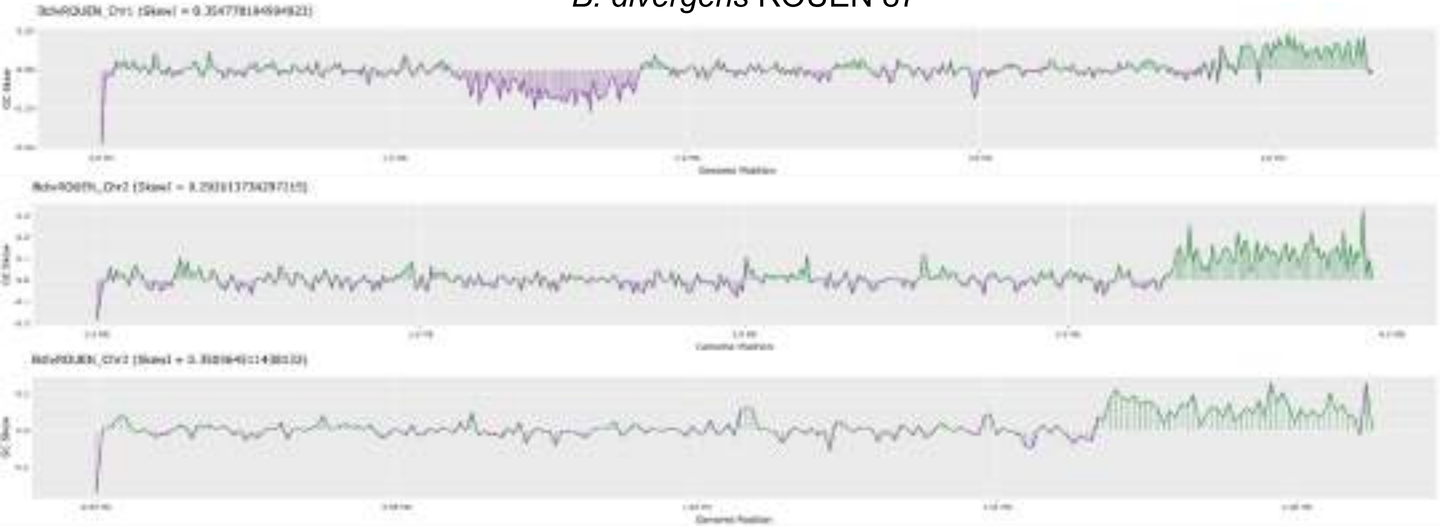

### Figure S11

### DHFR-TS

```
Babesia MO1            1   ---MVANYEGCGGLEIYVATAWN    RAIGFKNDIPWPHIREDFRFLARGTSYVDPEVKAK    NPDLMNVVVMGRKT  69
B.divergens Rouen 87   1   ---MVANYEGCGGLEIFVATAWN    RAIGFKNDIPWPHIREDFRFLARGTSYVDPEVKAK    NPDLMNVVIMGRKT  69
P.falciparum 3D7       1   mMeQVCDVFDIYAICACCKVESK[14]RGLGNKGVLPWKCNSLDMKYFCAVTTYVNESKYEK[22]SKKLQNVVVMGRTS  108
P.falciparum HB3       1   mMeQVCDVFDIYAICACCKVESK[14]RGLGNKGVLPWKCNSLDMKYFCAVTTYVNESKYEK[22]SKKLQNVVVMGRTN  108
B.duncani WA1          1   -MdLSTKYEGFSPIVMFVATDVK    GGIGFEGKIPWPHIPMDFSFFFRGTCYVEPEILNR    YPEIQNVVIFGRKT  71
B.microti              1   -M-------GMYKVCSIYASTPN    GGIGNEGKLPWKTLPRDLKHLQDITTAYGPD----    -HSVQNVVIMGRKT  59
Babesia bovis          1   ---MSNSYEGCGDLTIFVAVALN    KVIGHKNQIPWPHITHDFRFLRNGTTYIPPEVLSK    NPDIQNVVIFGRKT  69

Babesia MO1            70  YESIPASSRPLKNRINVVLSRNVK--EI-PGCLVFPSLTEAIRHVRSSVPHYKIFCLGGGEVYREVMENDLCDRIYLTRL  146
B.divergens Rouen 87   70  YESIPESSRPLKNRINVVLSRNVK--DI-PGCLVFPSLTEAIRHVRSSVPHHKIFCLGGGEVYREVMENDLCDRIYLTRL  146
P.falciparum 3D7       109 WESIPKKFKPLSNRINVILSRTLKkeDFdEDVYIINKVEDLIVLL-GKLNYYKCFIIGGSVVYQEFLEKKLIKKIYFTRI  187
P.falciparum HB3       109 WESIPKKFKPLSNRINVILSRTLKkeDFdEDVYIINKVEDLIVLL-GKLNYYKCFIIGGSVVYQEFLEKKLIKKIYFTRI  187
B.duncani WA1          72  YESIPANVFPLKKRHNVIISRTLT--HV-PGASVFNNLDDALRWVNESKRFHKTIIMGGVEIYKLALETGLVEKIYLTRI  148
B.microti              60  YISIPKSSRPLKDRINIVLSSSVS--DFgDGVIAAKSMQDAFDKL-EKMKFNKIFIIGGSSVYKEAYDLGIVEKVYVTRV  136
Babesia bovis          70  YESIPKASLPLKNRINVILSRTVK--EV-PGCLVYEDLSTAIRDLRANVPHNKIFILGGSFLYKEVLDNGLCDKIYLTRL  146

Babesia MO1            147 TEEYEGDVFFPEIPD-TFQITGISKTFSTDYVTFDFVVYEKVG    AIK-EKRPPTFDELLLT    GGELTVPTPKYV  216
B.divergens Rouen 87   147 TEEYEGDVFFPEIPD-TFQITGISKTFSTDYVTFDFVVYEKTG    AIK-EKRPPTFDELLLT    GGELTVPTPKYV  216
P.falciparum 3D7       188 NSTYECDVFFPEINEnEYQIISVSDVYTSNNTTLDFIIYKKTN[30]MKKLTEFYKNVDKYKIN[24]KNKNSIHPNDFQ  313
P.falciparum HB3       188 NSTYECDVFFPEINEnEYQIISVSDVYTSNNTTLDFIIYKKTN[30]MKKLTEFYKNVDKYKIN[24]KNKNSIHPNDFQ  313
B.duncani WA1          149 GNEFKADRFFPDITK-DFEIVGISQTFGTDFTTFDFVIYEKKG    LNQLDHVTKSFDEMLLT    GKPLKTVAPLYK  219
B.microti              137 NKELPADTFVTSVPP-IFEIVGISRTFSYNDIPFDFIIYMLKD    SRATSNVCVDIDDYLLT    EHEIRDFKYQFK  207
Babesia bovis          147 NKEYPGDTYFPDIPD-TFEITAISPTFSTDFVSYDFVIYERKD    CKT-VFPDPPFDQLLMT    GTDISVPKPKYV  216

Babesia MO1            217 ACPSVKVRYHQEFQYLDIIADVLSTGTLKPNRTGVDGISKFGYQMHFDLSQSFPLLTTKKVALRSIIEELLWFIRGSTNG  296
B.divergens Rouen 87   217 ACPSVKVRYHQEFQYLDIIADVLSTGTLKPNRTGVDGISKFGYQMHFDLSQSFPLLTTKKVALRSIIEELLWFIRGSTNG  296
P.falciparum 3D7       314 IYNSLKYKYHPEYQYLNIIYDIMMNGNKQSDRTGVGVLSKFGYIMKFDLSQYFPLLTTKKLFLRGIIEELLWFIRGETNG  393
P.falciparum HB3       314 IYNSLKYKYHPEYQYLNIIYDIMMNGNKQSDRTGVGVLSKFGYIMKFDLSQYFPLLTTKKLFLRGIIEELLWFIRGETNG  393
B.duncani WA1          220 ACPNIRIRYHQEFQYLDICADVLSTGTLKENRTGIDALSKFGYQMRFDLSQSFPLLTTKKVFLRGITEELIWFINGSTNG  299
B.microti              208 ALPNVTIRKHEEIQYLDIIADILSSGSENDDRTGVGVLSKFGYKMEFNLDESFPLLTTKKVFFKGIVEELLWFIKGDTSG  287
Babesia bovis          217 ACPGVRIRNHEEFQYLDILADVLSHGVLKPNRTGTDAYSKFGYQMRFDLSRSFPLLTTKKVALRSIIEELLWFIKGSTNG  296

Babesia MO1            297 NELLDKNVRIWELNGARSFLDNLGFTDREEHDLGPVYGFQWRHFGAKYLDMHADYSGQGIDQLKVVINKIKTNPNDRRLI  376
B.divergens Rouen 87   297 NELLDKNVRIWELNGARSFLDNLGFTDREEHDLGPVYGFQWRHFGAKYLDMHADYSGQGIDQLKVVINKIKTNPNDRRLI  376
P.falciparum 3D7       394 NTLLNKNVRIWEANGTREFLDNRKLFHREVNDLGPIYGFQWRHFGAEYTNMYDNYENKGVDQLKNIINLIKNDPTSRRIL  473
P.falciparum HB3       394 NTLLNKNVRIWEANGTREFLDNRKLFHREVNDLGPIYGFQWRHFGAEYTNMYDNYENKGVDQLKNIINLIKNDPTSRRIL  473
B.duncani WA1          300 NDLLKKNVRIWELNGKRSFLDNLGFYNREEHDLGPVYGFQWRHFGATYTDMHADYKGQGIDQLVNVINSIKNDPNSRRLI  379
B.microti              288 KILLEKGVRIWEKNGTREFLDSVGLNERKEHDLGPIYGFQWRHFGAEYKDCDTNYTGQGIDQLMEAIDKIKNDPNSRRII  367
Babesia bovis          297 NDLLAKNVRIWELNGRRDFLDKNGFTDREEHDLGPIYGFQWRHFGAEYLDMHADYTGKGIDQLAEIINRIKTNPNDRRLI  376

Babesia MO1            377 ICSWNVADLSKMALPPCHCLYQFYVRDGKLSCMLHQRSCDLGLGVPFNIASYAILTAMIAQVCGLKLGEFVHNLADAHVY  456
B.divergens Rouen 87   377 ICSWNVADLSKMALPPCHCLYQFYVRDGKLSCMLHQRSCDLGLGVPFNIASYAILTAMIAQVCGLKLGEFVHNLADAHVY  456
P.falciparum 3D7       474 LCAWNVKDLDQMALPPCHILCQFYVFDGKLSCIMYQRSCDLGLGVPFNIASYSIFTHMIAQVCNLQPAQFIHVLGNAHVY  553
P.falciparum HB3       474 LCAWNVKDLDQMALPPCHILCQFYVFDGKLSCIMYQRSCDLGLGVPFNIASYSIFTHMIAQVCNLQPAQFIHVLGNAHVY  553
B.duncani WA1          380 VCSWNVSDVPKMALPPCHLLFQFYVAQGKLSCMLHQRSCDLGLGVPFNIASYSILTAMIAQVCNLQLGEFVHNLADAHIY  459
B.microti              368 VCSWNVNDLSKMALPPCHCLFQFYVSQGRLSCIMYQRSADVGLGVPFNIASYSLLTLMIAQVCALRPGKFVHVLGNAHIY  447
Babesia bovis          377 VCSWNVSDLKKMALPPCHCFFQFYVSDNKLSCMMHQRSCDLGLGVPFNIASYSILTAMVAQVCGLGLGEFVHNLADAHIY  456

Babesia MO1            457 IDHVDAMKLQMSRIPYPFPQLKLNPAITNIEDFTIDDIIVQNYVSHPPIKMAMSA  511
B.divergens Rouen 87   457 VDHVDAMKLQMSRIPYPFPQLKLNPAIKNIEDFTIDDIIVQNYVSHPPIKMAMSA  511
P.falciparum 3D7       554 NNHIDSLKIQLNRIPYPFPTLKLNPDIKNIEDFTISDFTIQNYVHHEKISMDMAA  608
P.falciparum HB3       554 NNHIDSLKIQLNRIPYPFPTLKLNPDIKNIEDFTISDFTIQNYVHHEKISMDMAA  608
B.duncani WA1          460 VNHIDALKEQLTRVPYPFPLLKLNKDITDICDFKLEDVKVEGYSCHPTIKMEMAA  514
B.microti              448 KTHITALTKQIQRIPYPFPILKLNAQVKRIEDFVPSDINLLCYTCHPTIKMDMAA  502
Babesia bovis          457 VDHVDAVTTQIARIPHPFPRLRLNPDIRNIEDFTIDDIVVEDYVSHPPIPMAMSA  511
```

#### Figure S12

*B. divergens* Rouen87

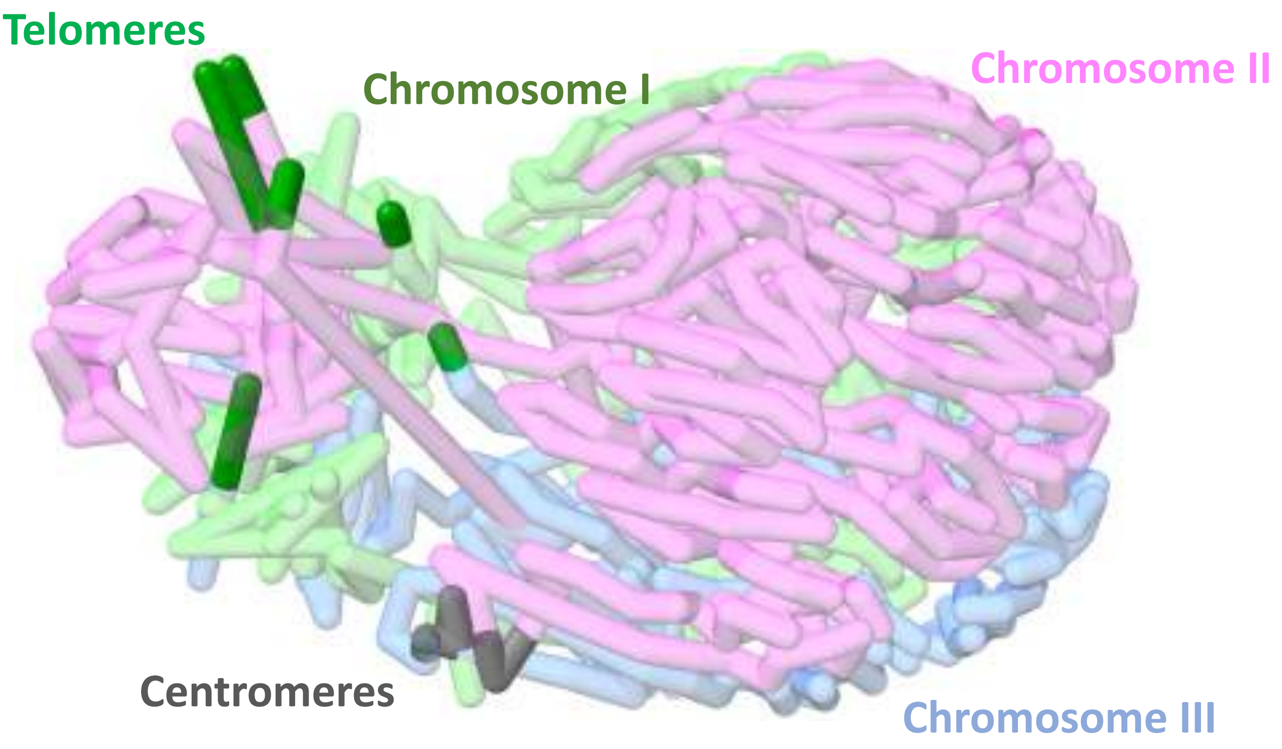

### Figure S13

Evolution of *B. divergens* and B. *MO1*

## A

Tree comparison with other *Babesia* spp (18S rRNA)

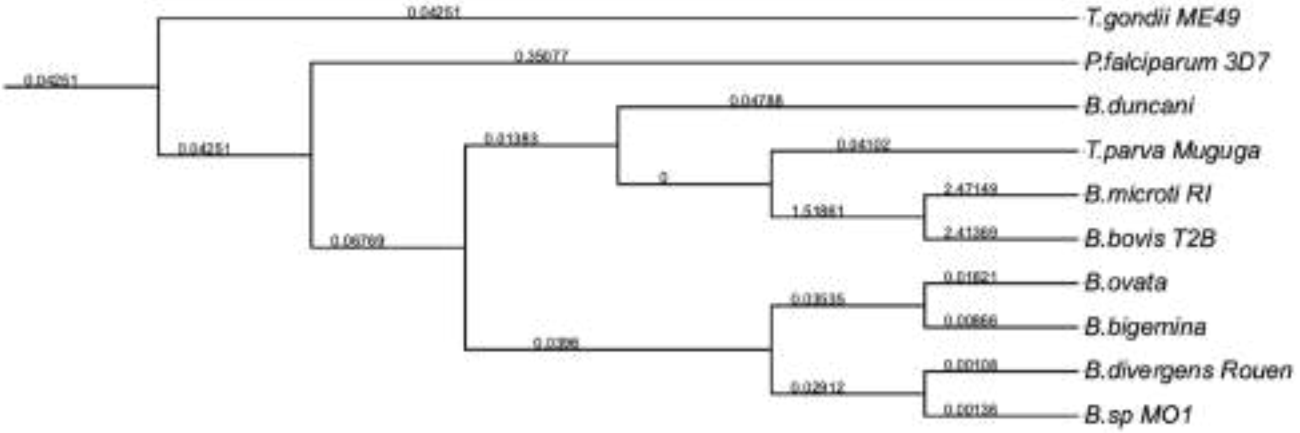


## B

Tree comparison with other *Babesia* spp (mitochondria)

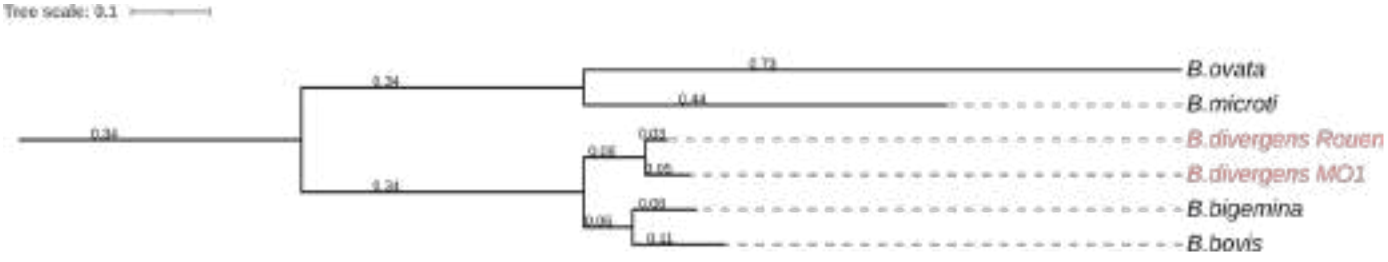
